# Localis-REX: a general protocol for time-resolved spatiofunctional profiling in live cells and *C. elegans*

**DOI:** 10.64898/2026.10.21.760500

**Authors:** Jinmin Liu, Dalu Chang, Marcus J. C. Long, Yimon Aye

## Abstract

We describe Localis-REX, a method for identifying organ- or subcellular locale-specific innate reactive electrophilic metabolite (REM)-responsive proteins in live *Caenorhabdtis elegans* and cultured cells. Localis-REX mimics the natural spatiotemporal production of REMs by expressing HaloTag transiently or stably in a specific organ or tissue of live worms or in a designated organelle or subcellular compartment in cultured cells. Following treatment with a sub-micromolar photocaged-REM and washout, light-triggered uncaging generates brief, localized REM pulses near Halo. REM-responsive proteins within the vicinity compete to capture the liberated REM before endogenous clearance processes, including glutathione adduction and redox metabolism, enabling selective identification of the most REM-responsive targets under minimally-perturned conditions. Localis-REX can be coupled with: (i) proteomics to map chemically-actionable local proteomes; or (ii) live cell and *in vivo* functional readouts to define biological responses to locale-specific REM-accumulation, such as translation-efficiency monitoring through genetic code expansion. The workflow requires 3–6 h, with downstream data acquisition and analysis requiring an additional 24–48 h, depending on the application (i) or (ii). We further describe integration of *in vivo* Localis-REX with ultraID spatial profiling in live *C. elegans*, enabling expression normalization across profiled locales. Mechanistic validation of newly-identified responder—REM-pairs is facilitated using established T-REX/Z-REX target-engagement and *on-target* REM-signaling interrogation protocols. Notably, the workflow was initally established as part of a master’s thesis project by a student co-author with no prior experience in *C. elegans* or cell-culture research by the master-student, highlighting its accessibility to well-trained junior researchers pursuing interdisciplinary studies.

**Key points:**

- Localis-REX is a technique for spatial and functional profiling of reactive metabolite-responsive proteins in nematode worms and mammalian cells. It includes steps for creation and maintainence of transgenic nematodes, performing live cell/worm-based spatiofunctional proteomics, and validation of reactive metabolite-driven protein modifications and their downstream cellular and behavioral effects.
- Localis-REX overcomes a key limitation in existing methods by enabling direct analysis of the spatial functions of reactive metabolite-responsive proteins, and the resulting modification-dependent signaling changes.

**3 Key references demonstrating the development and use of the protocol**

- Chang, D. et al. NCBP1 stress signaling drives alternative S6K1 splicing inhibiting translation. *Nat. Chem. Biol.* **2026** Feb 10. Online ahead of print. doi: 10.1038/s41589-025-02135-4.
- Liu, J. et al. Organ-specific electrophile responsivity mapping in live *C. elegans*. *Cell* **2024** *187* 7450. doi: 10.1016/j.cell.2024.10.014.
- Zhao, Y. et al. Function-guided proximity mapping unveils electrophilic-metabolite sensing by proteins not present in their canonical locales. *Proc. Natl. Acad. Sci. U.S.A.* **2022** *119* e2120687119. doi: 10.1073/pnas.2120687119.

## INTRODUCTION

Reactive metabolites (REMs) and xenobiotics function predominantly by interaction with defined REM-responsive proteins, which modulate their function upon covalent engagement with specific REMs^1, 2^. For many classes of REM (e.g., lipid-derived electrophiles, or mitochondria TCA cycle-derived REMs), target protein modification often occurs through reaction with a specific residue, typically cysteines^1, 3–5^. Identifying these sentinel proteins and their reactive residue(s) is thus crucial for understanding responses to endogenous REMs and related covalent drugs^6–10^. However, identifying “the best” sensors to a specific REM has proven particularly difficult. This is at least in part because functional responsivity of a specific protein to a specific REM is strongly context-dependent: our latest work documents examples of proteins only sensing and responding to REMs in specific organs^11^, or in one subcellular locale over another^12, 13^. Tools to identify endogenous protein responders to specific REMs and related broad reactivity covalent drugs such as Tecfidera® in defined living systems remain limited^11, 14^.

### Development of the Protocol

Methods to identify protein targets of specific small molecules are relatively common place. Classically, a large proportion of these use reactive molecule overload strategies and score protein labeling based on enrichment of labeled proteins, versus a control sample. Given that most reactive molecules react inherently with specific individual amino-acid residues [such as lipid-derived electrophiles (LDEs) with cysteine], it is unsurprising that the number of proteins that interact with reactive molecules is typically high. Most of these proteins are likely by-stander proteins, many of which are highly abundant or particularly stable under the reaction conditions. ABPP, particularly isoTOP-ABPP^15^ overcame a large number of these issues by providing a protocol that is quantitative and can rank protein reactivity based on achieving high occupancy. This method and its numerous incarnations have provided significant insights into protein reactivity, drug targets (including binding site-ID), and druggability^16^.

Other proteomics methods such as drug affinity responsive target stability (DARTs)^17^ and thermal shift assays and variants (TSAs/DSF/CETSA)^18, 19^ are also useful and can be applied on a “global” scale. Certainly TSAs, but also likely DARTs, are not necessarily apposite for profiling of covalent protein modifications. This is because the binding energy for a covalent modification by reactive electrophilic molecules (or ‘electrophiles’) is often almost entirely due to the formation of a covalent bond. It stands to reason that protein stability, and sensitivity to proteases may not be changed upon covalent ligand binding.

ABPP, DARTs and CETSA typically deploy native small molecules. However chemical modification of the electrophile, for instance, by affinity labels, and photocross linking are also possible. These methods are in principle applicable to all scenarios, although they tend to be effective only in specific instances. These approaches tend to require custom chemical synthesis and hence mandate making changes to the electrophile’s chemical structure. In many instances conservative changes can be made. However, it is important that the specific molecules in question be assessed for bioactivity prior to deployment as affinity probes.

In addition to these methods to defining chemical reactivity and chemical ligand—protein interactions, spatial profiling has become a go-to approach in biology. This typically deploys genetic methods to localize a specific (engineered) enzyme capable of creating highly reactive, short-lived reactive chemical species in a particular subcellular locale or organ. In this way, identification of proteins tagged by the specific reactive species informs on local proteome makeup. In this paradigm, reactivity of the short-lived chemical species, a parameter that defines spatial resolution, and speed of its generation, a parameter that defines temporal resolution, are crucial. Using these two parameters, APEX2 is probably the best profiling method, as it has high spatial, and uniquely high temporal resolution^20^. However, APEX2 mandates the use of toxic peroxide in high dose to activate short-lived chemical species generation, and hence other methods, such as ultraID^21^ that do not require peroxide addition, have become preferred. Newer methods such as µMap have now entered the literature^22^. These use metal complexes to sensitize diazarenes to form reactive carbenes, whose diffusion distance is particularly low, thus offering very high spatial resolution; however, this aspect may come at the cost of losing some interactions in large complexes due to the spatial resolution being too high.

Overall, it is therefore quite clear that methods with high resolution in terms of defining chemical reactivity or spatial distribution are well known. What is really missing from the canon is a method to unify these two paradigms. Unfortunately, most quantitative ABPP methods are designed to work on bulk proteomes, using absolute labeling percentage as a metric for electrophile sensitivity. Spatial profiling methods seek to do the opposite and deploy way too reactive probes to indiscriminately index local proteomes. To address these shortcomings, we developed Localis-REX, which effectively combines the benefits of ABPP with proximity profiling^23^. In this way Localis-REX constitutes one of the first protocols to achieve both spatial and functional (termed ‘spatiofunctional’ hereafter) profiling. The ability to examine these two key parameters at the same experiment strongly aids resolution, minimally by reducing variance across different samples.

#### REX technologies

Our previous published protocols have reported how reacitve molecule signaling *occurring on a specific responder protein of interest (POI)* can be interrogated in live cells (<u>T-REX</u>^24^) and zebrafish (<u>Z-REX</u>^25–27^). This suite of methods—part of “REX technologies^28^”—uses expression of a Halo-tag-fused POI, wherein Halo serves as an anchor to localize a cell/animal-permeable and bioinert photocaged-REM, which can be delivered from outside the cell/animal^27, 29, 30^. Unfortunately, despite their utility to understand protein-specific signaling events, the above protocols are not ideal for identifying new REM-responder proteins. Here we disclose a strongly complementary protocol “Localis-REX”, to quantitatively map organ-specific^11^ or subcellular locale-specific^12, 13^ REM-responders in live *C. elegans* and cells, via controlled, localized intracellular generation of specific REMs. In Localis-REX, the expressed Halo is not fused to any POI, but its organ-specific expression is driven by native tissue-specific promoters (in transgenic animals), or subcellular/organelle-restricted expression, by genetically encoding Halo with the corresponding localization sequence (**Fig. 1a**). Light-driven photouncaging allows rapid build-up of a defined REM within the vicinity of Halo. As the REM is available transiently in limited dosage, in an otherwise unperturbed background, only the best responders to a given REM are mapped.

**Figure 1.**
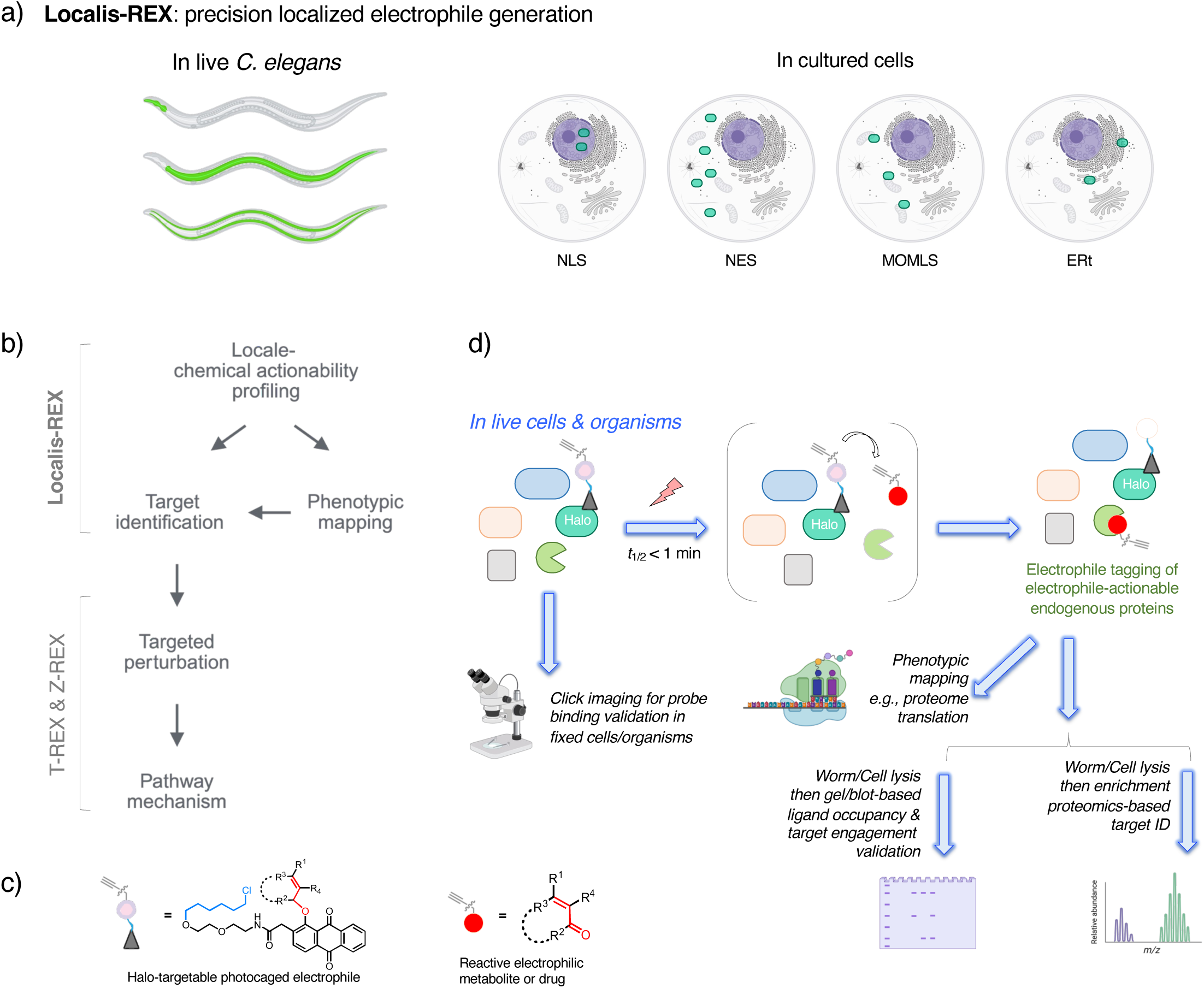
The Localis-REX approach and its two experimental applications. **(a)** This protocol demonstrates Localis-REX locale-specific spatiofunctional profiling workflow in **Left:** live *C. elegans* using tissue/organ-specific native promoters (e.g., *myo-2p* for pharyngeal muscle, *ges-1p* for the intestine, and *myo-3p* for body-wall muscle) driving locale-specific GFP-Halo expression, and **right:** live mammalian cells using expression vectors encoding Halo fused to subcellular localization sequence (e.g., NLS, NES, MOMLS, and ERt, respectively for nucleus, cytoplasmic, mitochondrial outer membrane, and endoplasmic reticulum-specific Halo expression). Green icons indicate Halo. These setups allow controlled localized buildup of reactive metabolites (REMs), offering enhanced control over timing, locale, REM-dosage, and -chemotype. **(b)** This protocol applies Localis-REX in two powerful ways: **top:** REM-responder identification coupled to mass-spectrometry-based target-ID; and **bottom:** mapping changes in downstream phenotypes (e.g., proteome translation efficiency) in response to precision upstream perturbation. Localis-REX as an upstream discovery pipeline can be coupled with our previously published protocols, T-REX^24^ and Z-REX^25–27^, to interrogate functional consequences of REM signaling triggered selectively by a specific REM-responder protein engagement with a specific REM. **(c)** In the Localis-REX workflow, live cells or animals are treated with cell/animal-permeable and biocompatible photocaged-REM probe tethered to a Halo-targetable hexyl chloride unit. **(d)** Following washout, light (5 mW/cm^2^, 365 nm, 5 min) exposure rapidly releases the REM or a related electrophilic drug (*t*_1/2_^photouncaging^ ∼ 0.5 min). Native responders within the vicinity of Halo compete for limited quantity (nm-µM) of REM made available transiently [prior to endogenous detoxification mechanisms, including reaction with glutathione (mM)]. The most kinetically-privileged responders are covalently REM-adducted rapidly (**top**). By integrating alkyne-motif non-invasively in the released REM, these responders can be analyzed by SDS-PAGE and western blot, or identified by enrichment-based digest LC-MS/MS (**bottom right**). Binding of the photocaged-REM to localized Halo in fixed cells/animals can also be validated by immunofluorescence imaging, post probe treatment and washout but prior to light-driven REM generation (**bottom left**).

Localis-REX has several unique features. No other method has the ability to generate endogenous or related drug-like electrophilic fragments with user control over timing, dosage, locale, and REM-chemotype in living systems. Breaking these concepts down, Localis-REX has particularly high temporal resolution (1 min). This is on par with or shorter than some of the most rapidly changing processes known in biology, for instance, synthesis of a 50 kDa protein (30 s)^31^, important steps during T-cell receptor signaling (5-10 minutes)^32^, and duration of specific phases in the cell cycle (30 min to several hours)^33^. Most competitive methods deploy much longer reaction times, at least in part because they cannot generate specific reactive entities in a cell. REX-technologies also give control of dosage (maximally, low micromolar in the total cell^10^). One of the largest variables when studying reactive small-molecule function in cells is permeation, stability, and distribution. Using peroxide as a quintessential reactive small signaling molecule whose diffusion in cells can be readily visualized, when peroxide is administered from the outside of a cell, a large gradient is established across the cellular membrane^34^. For this reason, peroxide is rarely able to penetrate into the nucleus (unless redox homeostasis is severely perturbed)^35^. Thus, the concentration of peroxide, and similarly, other reactive small molecules, is highly contingent on detoxification, time of exposure, and other locale- and context-specific cellular processes, including distance from the cell membrane. Localis-REX standardizes all these parameters by maintaining short exposure times, releasing from inside the cell, and allowing shifting of point of origin of release. The last point underscores the spatial control of REX-technologies, where the point of origin of the reactive signal can be controlled spatially (e.g., organelle). Finally, because many experiments using reactive small molecules are carried out on the duration of minutes to hours, the reactive metabolite under study could be modified metabolically, for instance by detoxification mechanisms. Again, by limiting times of exposure to the specific chemical signal, Localis-REX improves chemotype resolution of typical strategies. All these elements remain state of the art.

### Overview of the procedure

We here describe how to generate and validate relevant biological systems to study locale-specific reactive electrophilic metabolite (REM)-signaling. Applications covered in the protocol include the nematode *C. elegans* expressing Halo in specific regions/organs under organ-specific (OS) promoters, and cultured mammalian cells expressing Halo in specific compartments. Throughout this protocol, the REM delivered is 4-hydroxynonenal (HNE), released on demand from the HaloTag-bound photocaged precursor Ht-PreHNE. Adaptation to other REMs can be achieved using the previously described photocaged precursors to other lipid-derived electrophiles used in T-REX^24^ and Z-REX^25–27^ protocols. Localis-REX answers two distinct questions in living subjects:

i. what proteins are chemically-actionable in a given locale?;
ii. to what extent controlled REM-stress in a given subcellular locale induces functional changes in cellular responses.

For point (i), we describe how to perform REM-responsivity mapping in the two distinct systems. The dosage and treatment time of the Ht-PreHNE(alkyne) probe are first optimized for each Halo-transgenic nematode strain by a TMR-blocking titration, read out by in-gel fluorescence, in which TMR-ligand labels the HaloTag molecules that had not been probe-bound following Ht-PreHNE(alkyne) treatment in live worms (Steps 32-51). Localis-REX workflow comprises probe treatment, washout of unbound probe, photouncaging, and cell/animal lysis (Steps 55-60 for *C. elegans*; Steps 103-117 for cultured cells). Alkyne-tagged adducts are then conjugated to biotin-azide by copper-catalyzed click coupling and captured on streptavidin (Steps 77-95). (Should a more rapid whole-lysate-based readout be preferred, Cy5-azide is substituted for biotin-azide and the sample is analyzed by in-gel fluorescence, without enrichment of the HNEylated proteins). Enriched fraction is submitted for tryptic digest coupled quantitative mass spectrometry using using label-free quantification (LFQ)^36, 37^, stable isotope labeling by amino acids in cell culture (SILAC)^38^, or tandem mass tagging (TMT)^39–41^ strategies (Steps 153-169). We discuss the relative merits of these different methods of quantitation and hit selection and validation criteria in Experimental Design. The three negative controls required at each stage, namely, the use of Ht-PreHNE(no alkyne), no light exposure, and no Halo-transgene-harboring strains, are set out in Box 1 and cited from the steps at which they must be established.

We also present a strategy to perform organ-specific ultraID^21^ to profile local proteome abundance in specific organs examined in *C. elegans* (Box 3). ultraID is a compact, hyperactive biotin ligase that biotinylates nearby proteins, reporting their local presence and abundance, but not their function or responsiveness to specific biologically-relevant small-molecule ligands. ultraID datasets provide an orthogonal, abundance-based map of the same locales, against which REM responsivity can be assessed. Low overlap is informative demonstrating that Localis-REX hits are not biased by local protein abundance.

We additionally discuss streamlined mechanistic interrogations of top candidate hits in two directions: at the protein-specific level, as well as phenotypic level of responses, including using systems with endogenous REM-stress (**Fig. 1b**). At the protein-specific level, individual candidates are interrogated by T-REX^24^ and by mutagenesis of the implicated cysteine. At the organismal level, candidates are knocked down by RNAi feeding in *C. elegans* and scored for survival, motility, reproductive output, and the defecation motor program, with Oil Red O staining used to read out neutral-lipid content following electrophilic challenge (Steps 169-182). These phenotypic assays deliver a bolus dose of HNE(alkyne) rather than the photocaged probe, because these experiments ask whether a responder is required for the organism’s response to REM stress rather than identifying targets.

For point (ii), we demonstrate how to setup live-cell-based Localis-REX in tandem with established live-cell-based genetic code expansion (GCE) or BONCAT nascent protein translation readouts (Steps 137-142 and 143-152 respectively. GCE reports synthesis of a single nascent reporter protein; BONCAT that of the nascent proteome). This approach reports on changes in cellular translation efficiency in response to compartment-specific REM build-up (**Fig. 1c**).

The Procedure additionally covers the supporting stages required to reach these experiments: *C. elegans* culture, synchronization and stock preparation (Steps 1-15); male induction, genetic crossing and single-worm PCR genotyping, performed only when an organ-specific driver must be combined with a second genetic background (Steps 16-27); TMR-Halo staining to confirm that Halo is expressed in the expected organ and accessible to ligand in live animals (Steps 173-176); and immunofluorescence validation of probe localization in cultured cells (Steps 119-135).

### Applications

Precision REM-sensing and -signaling programs are crucial steps in various endogenous REM-regulatory pathways, including immune cell function^42, 43^, ferroptosis^44–47^ and mitochondrial signaling^48, 49^, and so on. They also impinge on reactive electrophilic drug target engagement, and can inform on novel drug design^50, 51^. The Localis-REX protocol is readily applicable to—and variants are compatible with— cells/tissues undergoing different signaling processes, including apoptosis, transcription, and differentiation. We thus envision a very broad interest and a large number of potential applications. The success and broad deployment of activity-based protein profiling (ABPP)^52^ methods that assay reactivity changes in a subset of proteins within the proteome, provide further strong support for these predictions. One benefit of REX-tools over ABPP-based approaches, is that with subtle changes to the protocol, REX-technologies offer a validated means to assay the importance of REM/REM-responder-pair engagement directly in intact living systems. Hits from other profiling strategies, such as ABPP, can be evaluated using REX-technologies. Thus, this latest tool, Localis-REX, in live cells and animals, alongside previous deployment of <u>T-REX</u>^24^ and <u>Z-REX</u>^25–27^ technologies further extend the remit of our work to several communities focusing on model organisms, including neurodegeneration, inflammatory signaling, and immunology.

### Comparison with other methods

Localis-REX, however, does not directly afford information provided by some well-known techniques. Localis-REX seeks to identify specific REM-responders. It does not therefore identify specific REM-responsive residues. Many ABPP-based methods can directly afford this information^15, 53^. DARTs and CETSA do not. Residue-specific information is important for designing mutants and potentially for drug development. However, it is well known that several REM-responders possess two or more REM-sensor residues; moreover, many potential REM-sensor cysteines within proteins are not commonly detected by classical ABPP-based screening approaches because they are located on peptides that have low proteotypicity, i.e. probability of being detected by MS^16^. Residue resolution can therefore come at the cost of target identification.

Unlike many quantitative proteomics profiling methods, Localis-REX also does not afford quantitative insights into percentage of protein modified; it only pinpoints REM-responders significantly differentially enriched from bulk proteomes. In practice, this is a consequence of the aims of Localis-REX, which seek to identify specific responders functionally responsive in specific biological locales. Captured responders may well exist in other locales (a different subcellular compartment, cell type, etc.) but these would not be REM-modified. This explains why traditional ABPP-methods could not identify such locale-specific responders as electrophile sensors in the first place. Nonetheless, this scoring strategy can be restrictive, particularly in model organisms, such as *C. elegans*, where the number of hit REM-responders remains low, as the method presently relies upon stoichiometric release of one REM molecule at maximum, per Halo.

In terms of spatial resolution, as discussed above, the rate of deactivation of a reactive chemical species defines the half-life of that particular species. ultraID, APEX2, and µMap all use much more reactive chemical species than Localis-REX. This indicates that spatial resolution of these profiling methods are likely higher than REX technologies. For instance, the intrinsic diffusion distance of a singlet carbene, deployed in Micromap, is in the nanometer range, the diameter of a 50 kDa globular protein^54, 55^. In contrast, the half-life of 4-hydroxynonenal (HNE), a well-studied lipid-derived REM which we have used in localis-REX, in close-to-physiological concentrations of GSH (5 mM), is 2 min, giving a diffusion distance of around 1 mm^4^. Glutathione (GSH) is the dominant cellular small-molecule nucleophile, which enables enzymatic detoxification and non-enzymatic clearance of electrophiles like HNE. This is far beyond the length of a cell. However, intracellular enzymatic detoxification of HNE (e.g., by glutathione transferases, and redox enzymes) is significantly faster than the non-enzymatic bimolecular reaction of HNE and GSH (10^4^ vs. 1 M^-1^s^-1^, respectively), likely restricting HNE’s diffusion distance.

Indeed, we have evidence for enzymatic restriction of HNE’s diffusion distance, as we have successfully profiled HNE-responsive proteins in the cytosol over the nucleus, and vice versa^12^. This implies that HNE’s diffusion distance in a cell is on the order of microns or less. This distance is on the order of many different subcellular structures, such as mitochondria, ER, and the nucleolus—an attribute that may render REX-technologies apposite for profiling such structures. This property also likely makes Localis-REX very effective at whole-cell profiling. However, it remains to be seen if Localis-REX can profile more spatially-restricted locales, where Micromap/APEX2/ultraID have already proven their worth. Ultimately, it is important to bear in mind that Localis-REX using HNE has very context-dependent spatial resolution, as we rely on detoxification processes to engender high spatial resolution. Other spatial profiling methods that deploy non-native reactive species are much less dependent on these processes.

Other key differences between Localis-REX and spatial profiling methods surround the fact that spatial profiling methods aim to achieve saturation of specific proteins in a specific locale. This is achieved at least in part through catalytic deployment of highly reactive species that engage indiscriminately with proteins. Localis-REX on the other hand releases a transient burst of moderately reactive molecules. This is because it seeks only to identify proteins with a specific REM-responsivity in a locale. In principle, REM-responders identified by Localis-REX should be a subset of those identified by other spatial profiling methods. However, spatial profiling methods certainly gravitate towards highly abundant, stable, and potentially large proteins, that can be labelled excessively. REX-methods by virtue of the low reactivity REMs deployed and them identifying kinetically reactive proteins, do not. Accordingly, spatial profiling and REX methods are particularly complementary and it is for this reason that we will deploy them in tandem in certain circumstances as described in this protocol where we perform organ-specific (OS) Localis-REX in parallel to OS ultraID, both in live *C. elegans*.

### EXPERIMENTAL DESIGN

#### General considerations

In this work, Localis-REX in *C. elegans* and in mammalian cells is described as two representative examples. Considerations involve the strength of Halo-expression and specific REM(s) to be profiled [a range of different photocaged-REM probes are available to REX-technologies^24^, and the same probes are used for Localis-REX and T(Z)-REX]. HNE is used here as a representative reactive electrophilic metabolite (REM), and the workflows described are specific to it.

##### General considerations for *C. elegans* Localis-REX

Success of Localis-REX is critically dependent on the selection of tissue-specific promoters or subcellular localization sequences to drive Halo expression. In *C. elegans*, many promoters drive target gene expression in specific organs/cell types. These may also show expression in other regions to a lesser extent. Expression regions may also change across development stages. Detailed information about expression patterns can be obtained from Wormbase, published single-cell RNA sequencing data, or by perusal of published work on the promoter of interest^56, 57^. The chosen promoter is recommended to confer precise tissue-specific expression, as off-target expression in other regions will introduce noise in the subsequent mass spectrometry analysis. In this paper, *myo-2* promoter for pharyngeal muscle expression^58^, *ges-1* promoter for intestinal expression^59^, and *myo-3* promoter for body-wall muscle expression, were chosen. Our constructs were designed with a fluorescent reporter to validate expression in specific tissues (**Fig. 2a**). These systems give relatively similar levels of Halo expression per worm. It is noteworthy that certain promoters drive expression only in a few cells. For example, the *gpa-4* promoter drives protein expression in two cells (ASIL and ASIR) in adult worms^60^. When performing Localis-REX experiment with such strains, the sample amount needs to be increased to obtain a sufficient quantity of enriched proteins for mass spectrometry analysis. Thus basic biochemistry, assignment of expression levels, and appropriate controls and comparisons must be considered in the planning stages (for instance, if using worms with large Halo expression level differences, it may be prudent to include wild-type worms as a global background comparison).

**Figure 2.**
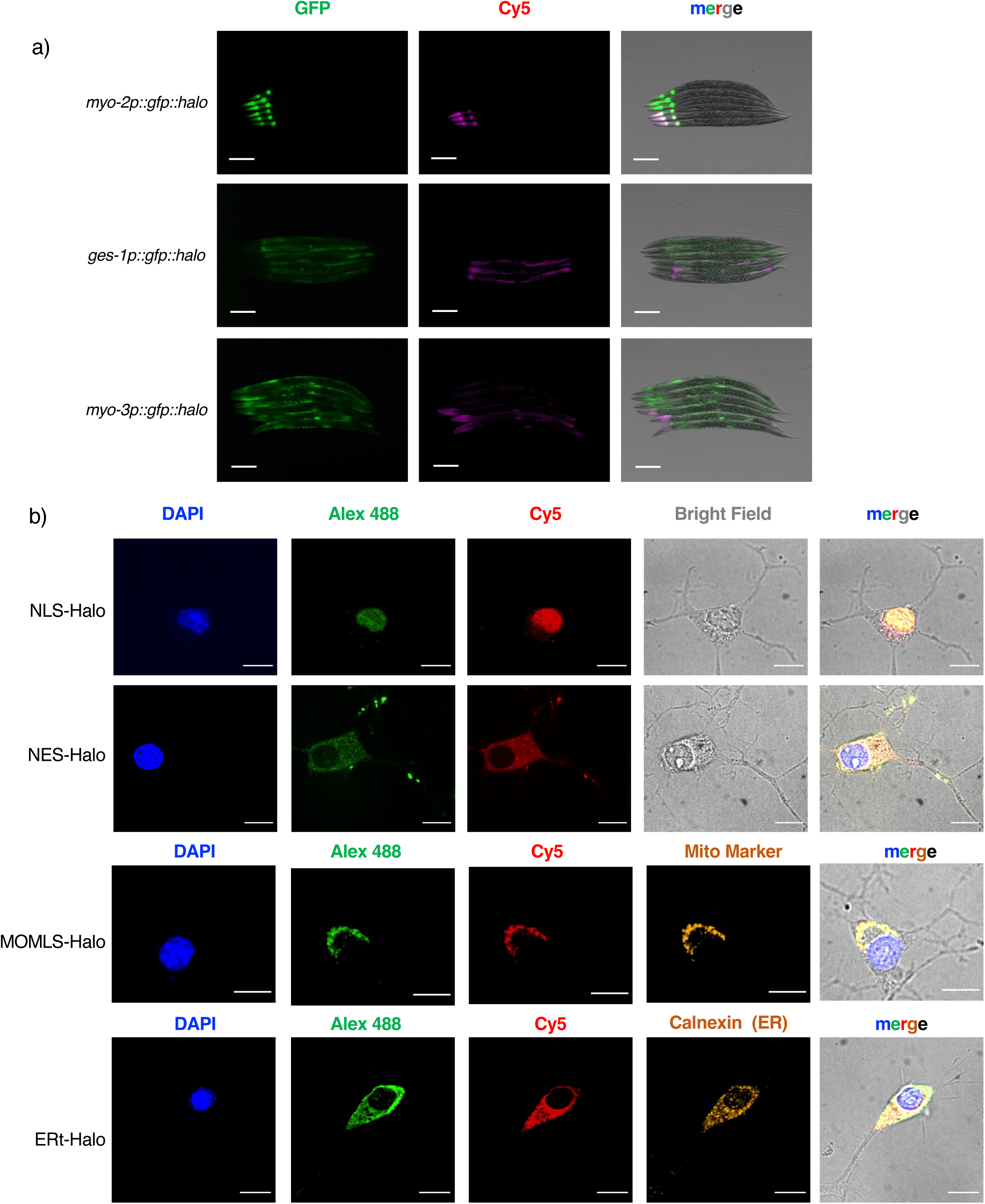
Validation of locale-specific Halo expression and photocaged-probe binding. **(a)** Locale-specific Halo expression and photocaged-REM probe binding were validated by immunofluorescence and Click-imaging in fixed worms. *myo-2p* drives GFP-Halo expression in pharyngeal muscle, *ges-1p* for the intestine, and *myo-3p* for body-wall muscle. The Cy5 channel represents the bound probe coupled with azide-Cy5 following Click reaction. Each image shows 3 respective age-synchronized organ-specific GFP-Halo-expressing hermaphrodites at day 1 adult stage with alkyne-coupled probe treatment (3 lower worms), against 3 with no alkyne-coupled probe treatment (3 upper worms). Scale bars: 200 μm. This figure is adapted from previously published data in reference^11^. **(b)** Immunofluorescence imaging validates expected localization of Ht-PreHNE(alkyne) covalently bound to Halo expressed at designated subcellular locales in fixed cells. Differentiated N2A cells ectopically expressing NLS-, NES-, MOMLS- or ERt-Halo were treated with Ht-PreHNE(alkyne), washed, fixed and permeabilized. Probe-bound Halo was revealed by Click coupling to Cy5-azide, and the Halo was detected in parallel by immunostaining. DAPI and MitoTracker Red were used as a nuclear and mitochondrial marker, respectively. ER-resident Calnexin was used as an ER marker. Conditions without Cu (hence, no Click coupling) were used as control for background labeling with Cy5-azide. Imaging was performed using a Nikon spinning disk confocal microscope equipped with a 100x objective. Scale bars: 10 µm. This figure is adapted from previously published data in reference^13^.

One begins *C. elegans* Localis-REX experiments by choosing locales of interest, and potentially genetic backgrounds or conditions, in which to perform spatiofunctional profiling. Researchers could also use Localis-REX to address how locale-specific REM-responsiveness alters as a function of age, stress levels, or the extent of proteostasis imbalance. Once precise test conditions are determined, controls should be assigned. Comparison between different locales/conditions under which Localis-REX has been undertaken is in principle, sufficient. However, it is generally a good idea to have at least one basal control (for instance, untreated, or no-alkyne-containing probe) to serve as a “true” background (**Fig. 1c**, **Fig. 3a**), especially in *C. elegans*, where endogenous biotinylation is common. For some conditions, even more controls could be needed, such as stressed cells with no probe, or different ages. These variables all have to be considered in the planning, factoring in number of samples intended to be processed (cost, statistical analysis), comparison groups, etc. One important aspect that is particularly important to bear in mind when planning Localis-REX experiments is that T-REX^24, 25^ allows a very simple, but particularly powerful means to investigate and quantitatively rank protein-specific REM-responsivity^24, 28^. This is an essential step, which is the gold standard of validation. Providing these downstream validations are factored into the planning, it can be argued that exhaustive controls during the screening stage are not necessary (**Fig. 1b**).

**Figure 3.**
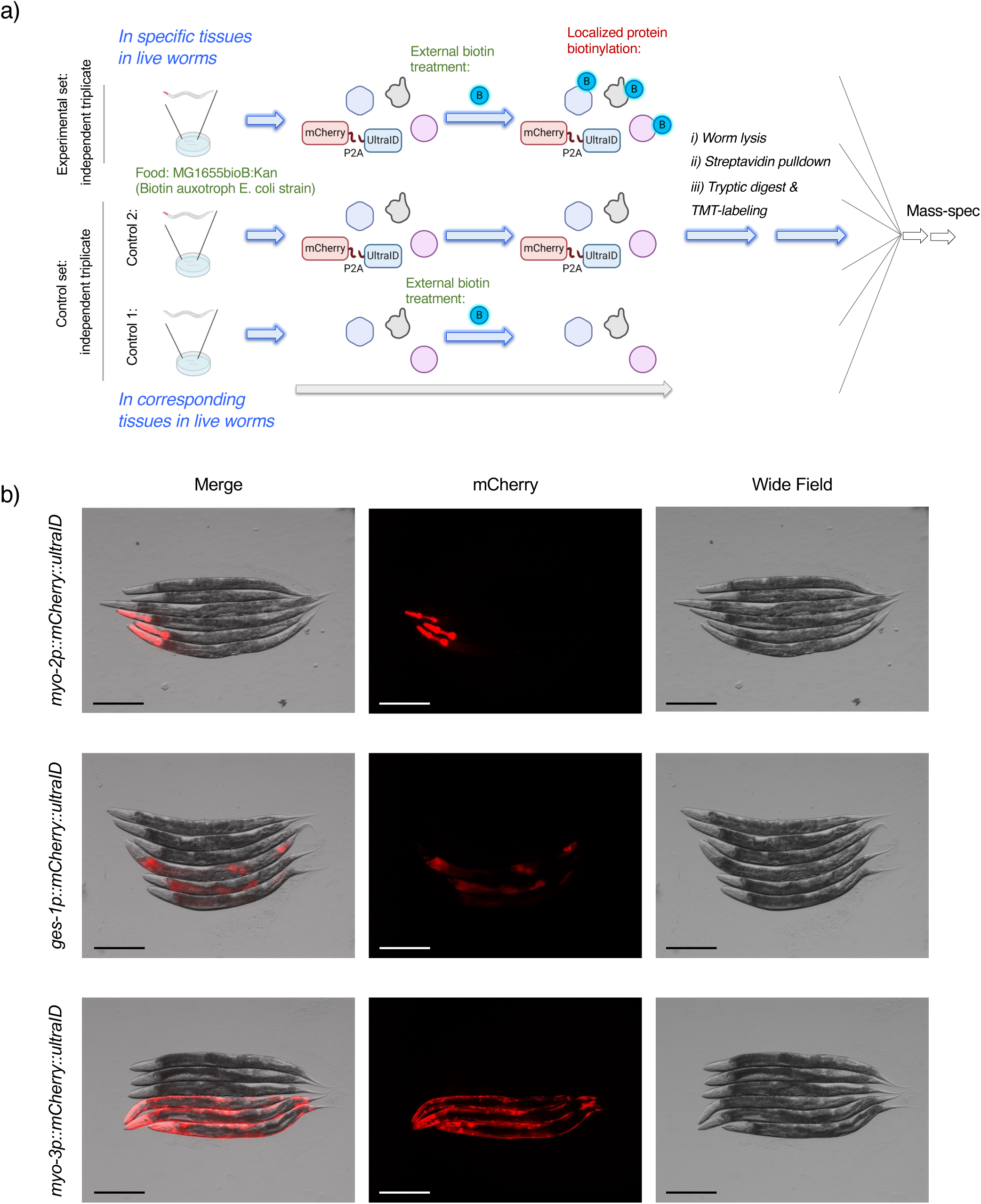
Organ-specific local proteome abundance mapping by OS-ultraID. **(a)** *C. elegans* organ-specific local proteome abundance mapping (OS-ultraID) workflow in live worms. Due to the higher level of endogenously biotinylated proteins in *C. elegans*, two sets of controls [no ultraID expressing strains (control 1), and no external biotin treatment (control 2)] were included. All groups were fed with *E. coli* MG1655bioB:Kan. For LC-MS/MS identification of biotinylated local proteomes, animals were lysed, followed by streptavidin pulldown, trypsin digestion, and subjected to TMT labeling. This figure is reproduced from previously published data in reference^11^. **(b)** OS-ultraID was validated by organ-specific expression of ultraID enzymes using live imaging. Worms expressing mCherry-P2A-FLAG-ultraID (at 3 specific tissues identical to Halo expression in OS-Localis-REX strains: *myo-2p* for pharyngeal muscle, *ges-1p* for the intestine, and *myo-3p* for body-wall muscle) were immobilized with levamisole. Each image shows 3 respective age-synchronized transgenic hermaphrodites at day 1 adult stage (lower) against 3 age-matched wild-type N2 hermaphrodites (upper). Scale bars: 200 μm. This figure is reproduced from previously published data in reference^11^.

Nonetheless, covalent modification of a protein by a reactive molecule is a second-order process. Thus protein-specific modification is dependent on concentration of REM (that we hold constant) but also concentration of each responder protein, as a function of tissue-specific expression. Thus, enrichment can happen in Localis-REX due to overexpression of a specific responder protein in one locale over another. We recommend using ultraID (or another locale-specific abundance profiling method^23, 61, 62^) to address these issues, allowing triaging of hits caused simply by mass action to be removed (**Fig. 3a**).

Regardless, well-performed Localis-REX experiments will create, based on our data, tens to hundreds of hit responders. The subsequent question is which of these hits reflect locale-specific reactivity rather than simply being caused by high local abundance. We presented a representative example to show that most of our Localis-REX hits are not biased or predicted by high local abundance (**Fig. 4**). In **Fig. 4a**, the upper red row shows the Localis-REX responder signals, whereas the lower blue row shows organ-specific transcript abundance for the same genes, derived from published single-cell RNA-seq data. These patterns are largely discordant: > 70% of significant responders (red asterisks) are not correlated with the most organ-enriched transcripts. **Fig. 4b** provides the corresponding control. OS-ultraID abundance closely tracks the same transcriptomic data, indicating that the discordance observed in **Fig. 4a** is not a workflow artefact.

**Figure 4.**
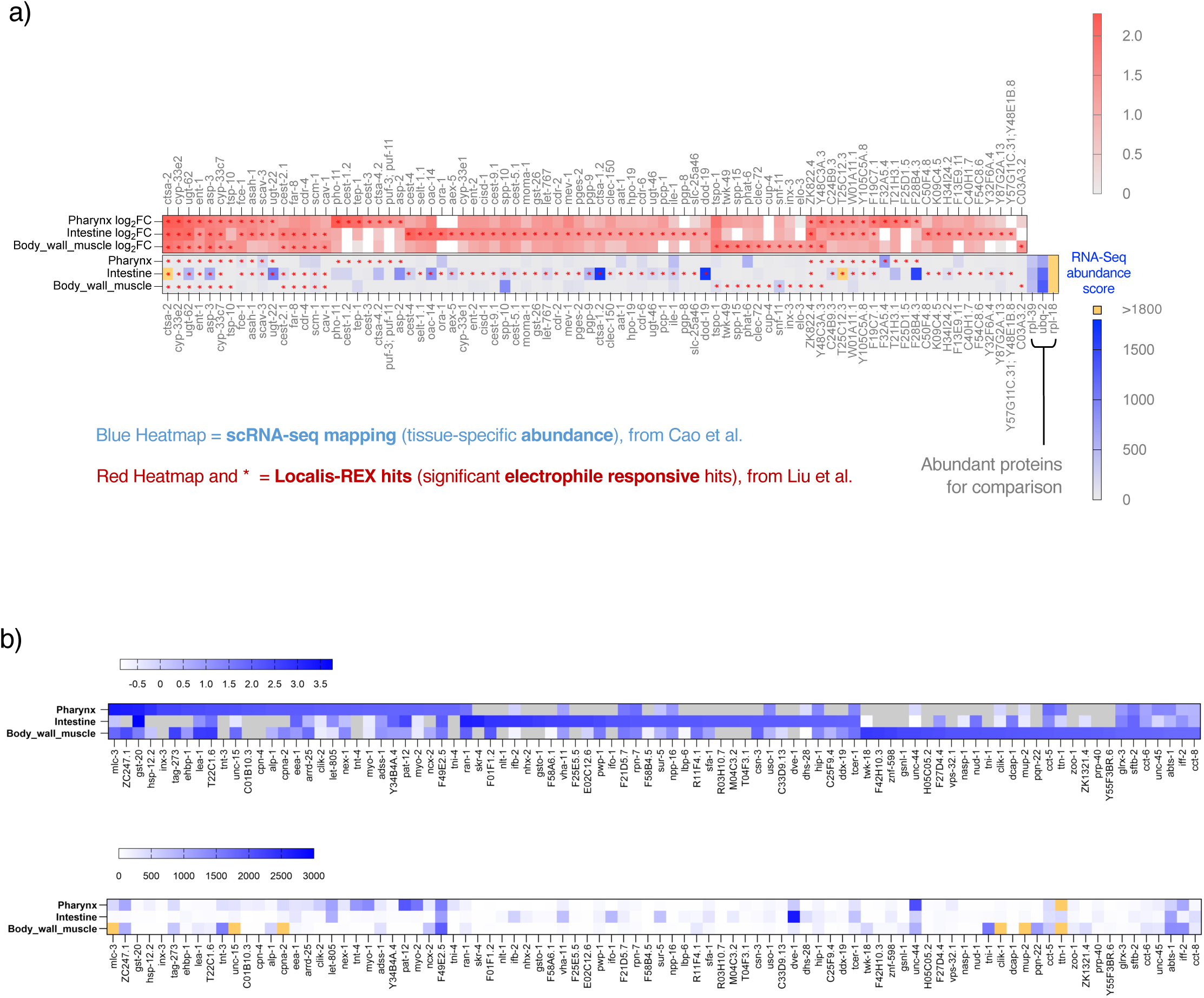
Localis-REX responders and OS-ultraID abundance compared with organ-specific transcriptome data. **(a)** Heatmaps comparing REM-responders derived from *C. elegans* Localis-REX (in red, upper row) and organ-specific abundance derived from the previously published *C. elegans* single-cell RNA-sequencing datasets (in blue, lower row) targeting at the three identical organs, pharynx, intestine, and body-wall muscles. Red asterisks correspond to statistically significant hit REM-responders mapped by Localis-REX. The comparison demonstrates that organ-specific REM responsivity is not necessarily correlated with organ-specific abundance. **NOTE:** Highly abundant *C. elegans* proteins: RPL-39, UBQ-2, and RPL-18, were not identified and did not emerge as hit REM-responders in Localis-REX. The blue heatmap is adapted from previously published data in reference^11, 56^. Data in the red heatmap were previously reported in reference^11^. **(b)** Heatmaps comparing organ-specific local proteome abundance derived from *C. elegans* OS-ultraID (upper, top 30 hits in each strain) and organ-specific abundance derived from the previously published *C. elegans* single-cell RNA-sequencing datasets (lower row) targeting at the three identical organs, pharynx, intestine, and body-wall muscles. The two datasets exhibit highly similar distribution patterns and correlation profiles, indicating that in worms RNA and protein abundance largely parallels each other. Gray boxes represent corresponding proteins not identified. Yellow boxes correspond to transcript scores >3,000. This figure is reproduced from previously published data in reference^11^.

We have also outlined how we go about triaging expression-independent hits bioinformatically and experimentally. Unless a protein does not have any orthologs in humans, it is our opinion that all proteins should be viewed on their merit and investigated accordingly. We outline below *C. elegans*-based screening methods using RNAi and other relatively high-throughput assays that can be coupled to organ-specific (OS)-Localis-REX, bolus dosing, or simply, standard nematode assays to assign importance of hits. Such screens should be chosen to reflect the organs and conditions screened in the assay.

Hit responders can be carried forward to screening in cultured mammalian cells, and/or on recombinant proteins. Conserved hit responders are not only relevant to human health, but are also likely relevant to informing on evolution of REM-signaling, an area that we are particularly interested in. It is our hope that by mapping evolutionary trajectories of specific responder protein-cysteines, we can decode responsive cysteine REM-modification (**Fig. 5**). As examples, CYP2A6, NCBP1, and UBE2V2 sequence logos illustrate this analysis: a tall, near-invariant cysteine at the aligned position supports evolutionary constraint and prioritizes the site for validation in cultured cells or recombinant protein. The output is therefore a short list of conserved responder-cysteine pairs after the site-specific validation via T-REX^24^, rather than proteins alone.

**Figure 5.**
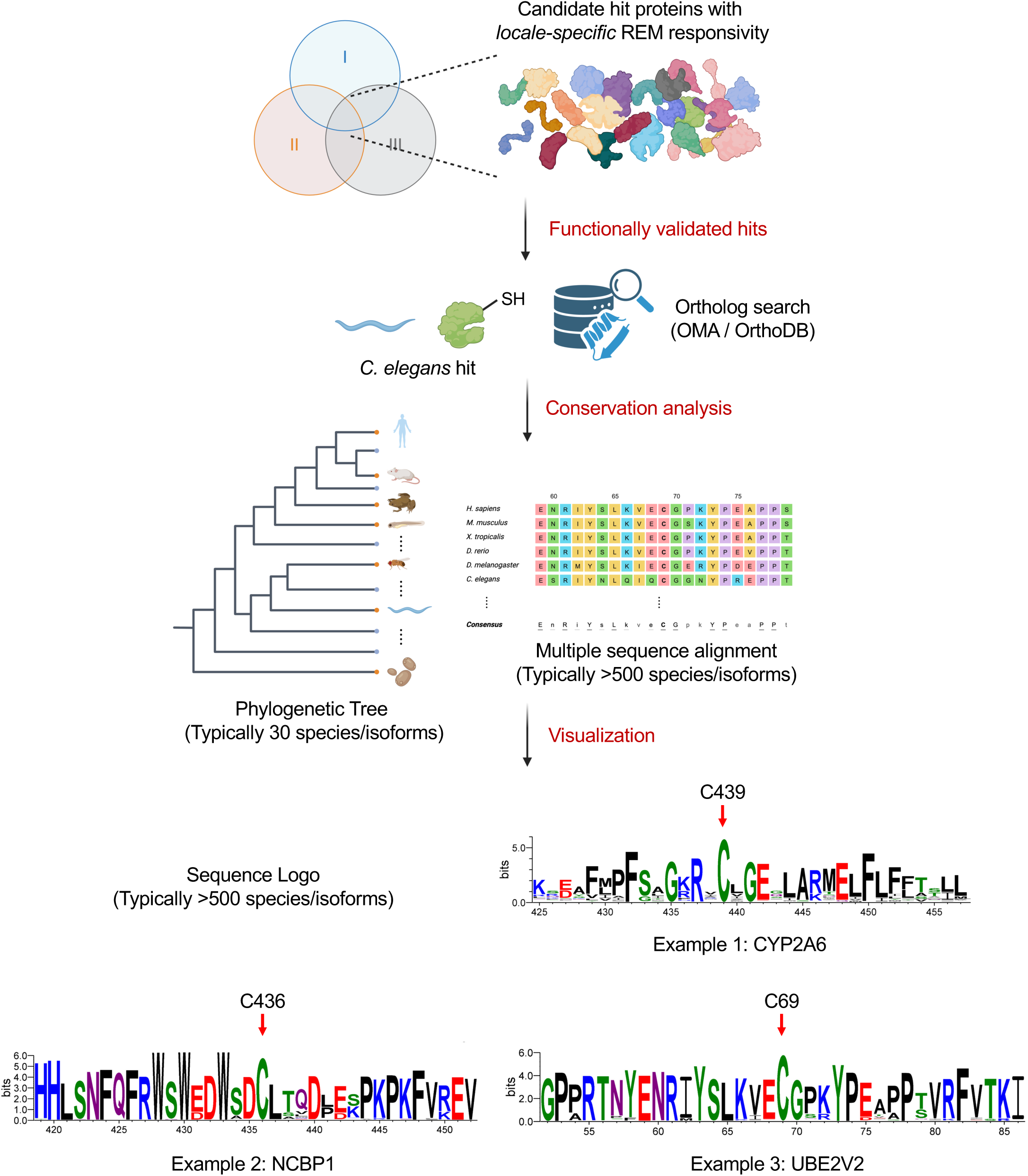
Cross-species conservation workflow for prioritizing Localis-REX responders. Schematic of cross-species conservation workflow applied to *C. elegans* tissue-specific REM-responder hits mapped by Localis-REX. Candidate hit proteins from Localis-REX coupled to enrichment proteomics-based target ID experiments, are prioritized using the target filtering and analysis workflow (Steps 165-168) described in ‘Procedure’. The Venn diagram shows the number of statistically significant REM-responder hits scored in each of three independent biological replicates and their overlap. Hits recovered in all three replicates, corresponding to the central intersection, are prioritized for the validation and conservation analyses that follow. After T-REX gel-based REM-responsivity validation^24^, REM-responder hits as scored by T-REX are subjected to ortholog identification using OrthoDB as the primary database and OMA (Orthologous Matrix) as an optional cross-check where applicable. Orthologous protein sequences are collected and aligned by multiple sequence alignment, typically using >500 species/isoforms. (Image shows partial amino acid sequence of cysteine-site-specific REM-responder UBE2V2(C69)^10^ aligned across 6 indicated species). A representative subset of aligned orthologs, typically ∼30 species/isoforms spanning major clades, is used for phylogenetic tree reconstruction. Sequence logo visualization reports conserved regions surrounding the REM-responsive protein-cysteine. Representative examples are shown for three previously validated REM-responders: CYP2A6(C439) (936 species), NCBP1(C436) (1509 species), and UBE2V2(C69) (1245 species). Red arrows denote the REM-responder cysteine position. Protein names and cysteine sites numbering refer to the human orthologs. CYP2A6 sequence logo data is reproduced from previously published image^11^.

While undertaking the above analysis, it is helpful to keep in mind that some of the most highly conserved cysteines are likely those involved in structural or functional properties of the protein. Although such cysteines have been linked to REM-sensing and signaling, in many cases, surface cysteines with no apparent structural or functional role engage in REM-sensing. These surface cysteines are not necessarily conserved across large amounts of evolutionary space and hence conservation is merely a guide to help prioritize when a large number of hits have been detected.

##### General considerations for Localis-REX in cultured cells

As above, prior to embarking on a locale-specific protein responder screen, one has to choose the locale to be profiled and the specific cell line(s) and/or conditions (**Fig. 1a**, **Fig. 2b**). This sort of planning can be based on a hypothesis or underlying biological questions. However, it can also be planned based on a screen of different locales for sensitivity to release of a specific REM-chemotype. Overall, where possible, we advocate the latter, because the workflow is feed-forward, linking phenotypic readouts to changes in a specific REM.

Once specific locales for profiling have been chosen, once again, controls and the like have to be picked. Just as for *C. elegans*, it is important to pick relevant controls, minimally the cell line where no REM, or the REM without the reactive handle (Michael acceptor motif) has been released. Again, if the intention is to compare various treatment conditions, or different cell lines, more controls will be necessary. Nonetheless, because of the power of REX-technologies, we anticipate tens to potentially hundreds of enriched hits from locale-specific Localis-REX in cultured cells. Obviously, the number of hits recorded depends on the specific thresholds applied to score as hits. For cell culture, we typically advise looking at the most statistically-enriched hits, although this ranking is not directly quantitative, and hence it may be necessary to investigate proteins with a lower overall score. In such situations, starting from areas where REM-buildup causes a phenotype (**Fig. 1c**) allows focusing on proteins whose functions are linked to the phenotypes in question. This approach may miss “unknown” interactions, although it is a means to triage hit responders.

In terms of this protocol, we discuss the use of genetic code expansion (GCE) either through amber suppression of a non-sense mutation in actin, or through whole genome labeling. Both methods work well and in our hands give very similar outputs^13^. Arguably, looking at the whole proteome is favored because it is not examining artificial mRNA or protein production. However, suppression of single proteins offers a single band that is particularly appropriate for quantitative work. On the other hand, BONCAT provides a simple procedure analogous to whole protein GCE suppression. In our hands, these readouts are highly complementary, and serve mainly as orthogonal readouts strengthening the data.

#### Plasmid design for Localis-REX in mammalian cells and *C. elegans*

For the design of expression constructs, different strategies are taken depending on the host system. For expression in mammalian cell lines, constructs typically incorporate a FLAG- or HA-epitope tag to facilitate the validation of protein expression via western blot. In our experience, the HA-tag offers higher specificity. For expression in *C. elegans*, we generally fuse a fluorescent protein engineered with synthetic introns to the N-terminus of the Halo (or ultraID). This element serves a dual purpose: the presence of introns within GFP enhances expression; mature GFP provides a visual reporter to confirm stable and specific localization. A P2A self-cleaving peptide sequence may be inserted between the fluorescent protein and the Halo/ultraID. Spatial restriction is achieved differently in the two systems. In cultured cells, it is achieved at the protein level, by adding a targeting sequence to Halo. We discuss the NLS-Halo (nucleus), NES-Halo (cytosol), MOMLS-Halo (mitochondrial outer membrane), and ERt-Halo (endoplasmic reticulum membrane) as examples. In *C. elegans*, it is achieved at the transcriptional level by a organ- or cell-type-specific promoter. For instance, GFP-Halo expression is driven by *myo-2p* (pharyngeal muscle), *ges-1p* (intestine), or *myo-3p* (body-wall muscle). Other promoters and localization sequences are also applicable as needed. For the strategies of plasmid construction, refer to the detailed discussion of ligase-free cloning steps in the Z-REX protocol^25^.

#### C. elegans maintenance

We describe the use of two standard nematode growth media: MYOB plates^63^ and standard Nematode Growth Medium (NGM) plates^64^. The distinctions between these media lie in their concentrations of Mg^2+^ and Ca^2+^ and phosphate, as well as their final pH. Wild-type N2 strains grow robustly on both media, certain mutant backgrounds may exhibit divergent phenotypes due to the lack of divalent ions or phosphate. This differential response can be strategically leveraged during phenotypic screening for the functional importance of a given target under varying micronutrient conditions.

An important factor in all *C. elegans* experiments is the precise control of the culture environment. The physiological states of animals, particularly behavioral phenotypes, are sensitive to the quality of the bacterial food source and the presence of microbial contaminants. As some bacteria remain viable within the worm intestine, contamination can be readily disseminated throughout a population and is difficult to remove via simple picking and transfer. Frequent population synchronization/bleaching^65^, as necessary, should be performed to ensure a clean, homogenous starting culture. Additionally, the *E. coli* strain used as well as its viability can influence worm physiology (e.g., RNAi delivery through feeding^66, 67^). Therefore, maintaining consistency in food preparation and condition is essential for reproducibility in related assays, especially phenotypic characterizations. Two *E. coli* strains are used and they are not interchangeable. OP50, a uracil auxotroph, is the standard food source for routine maintenance and for all Localis-REX experiments. HT115(DE3) is RNase III-deficient and IPTG-inducible, and is used only where double-stranded RNA is delivered by feeding, in the RNAi phenotypic screening steps, such as survival, motility, and fertility test. RNAi does not form part of target discovery. Localis-REX returns a ranked list of locale-specific electrophile responders. Performing RNAi then knocks down individual candidates so that their contribution to organism-level phenotypes can be scored, with and without electrophilic challenge (Step 183). Furthermore, the thickness of the bacterial lawn directly affects defecation intervals. Therefore, in this protocol, we control the optical density of the bacterial culture used for seeding and standardize the drying time of fresh plates at room temperature.

#### *C. elegans* transgenic strain generation

We provide details for several transgenic strain that we have used, however, users may want to generate new transgenics. A transgenic strain with Halo expression is necessary to perform Localis-REX in *C. elegans*. The transgenic strains are routinely created using microinjection of DNA into the gonad^68^ into young adult worms, as we have described^69^. This approach creates worms with an extrachromosomal array that consists of many thousands of concatenated plasmids. For this reason, several plasmids can be co-injected and will all be incorporated into the final array (for instance a Halo expressing plasmid and a marker plasmid). Typically, transgenic progeny are marked by a fluorophore or a dominant phenotypic marker (e.g. *rol-6*^68^). Arrays are transmitted onto subsequent generations at a defined frequency particular to a specific array-typically arrays with transmission of >60% are considered useful for biochemical experiments, although the higher the better. When working with multiple strains, ideally transmission percentages should be matched. A general rule of thumb is that arrays show mosaic expression at a similar frequency to the inheritance transmission frequency. During routine maintenance of transgenic lines with extrachromosomal arrays, frequent selection is necessary to prevent loss of the desired genotype due to competition with background strains, especially before stock preparation.

For biochemical experiments such as those in this protocol, it is often advantageous to integrate the array into the *C. elegans* genome. In these instances, transmission in a homozygotic strain rises to 100 percent, and mosaicism is lost. Integration is usually achieved by irradiating the worms to introduce DNA damage^70^, and then screening for worms that show 100% transmission frequency. For this procedure, it is usually a good idea to start with arrays that transmit at around 40% frequency to aid identification of integrated worms. Integrated transgenic lines should be backcrossed with the parental WT strain at least five times to minimize mutations introduced during X-ray or gamma irradiation.

#### *C. elegans* homozygous knock-in line generation

Knock-in lines are not required to perform Localis-REX. However, they can be crucial at the downstream validation stage, where the cysteine implicated in a candidate responder is mutated at the endogenous locus to test whether an observed phenotype depends on that residue. We used this approach for cyp-33e1(C295A) and cyp-33e1(C439A)^11^. Like the array-containing worms above, these worms can be readily created by Biotechnology companies, such as SUNY.

#### *C. elegans* age and developmental stage selection

The specific experimental objective determines the developmental stage of the worms used. Users unfamiliar with *C. elegans* staging should consult WormAtlas (wormatlas.org) and the WormBook (wormbook.org) developmental staging chapter. In practice, the two discriminators used in this protocol are the characteristic vulval morphology of L4 hermaphrodites and the presence of embryos in adults. For initial proteomic discovery and mass spectrometry sample preparation with Localis-REX photocaged-probe treatment (steps 33-40), mixed-stage worm populations are collected. In contrast, most imaging, functional validation, and phenotypic assays are performed with age-synchronized populations (Steps 9-15). External stimuli or treatments are typically applied during the L4 to young adult stage, such as bolus HNE(alkyne) exposure (step 183). For RNA interference, initiation is commonly performed from the L1 larval stage. While this protocol focuses on L1-stage initiation, the specific RNAi strategy should be determined according to experimental requirements. For example, previous studies performed parental RNAi by exposing the maternal (P0) generation and analyzing the F1 progeny, often producing stronger phenotypes, such as increased developmental arrest or lethality. For controls, let-767 RNAi is used as a positive control, which induces arrest at the L2–L3 stage and subsequent lethality^71^. The empty L4440 vector serves as a negative control. The RNAi feeding library employed in this protocol is derived from the Ahringer library^67, 72^.

#### *C. elegans* male enrichment and setting up genetic crosses

Crossing is required in this protocol when an organ-specific GFP-Halo driver must be combined with a second genetic background, or to generate new strains under multiple genetic backgrounds for phenotypic validation. For example, when introducing a fluorescent reporter (LIU1 strain) to a candidate cysteine knock-in line, or when recovering a strain that has lost its extrachromosomal array. The considerations below apply in those cases, and strains are otherwise propagated by self-fertilization.

Male induction is essential to set up genetic crosses, as males represent only ∼0.1% of the general N2 population^73^. Several stress conditions increase the male frequency, including chronic irradiation or heat shock. Heat stress is the most commonly used method, as described in Step 16. It typically generates a detectable (2 – 5 %) male population in most strains^74^. There is an inverse relationship between male frequency and overall fertility under heat stress. (i.e., harsher conditions tend to generate a higher male percentage but fewer total progeny). Two recommended heat-shock times and temperatures are described in this protocol, though both conditions can be adjusted according to specific needs. To improve chances of finding males, a larger number of starting L4 hermaphrodites is also possible. However, the progeny needs to be divided to avoid overcrowding, which complicates male selection and picking.

When working with fluorescent (such as GFP-halo as a dominant fluorescent marker) or disease-model strains, it is preferable to induce males from fluorescent strains without mobility or viability defects. In many cases, hermaphrodites also produce a proportion of self-crossed progeny during the genetic crossing. Because fluorescent transgenes are mostly dominant genes, fluorescence in the F1 then distinguishes cross-progeny from the self-progeny that are produced alongside them. Where the intended male parent carries a marker associated with impaired movement or altered locomotion (e.g. *rol-6(su1006)*, a widely used dominant co-injection marker), crossing efficiency is reduced, and it is preferable to use that strain as the maternal parent.

Because *C. elegans* employs an XO (male) and XX (hermaphrodite) sex determination system^75^, crosses between hermaphrodites and males yield a higher proportion of viable males among the progeny, and a male stock can be maintained by serial crossing (Steps 18-22). A stock of males can thus be maintained by picking ∼20 young males and 3-4 L4 hermaphrodites at each passage. Males can also be frozen using standard freezing protocols given in Steps 28-31. In addition, males in the F1 generation following heat shock are often less fertile than those in the F2 generation. It is thus recommended to use F2, or above, males when setting up crosses.

When setting up genetic crosses, it is helpful to know the chromosomal locations of the target genes, as this allows researchers to predict the resulting phenotypes and potential difficulties in crossing. Genes located on different chromosomes are generally preferred, because genetic linkage can impede the creation of homozygous doubly transgenic lines.

#### Validations of requisite transgenic *C. elegans* strains including homo-versus heterozygosity differentiation and point mutations

The requirement for stable homozygosity in transgenics depends heavily on the downstream experimental applications. For mass spectrometry-based localis-REX and ultraID hits identification, integrated homozygous strains is not strictly necessary. Strains with extrachromosomal arrays can generally provide sufficient protein yields for proteomics analysis, provided that the enriched proteins are validated by SDS-PAGE with Coomassie staining or straptavidin blot. For strains with limited Halo or ultraID expression (e.g., expression in specific neuronal populations), using stable integrated strains is still highly recommended.

Conversely, for rigorous phenotypic assays and functional validations, the use of stable, homozygous integrated strains is essential. Except where homozygotes are not viable, it is critical that experimental *C. elegans* strains be homozygotes, as key readout signals from fluorescent protein, or other related phenotypes may differ between homozygous and heterozygous animals. Furthermore, due to self-fertilization, it is impossible to obtain and maintain stable heterozygous populations.

Homozygosity of a dominant fluorescent marker can be assessed phenotypically. A heterozygous animal, when self-fertilized, will produce some non-fluorescent progeny, whereas a homozygous animal will produce 100% fluorescent progeny. Strains expressing Halo or Halo-tagged proteins can be stained with Halo-TMR ligand, producing a similar phenotype to dominant fluorescent markers. Most integrated transgenes or mutations (e.g. knock-in alleles via CRISPR-Cas9^76^) do not display any obvious phenotypic differences between homozygous and heterozygous animals. In these cases, validation is typically achieved by genotyping single worms. Individual worms are lysed, and the genomic region of interest is amplified via PCR. For insertions or deletions, the product size difference can be visualized on an agarose gel. For point mutations or small indels, PCR products must be purified and confirmed by Sanger sequencing.

#### Imaging and movies of live and fixed *C. elegans*

Proficiency in the manipulation of *C. elegans* samples is essential, particularly for imaging procedures. For high-quality image acquisition, we recommend aligning multiple animals on a single agarose pad. During live imaging, motility is commonly reduced by levamisole treatment (1 - 5 mM in M9 buffer). This applies to the general recording of fluorescent protein expression in live animals and TMR-Halo staining and imaging workflows in the Procedure (Steps 174-177). However, be careful as paralysis with levamisole in aged worms (day 5 adults or older) can induce intestine extrusion through the vulva and compromise the entire sample. Additionally, worms exhibit an initial contraction phase followed by relaxation into paralysis after levamisole application, resulting in slight morphological changes. To ensure consistency and reproducibility, the time interval between levamisole addition and image capture should be standardized, together with minimizing mechanical perturbations and sample repositioning.

#### Endogenously biotinylated proteins in *C. elegans*

Endogenously biotinylated proteins are abundant in *C. elegans* and present a challenge for biotin-based labeling strategies, such as streptavidin-based enrichment. Prominent endogenous biotinylated proteins include mitochondrial carboxylases such as PCCA-1, PYC-1, and MCC-1^77^, which are consistently detected in streptavidin blots, SDS-PAGE and staining for enriched proteins, and mass spectrometry analyses. These strong background signals can reduce assay sensitivity.

To minimize this interference, we performed preclearance steps for OS-Localis-REX samples, in which endogenously biotinylated proteins are depleted by streptavidin beads prior to the Click reaction. In OS-ultraID workflows, where biotinylation occurs before lysis, animals are fed MG1655 bioB::Kan *E. coli* strain (JLFb1) to minimize exogenous biotin^78^, and two negative control groups are included. One is without extra biotin treatment, and one is without ultraID expression. These ultraID datasets should be interpreted as local abundance maps obtained under the defined MG1655 bioB::Kan feeding condition.

#### SILAC labeling for Localis-REX in live cells

Local electrophile-responsive proteins are key sensors of electrophiles released by Localis-REX in live cells and initiate downstream signaling pathways. To identify such privileged sensors in mammalian cells, we include a procedure for Stable Isotope Labeling by Amino acids in Cell culture (SILAC, steps 97-100, 105-107)^38^ combined with LC-MS/MS. Cells are cultured with either D_4_-Lysine and ^13^C_6_ Arginine (“Medium” label), ^13^C_6_, ^15^N_2_ Lysine and ^13^C_6_ or ^15^N_4_ Arginine (“Heavy” label), or unlabeled Lysine and Arginine (“Light” label). Localis-REX is then performed in Light, Medium, or Heavy SILAC cells, with appropriate isotope-labeled controls. The Medium group can be omitted if only two conditions are designed to be compared. All conditions are pooled, followed by biotin pull-down to enrich potential protein targets. Digested peptides from Localis-REX samples are subsequently analyzed by LC-MS/MS, and target proteins are distinguished by characteristic mass shifts relative to controls. Moreover, label-swapping between Localis-REX and control conditions enables reciprocal verification, further strengthening target identification. This is an optional extension and is not covered by the steps below.

#### Protein candidate selection and validation following Localis-REX in live cells and *C. elegans*

Following the identification of putative REM-responders via Localis-REX, candidates for downstream phenotypic validation need to be prioritized. This can be done by numerous criteria that may fit with a specific research question, or laboratory focus. However, for unbiased screens, we use the following criteria: (1) presence of human orthologs with disease relevance; (2) no previously known electrophile sensitivity or druggability; and (3) conservation of at least one cysteine residue in both human and *C. elegans* orthologs. The candidates selected are recommended to reflect both unique and common hits across regions analyzed (**Fig. 1c**).

#### Phylogenetic and sequence conservation analyses during Localis-REX hit selection process

Following a primary Localis-REX screen, phylogenetic and sequence conservation analyses can be applied to candidate proteins, which often implies a fundamental and conserved biological role. The analysis pipeline begins by identifying orthologs of sensor protein candidates across a specific set of species (e.g., representative metazoan species from *C. elegans* to *H. sapiens*) in phylogenomic databases (e.g., OrthoDB, OMA, eggNOG). For species like human, mouse, frog, zebrafish, and *C. elegans*, Alliance of Genome Resources (AGR) compares the orthologs across multiple databases and algorithms. These sequences are then subjected to multiple sequence alignment using Clustal Omega^79^ or MUSCLE^80^ to identify conserved domains and specific residues. The resulting alignments are subsequently used to construct a phylogenetic tree using methods such as Neighbor-Joining or Maximum Likelihood (e.g., with MEGA^81^ or PhyML^82^ software). Specific sequences of interest can be analyzed, for instance by sequence logo using WebLogo _3_83.

#### Live-cell-based Localis-REX coupled to protein translation reporter assays

How changes in localized metabolite build-up impact specific cellular processes remain challenging to interrogate in living systems. Coupling Localis-REX’s ability to generate REMs with absolute timing and spatial control to a specific reporter assay addresses this major technology gap. Capturing nascent protein synthesis is challenging, due to highly dynamic mRNA translation and limited methods to identify newly synthesized protein from the large proteomic pool. The best methods allow selective incorporation of an orthogonal handle into nascent proteins to allow quantification of synthesis. We exemplify this capability by applying Localis-REX in tandem with established reporter assays for protein synthesis, namely: genetic code expansion (GCE) method reporting either a specific protein-targeted (Steps 136-141)^84^, or whole proteome-targeted, translation efficiency; and biorthogonal non-canonical amino acid (BONCAT) method-based nascent proteome remodeling efficiency (Steps 143-152)^85^.

Orthogonal aminoacyl-tRNA synthetase/tRNA pairs have been introduced in cells to recognize stop codons, such as amber stop codon. Such “suppressors” allow incorporation of an unnatural amino acid into a specific position within proteins of interest. In this paper, we use a synthetic HA-tagged Actin, in which codon for K118 is replaced by an amber codon-UAA as a translation reporter (HA-tagged Actin(K118TAG)). Mammalian cells are co-transfected with Halo for Localis-REX and MmPyIRS/ HA-Actin(K118TAG) pair for GCE. After the uncaging step is complete (Step 105), cells are incubated with Bicyclo[6.1.0]nonyne-Lysine (BCNK) is introduced to allow translation of HA-Actin(K118BCNK). HA-Actin(K118BCNK) translation can be assessed by either Cy5 in-gel fluorescence, or western blot measuring anti-HA signal corresponding to HA-Actin(K118BCNK). HA-Actin(K118*) is unstable and is not detected.

Whole proteome translation can also be measured using an analogous strategy. In this case, we express an orthogonal Valyl-tRNA synthetase, Val-MmPylRS, which incorporates a tRNA recognizing the Val codon, replacing valine with pyrrolysine-or Clickable analogs, such as BCNK across the whole proteome. The BCNK incorporated nascent proteome can be visualized by in-gel fluorescence following Diels-Alder reaction with Cy5-azide.

Biorthogonal Non-canonical Amino Acid Tagging (BONCAT) utilizes L-azidohomoalanine or L-homopropargylglycine to replace methionine that labels and tracks newly synthesized proteins in the cells. In our case, post Localis-REX, cells are incubated in methionine dropout medium, supplemented with L-azidohomoalanine (50 µM), 4-hour incubation allows L-azidohomoalanine to incorporate into the newly synthesized proteome. Exploiting the azide function on L-azidohomoalanine, Click chemistry with Cy5-alkyne probe is performed following cell lysis. Nascent proteins are further visualized and quantified through in-gel fluorescence.

Additional specific considerations are provided below within the corresponding experimental steps.

##### Box 1: Recommended positive, negative, and quality controls

The following controls should be integrated in Localis-REX and ultraID experiments to ensure data validity, rigor, and interpretability.

- ‘No-alkyne’ control for Localis-REX MS-based target mapping experiments: this control, i.e., the use of Ht-PreHNE in place of Ht-PreHNE(alkyne), establishes the baseline of nonspecific enrichment following Click coupling to biotin-azide and subsequent streptavidin pulldown; defines the denominator of the TMT fold-change ratio; and is essential for Gaussian threshold establishment.
- ‘No light’ control [+Ht-PreHNE(alkyne), –UV], ‘no-probe’ control [–Ht-PreHNE(alkyne), +UV], and ‘vehicle DMSO alone’ control [–Ht-PreHNE(alkyne), –UV], representing technical negative controls of Localis-REX-mediated proteomics-based spatiofunctional mapping are essential for the following assays: in-gel fluorescence validation and biotin pulldown validation (by streptavidin blots or Coomassie Blue staining). These controls collectively verify that *in vivo* REM generation, capturing REM-responsive proteins, and REM-dependent downstream cellular and phenotypic responses are NOT triggered inadvertently by photocaged probe alone or light exposure alone or DMSO treatment.
- Pulldown robustness and reproducibility validation for Localis-REX or ultraID proteomics: prior to proteomics sample processing and data acquisition, reproducible outcomes from enrichment leveraging above-mentioned streptavidin blot- or Coomassie Blue gel-based readouts should be performed (**Extended Fig. 1**).
- Bulk HNE(alkyne) treatment also serves as a useful technical positive control in above in-gel fluorescence and biotin pulldown assays, particularly for researchers new to these workflows.
- In Localis-REX-mediated downstream phenotypic mapping assays, the above controls/experiments are conducted with Ht-PreHNE as opposed to Ht-PreHNE(alkyne), such that functional signaling and phenotypic consequences triggered by the native unfettered reactive metabolite HNE are evaluated.
- Wild-type *C. elegans* control for worm imaging and ultraID locale-specific abundance mapping: This control serves as background signal reference point for non-transgenic animals for both imaging (background autofluorescence) and proteomics outcomes (non-specific MS hits or hits altered due to transgene expression).
- ‘No external biotin’ control for ultraID locale-specific abundance mapping: this control affirms that hits captured from proximity-dependent biotinylation is induced by exogenous biotin addition, as opposed to identification of endogenously biotinylated proteins prevalent in *C. elegans*.
- ‘Localization’ validation for Localis-REX and ultraID proteomics mapping experiments: before proceeding with respective proteomics mapping experiments, this control helps confirm correct subcellular localization of Halo or ultraID protein using fluorescence microscopy.
- Dosage and treatment time determination for photocaged-REM probe or exogenously administered biotin: in Localis-REX, this assessment determines minimum probe concentration required to saturate Halo binding site *in vivo* and in live cells across different tissue-specific Halo transgenic worm strains or subcellular locales respectively. In ultra-ID, this assessment helps to optimize exogenous biotin concentration and treatment duration across different ultraID transgenic worm strains.

**[Production end of Box 1]**

## MATERIALS

### BIOLOGICAL MATERIALS

#### BACTERIA LINES

OP50

HT115(DE3)

JLFb1 (MG1655 bioB::kan)

These strains are available from Caenorhabditis Genetics Center (CGC).

#### WORM LINES

Wild-type (N2),

LIU1 (*dhs-3p::dhs-3::gfp*), outcrossed x 6

IG1839 (*frSi17 [mtl-2p::rde-1::rde-1 3’UTR] **II**; frIs7 [nlp-29p::gfp + col-12p::DsRed] **IV**; rde-1(ne300) **V***), outcrossed x 0

WM118 (*neIs9 [myo-3::HA::rde-1::rde-1 3’UTR + rol-6(su1006)] **X**; rde-1(ne300) **V***)

CYA03 (*rexEx1 [myo-2p::gfp::halo::unc-54 3’UTR + mec-7p::mRFP]*)

CYA04 (*rexEx2 [ges-1p::gfp::halo::unc-54 3’UTR + mec-7p::mRFP]*)

CYA05 (*rexEx3 [myo-3p::gfp::halo::unc-54 3’UTR + mec-7p::mRFP]*)

CYA06 (*rexEx4 [myo-2p::mCherry::P2A::ultraID::unc-54 3’UTR]*)

CYA07 (*rexEx5 [ges-1p::mCherry::P2A::ultraID::unc-54 3’UTR]*)

CYA08 (*rexEx6 [myo-3p::mCherry::P2A::ultraID::unc-54 3’UTR]*)

These strains are available from Caenorhabditis Genetics Center (CGC), or upon request, from the Aye laboratory, Oxford (UK).

#### CELL LINES

HEK293T/17 (ATCC CRL-11268; RRID:CVCL_1926)

N2A (ATCC CRL-131; RRID:CVCL_0470)

**CRITICAL** The cell lines used in your research should be regularly checked to ensure they are authentic and are not infected with mycoplasma.

### REAGENTS

#### Reagents for cells or *C. elegans* maintenance and validation

- DMEM (e.g., GIBCO cat. no. 41965039)
- MEM, + GlutaMAX (e.g., GIBCO cat. no. 41090036)
- Opti-MEM (e.g., GIBCO cat. no. 51985026)
- MEM Non-Essential Amino Acids Solution (100X) (NEAA) (e.g., GIBCO, cat. no. 11140035)
- Penicillin-Streptomycin (Pen-Strep, 10,000 U/mL) (e.g., GIBCO, cat. no. 15140122)
- DPBS, no calcium, no magnesium (e.g., GIBCO, cat. no. 14190094)
- Sodium Pyruvate (100 mM) (e.g., GIBCO, cat. no. 11360039)
- TrypLE Express Enzyme (e.g., GIBCO, cat. no. 12605028)
- Fetal Bovine Serum (FBS) (Sigma-Aldrich, cat. no. F0392)
- Dimethyl sulfoxide (DMSO) (e.g., Sigma-Aldrich, cat. no. D8418; CAS: 67-68-5)
- Isopropanol (e.g., Sigma-Aldrich, cat. no. 278475; CAS: 67-63-0)
- K_2_HPO_4_ (e.g., Sigma-Aldrich, cat. no. P3786; CAS: 7758-11-4)
- KH_2_PO_4_ (e.g., Sigma-Aldrich, cat. no. P0662; CAS: 7778-77-0)
- CaCl_2_ (e.g., AppliChem, cat. no. A4689; CAS: 10035-04-8)
- ZnSO_4_ (e.g., Sigma-Aldrich, cat. no. Z0251; CAS: 7446-20-0)
- FeSO_4_ (e.g., Sigma-Aldrich, cat. no. F8633; CAS: 7782-63-0)
- MnCl_2_ (e.g., Sigma-Aldrich, cat. no. M5005; CAS: 13446-34-9)
- CuSO_4_ (Sigma-Aldrich, cat. no. 209198; CAS: 7758-99-8)
- Na_2_HPO_4_ (e.g., Sigma-Aldrich, cat. no. S9390; CAS: 7782-85-6)
- Na_2_EDTA (e.g., Sigma-Aldrich, cat. no. E5134; CAS: 6381-92-6)
- MgSO_4_ (e.g., Thermo Fisher Acros, cat. no. 10003812; CAS: 7487-88-9)
- Glycerol (e.g., Thermo Fisher Acros, cat. no. 158920100; CAS: 56-81-5)
- NaCl (e.g., AppliChem, cat. no. A2942; CAS: 7647-14-5)
- CaCl_2_ (e.g., AppliChem, cat. no. A4689; CAS: 10035-04-8)
- Citric acid monohydrate (e.g., Sigma-Aldrich, cat. no. C1909; CAS: 5949-29-1)
- Potassium citrate tribasic monohydrate (e.g., Sigma-Aldrich, cat. no. C3029; CAS: 6100-05-6)
- Yeast extract (e.g., AppliChem, cat. no. A3732)
- Tryptone (e.g., Apollo Scientific, cat. no. BIT1332)
- Agarose (e.g., Apollo Scientific, cat. no. BIA1176; CAS: 9012-36-6)
- Tris(hydroxymethyl)aminomethane (Tris) (e.g., Apollo Scientific, cat. no. BI2888; CAS: 77-86-1)
- Sodium hypochlorite solution 10-14% (Bleach) (e.g., Fisher Scientific, cat. no. 15450837; CAS: 7681-52-9)
- HCl (e.g., Reactolab, cat. no. 99400; CAS: 7647-01-0)
- KOH (e.g., Reactolab, cat. no. 99510; CAS: 1310-58-3)
- Ampicillin (e.g., AppliChem, cat. no. A0839; CAS: 69-52-3)
- Carbenicillin (e.g., AppliChem, cat. no. A1491; CAS: 4800-94-6)
- Tetracycline (e.g., AppliChem, cat. no. A2228; CAS: 64-75-5)
- Isopropyl β-D-1-thio-galactopyranoside (IPTG) (e.g., Goldbio, cat. no. I2481C25; CAS: 16758-34-2)
- Standard reagents for DNA gel electrophoresis
- SYBR Safe DNA Gel Stain (Thermo Fisher Invitrogen, cat. no. S33102)
- Proteinase K (Sigma-Aldrich, cat. no. P2308)
- Phusion HF Buffer Pack (Thermo Fisher, cat. no. F518L)
- Phusion HotStart II polymerase (Thermo Fisher, cat. no. F549L)
- Set of dATP, dCTP, dGTP, dTTP (Promega, cat. no. U1240)

#### Reagents for cells or *C. elegans* imaging

- Paraformaldehyde (PFA) (e.g., Sigma-Aldrich, cat. no. P6148; CAS: 30525-89-4) **! CAUTION** Carcinogen: Handle inside fume hood with gloves, goggles, and lab coat, when preparing formaldehyde solution. Do not heat in sealed bottles.
- Oil Red O solution (e.g., Sigma-Aldrich, cat. no. O1391; CAS: 1320-06-5)
- Methanol (e.g., Thommen Furler, cat. no. 203-VL54K; CAS: 67-56-1)
- Tween-20 (e.g., Sigma-Aldrich, cat. no. P9416; CAS: 9005-64-5)

#### Reagents for Localis-REX

- 4-(2-hydroxyethyl)-1-piperazineethanesulfonic acid (HEPES) (e.g., Apollo Scientific, cat. no. BI8181; CAS: 7365-45-9)
- HaloTag-targetable precursor to HNE(alkyne) and HNE(no alkyne) [Ht-PreHNE(alkyne) and Ht-PreHNE(no alkyne) (see original protocol^24^)]

**! CAUTION** Although these compounds do not have observable toxic or harmful effects toward live animals such as larval zebrafish and *C. elegans* at the dosages being applied^11, 25, 26, 69^, their potential adverse effects on man have not been actively assessed. Thus, they should be handled with the usual standard of care and good laboratory practice in chemical reagent handling, whereby researchers must be equipped with standard personal protective equipment (gloves, safety goggles, and a lab coat).

**! CRITICAL**

The stock of Ht-PreHNE(alkyne) and Ht-PreHNE(no alkyne) should be stored as one-shot aliquots in DMSO at −80 °C. Stock concentration of probe should be at least 30 mM to make sure final DMSO in Medium is less than 0.05%.

- HNE(alkyne) (see original protocol^24^)
- cOmplete, Mini, EDTA-free Protease Inhibitor Cocktail (e.g., Roche, cat. no. 11836170001)
- Zirconia beads (e.g., BioSpec cat. no. 11079107zx)
- *t*-BuOH (e.g., Sigma-Aldrich, cat. no. 471712; CAS: 75-65-0)
- Sulfo-cyanine5 (Cy5)-azide (e.g., Lumiprobe, cat. no. B3330)
- Biotin-dPEG11-azide (e.g., Iris Biotech, cat. no. PEG4340; CAS: 956494-20-5)
- Cu(II)-TBTA, 10 mM in 55% DMSO (Lumiprobe, cat. no. 21050)
- TCEP·HCl (e.g., Goldbio, cat. no. TCEP25; CAS: 51805-45-9)
- (+)-Sodium L-ascorbate (e.g., Sigma-Aldrich, cat. no. A4034; CAS: 134-03-2)
- Lithium dodecyl sulfate (LDS) (e.g., Sigma-Aldrich, cat. no. L9781; CAS: 2044-56-6)
- Sodium dodecyl sulfate (SDS) (e.g., Apollo Scientific, cat. no. BIB2008; CAS: 151-21-3)
- β-mercaptoethanol (BME) (e.g., Sigma-Aldrich, cat. no. M3148; CAS: 60-24-2)

**! CAUTION** BME must be handled exclusively in a fume hood with standard personal protective equipment.

- Triton X-100 (e.g., Sigma-Aldrich, cat. no. X100; CAS: 9036-19-5)
- Standard reagents for protein gel electrophoresis and western blotting
- Bradford Dye (e.g., BioRad cat. no. 5000205)
- Bovine Serum Albumin (BSA) (e.g., Fisher, cat. no. BP1600-100)
- Bovine Serum Albumin Standard Ampules (Thermo Fisher, cat. no. 23209)
- His6-TEV-S219V (see original protocol^24^)
- Anti-β-actin-HRP conjugated primary antibody (Sigma, cat. no. A3854; RRID:AB_262011)
- Anti-HaloTag monoclonal antibody (Promega, cat. no. G9211; RRID:AB_2688011)
- Anti-HaloTag pAb from Rabbit (Promega, cat. no. G9281; RRID:AB_713650)
- GFP antibody (B-2) (Santa Cruz Biotech, cat. no. sc-9996; RRID:AB_627695)
- DYKDDDDK polyclonal antibody (Thermo Fisher, cat. no. PA1-984B; RRID:AB_347227)
- Anti-GFP antibody (Abcam, cat. no. ab6673; RRID:AB_305643)
- Donkey anti-Goat IgG H&L (Alexa Fluor® 488) (Abcam, cat. no. ab150133; RRID:AB_2832252)
- Goat Anti-Rabbit IgG H&L (Alexa Fluor® 488) (Abcam, cat. no. ab150077; RRID:AB_2630356)
- Goat Anti-Mouse IgG H&L (Alexa Fluor® 568) (Abcam, cat. no. ab175473; RRID:AB_2895153)
- Streptavidin-HRP (Abcam, cat. no. ab7403)
- Anti-rabbit IgG, HRP-linked Antibody (Cell Signaling Technology, cat. no. 7074S; RRID:AB_2099233)
- Anti-mouse IgG, HRP-linked Antibody (Cell Signaling Technology, cat. no. 7076S; RRID:AB_330924)
- Anti-Calnexin monoclonal antibody (Santa Cruz Biotech, cat.no. sc-23954)
- MitoTracker Red (Thermo Fisher, cat. no. Cam7512)
- High capacity streptavidin agarose (Thermo Scientific cat. no. 20359)
- Acetone (e.g., Thommen Furler, cat. no. 133-VL54K; CAS: 67-64-1)

#### Reagents for SILAC

- SILAC drop-off media 1X DMEM (Thermo Fisher # 89985)
- 10% dialyzed FBS (Sigma Aldrich # F0392)
- L-Lysine 2HCl (4,4,5,5-D_4_) (Cambridge Isotope Lab, DLM-2640-PK)
- L-Lysine 2HCl (^13^C_6_, ^15^N_2_) (Cambridge Isotope Lab, CNLM-291-H-0.1)
- L-Arginine HCl (^13^C_6_) (Cambridge Isotope Lab, CLM-2265-H-PK)
- L-Arginine HCl (^13^C_6_, ^15^N_4_) (Cambridge Isotope Lab, CNLM-539-H-0.1)

#### Reagents for BONCAT experiment

- DMEM no methionine (Thermo Fisher 21013024)
- L-Azidohomoalanine hydrochloride (Sigma-Aldrich, 900892)

#### Reagents for GCE experiment

- BCNK (Bicyclo[6.1.0] nonyne-Lysine) (Sichem, sc-8016)

**CRITICAL** BCNK should be stored as single use aliquots at −80 °C. Prepare a 100 mM stock solution in 100 mM NaOH. Aliquots are typically stable for at least 6 months.

- Tetrazine-Cy5 (Lena Bioscience, CLK-015-05)
- Cyanine5 (Cy5) alkyne (e.g., Lumiprobe, cat. no. B30B0)

#### Reagents for mass spectrometry

- Acetonitrile, LC-MS grade (e.g., Sigma-Aldrich, cat. no. 1000291000; CAS: 75-05-8)
- Formic acid, LC-MS grade (e.g., Sigma-Aldrich, cat. no. 5330020050; CAS: 64-18-6)
- TMT6plex isobaric labeling reagents (Thermo Fisher, cat. no. 90066)
- TMT11plex isobaric labeling reagents (Thermo Fisher, cat. no. A37725)
- Trypsin, LC-MS grade (Thermo Fisher, cat. no. 90058)

#### Plasimds and Oligos

- pWormgate2-myo-2p-gfp-halo (Addgene 227169)
- pWormgate2-ges-1p-gfp-halo (Addgene 2271670)
- pWormgate2-myo-3p-gfp-halo (Addgene 227171)
- pWormgate2-myo-2p-mCherry-P2A-flag-UltraID (Addgene 227172)
- pWormgate2-ges-1p-mCherry-P2A-flag-UltraID (Addgene 227173)
- pWormgate2-myo-3p-mCherry-P2A-flag-UltraID (Addgene 227174)
- pCS2+8 NLS-Halo (available upon request)
- pCS2+8 NES-Halo (available upon request)
- pCS2+8 MOMLS-Halo (available upon request)
- pCS2+8 ERt-Halo (available upon request)
- Actin(K118TAG) (available upon request)
- MmPylRS (available upon request)
- Val-MmPylRS (available upon request)
- L4440 (Addgene 1654)
- 295A-seq-s: GAACCAATGGGAGGACTTA
- 295A-seq-a: TAACCGATAGAGGCAATGA
- 439A-out-s: AGTGGAAATCGGTCCGTTGG
- 439A-out-a: ACTATCACCGCCGGAGTCAA

**CAUTION!** Regarding organic solvents, acids, bases, and other chemical reagents, flammable solvents (methanol, acetone, etc.) must be kept away from open flames and heat sources. DMSO penetrates skin rapidly and may carry dissolved reagents; change gloves promptly after contact. Avoid skin and eye contact with acids (HCl) and bases (KOH). Consult the specific Safety Data Sheets for other chemical reagents, including components that come with molecular biology kits, and handle all reagents with appropriate personal protective equipment.

### EQUIPMENT

- Standard equipment for protein electrophoresis and blotting (e.g. Bio-Rad)
- Standard equipment for nucleic acid agarose gel electrophoresis (e.g. Bio-Rad)
- Pipettes (e.g., Pipetman or Eppendorf)
- Thermal cycler (e.g. Applied Biosystems Veriti or Bio-Rad T100)
- Hand-held UV lamp with 365-nm light (e.g., Spectroline ENF 240C)

**CAUTION!** Although the hand-held low-power lamp (5 mW/cm²) alongside short illumination time (≤ 5 min) used in this protocol poses minimal hazard, exposing unprotected eyes directly to the lamp source should be avoided. We refer readers to accompanying movie clips in this protocol and associated protocol T-REX^24^. During pre-warming period, covering the lamp with aluminum foil to shield the beam path is recommended. UV-protected goggles are also recommendable.

- Accumax UV meter (Spectroline, cat. no. XS-365)
- Petri dishes and multiwell plates (e.g., VWR cat. no. 25384-164 and 25384-088)
- Tissue culture dishes and multiwell plates (e.g., TPP tissue culture dishes)
- Glass Pasteur pipette (e.g., Fisher cat. No. 11765098)
- Fluorescence stereomicroscope (e.g., Leica M205 FA, with FluoCombi III accessory)
- Fluorescence microscope light source (e.g., Lumencor SOLA U-nIR Light Engine)
- Stereo microscope (e.g., AmScope SE306 Series)
- Glass microscope slides and coverslips
- ChemiDoc-MP imaging system (Bio-Rad) or Fusion FX imager (Vilber)
- Hemocytometer (Hausser Scientific cat. no. 1492) or glass stage micrometer (Meiji Techno. cat. no. MA285)
- HPLC system for LC-MS analysis (e.g., Dionex Ultimate 3000 RSLCnano UHPLC system)
- Mass spectrometer (e.g., Q Exactive HF Orbitrap mass spectrometer, or comparable models with sufficient MS/MS resolution for accurate TMT reporter ion quantification, typically ≥30,000 at *m*/*z* 200)
- Acclaim PepMap 100 C18 capillary trap column (Thermo Fisher, 3 μm, 20 mm, 75 μm ID; cat. no. 164946)
- Acclaim PepMap 100 C18 analytical nanoLC column (Thermo Fisher, 2 μm, 500 mm, 75 μm ID; cat. no. 164942)
- ReproSil-Pur 120 C18-AQ silica beads (Dr. Maisch, 1.9 μm, cat. no. r119.aq)
- Fused silica capillary tubing for in-house analytical column packing, 75 μm ID
- NanoViper fittings/capillaries (Thermo Fisher)
- Plate reader (e.g., Agilent BioTek Cytation 5)
- Vortexer (e.g., Vortex-Genie 2)
- Refrigerated centrifuge (e.g. Eppendorf 5425R)
- Microwave (e.g., cookology)
- Ultrasonic bath (e.g., Diagenode Bioruptor Pico sonication device)

#### SOFTWARE

- ImageJ/FIJI^86^

(NIH, https://fiji.sc)

- LAS X

(Leica Microsystems, https://www.leica-microsystems.com/products/microscope-software/p/leica-las-x-ls/)

- MaxQuant^87^

(MPI Biochemistry, https://www.maxquant.org)

- Proteome Discoverer

(Thermo Fisher, https://www.thermofisher.com/order/catalog/product/PROTEOMEDISC3)

- Perseus^88^

(MPI Biochemistry, https://www.maxquant.org/perseus/)

### REAGENT SETUP

#### *C. elegans* culture and incubation

- **S buffer:**Dissolve 5.85 g NaCl, 1 g K_2_HPO_4_, and 6 g KH_2_PO_4_ into 1 L ddH_2_O, sterilize by 0.22 μm filter and autoclaving. Store this solution at room temperature (20 - 25 °C) for up to 1 year.
- **M9 buffer:**Dissolve 11.3 g Na_2_HPO_4_·7H_2_O (42 mM), 3.0 g KH_2_PO_4_ (22 mM), and 5.0 g NaCl (86 mM) in 1 L ddH_2_O, sterilize by autoclaving. Dissolve 2.5g MgSO_4_·7H_2_O (100 mM, 100x solution) in 100 mL ddH_2_O, sterilize by 0.22 μm filter. Mix 10 mL of 100 mM MgSO_4_ solution and 1 L of phosphate solution to make M9 buffer before use. Store this solution at room temperature for up to 6 months.
- **Stock Making buffer:** Mix S buffer and glycerol to make 30% glycerol (v/v) buffer, sterilize by autoclaving. Store this solution at room temperature for up to 6 months.
- **LB (lysogeny broth) medium:** Dissolve 10 g NaCl, 10 g tryptone, and 5 g yeast extract in 1 L ddH_2_O. Sterilize by autoclaving and store at room temperature for up to 1 month.
- **Trace metals solution (100x):** Dissolve 930 mg Na_2_EDTA·2H_2_O (5 mM), 345 mg FeSO_4_·7H_2_O (2.5 mM), 100 mg MnCl_2_·4H_2_O (1 mM), 145 mg ZnSO_4_·7H_2_O (1 mM), and 12.5 mg CuSO_4_·5H_2_O (1 mM) in 500 mL ddH_2_O. Sterilize by autoclaving and store in the dark at 4 °C for up to 6 months.
- **Potassium citrate (100x):** Dissolve 10 g citric acid monohydrate (95 mM) and 146.8 g tri-potassium citrate monohydrate (905 mM) in 500 mL ddH_2_O. Sterilize by autoclaving and store at 4 °C for up to 6 months.
- **S Complete (S medium):** Mix 1 L S buffer with 1 mL cholesterol (5 mg/mL in EtOH), 10 mL trace metals solution, 10 mL potassium citrate solution, 3 mL sterilized 1 M CaCl_2_, and 3 mL sterilized 1 M MgSO_4_. Do not autoclave and use it freshly.
- **20% alkaline hypochlorite solution**: Prepare freshly before use by mixing 3 mL commercial bleach (between 10 and 14% NaOCl), 3.75 mL 1M NaOH, and 8.25 mL ddH_2_O.

#### Click reaction (for both live-cell- and worm-based assays)

- **Basal HEPES buffer:** Dissolve 11.9 g HEPES (50 mM) and 8.77 g NaCl (150 mM) in 900 mL ddH₂O. Adjust pH to 7.6 using NaOH. Bring the final volume to 1 L with ddH₂O. Sterilize the solution by 0.22 μm filter. Store at 4 °C for up to 6 months (and protected from light); warm up to room temperature prior to use.
- **CuSO_4_ solution:** Dissolve 2.5 g CuSO_4_·5H_2_O (100 mM) in 100 mL ddH_2_O. Sterilize by 0.22 μm filter and store at room temperature for up to 1 year.
- **TCEP stock solution:** Dissolve 2.87 g TCEP·HCl (100 mM) in 100 mL of 500 mM HEPES (11.92 g) (pH pre-adjusted to 7.6). Aliquot and store at −20 °C for up to 6 months. Prevent freeze-thawing.
- **Cy5-azide stock solution:** Dissolve 1 mg sulfo-Cyanine5 azide in 2.59 mL DMSO to make 0.5 mM stock solution. Aliquot as 10 µL stocks and store at −20 °C for up to 6 months. Prevent freeze-thawing.
- **Biotin-azide stock solution:** Dissolve biotin-PEG_n_-azide in DMSO to make 100 mM stock solution (typically 100 mg of biotin-PEG_11_-azide in 1.26 mL DMSO). Aliquot as 10 µL stocks and store at −20 °C for up to 6 months. Prevent freeze-thawing.
- **20% SDS (w/v) solution:** Dissolve 20 g SDS in 100 mL ddH_2_O by stirring overnight at room temperature to ensure complete dissolution. Heating (50 – 60 °C) or sonication can be used to accelerate the dissolution if necessary. Vortexing is not recommended, as it can generate bubbles that trap undissolved solids. Store this solution at room temperature for up to 1 year.
- **16% LDS (w/v) solution:** Dissolve 16 g LDS in 100 mL ddH_2_O by stirring overnight at room temperature to ensure complete dissolution. Heating (50 – 60 °C) or sonication can be used to accelerate the dissolution if necessary. Vortexing is not recommended, as it can generate bubbles that trap undissolved solids. Store this solution at room temperature for up to 1 year.

#### *C. elegans* fixing and imaging

- **10% (w/v) paraformaldehyde (PFA):** Dissolve 20 g PFA powder in 160 mL PBS by stirring and heating to 50 °C in a fume hood. Add saturated NaOH solution slowly until it is clear and then adjusted pH to 7 with HCl based on pH paper. Add extra PBS until the final volume is 200 mL. Aliquot as 4 mL stocks and store at −20 °C for up to 6 months.
- **PDT buffer:** Add 1 mL DMSO (1% v/v) and 100 µL Tween-20 (0.1% v/v) in 100 mL PBS. Stir it until fully homogeneous. Store at room temperature for up to 1 month.
- **Immunofluorescence (IF) imaging blocking buffer:** Add 10 mL (10% v/v) heat-inactivated fetal bovine serum (FBS), 2 g (2% w/v) BSA, and 50 µL (0.05% v/v) Tween-20 in 90 mL PBS. Stir it until fully homogeneous. Store at 4 °C for up to 1 month.
- **Oil red O staining solution:** Mix 30 mL 0.5% Oil Red O in 100% isopropanol and 20 mL ddH_2_O. Stir or rotate end-to-end for at least overnight. Filter the solution by 0.22-μm filter right before the experiment.

**CRITICAL**: Precipitation occurs after mixing Oil Red O with ddH_2_O. Ensure the working solution is free of visible precipitates before application, as particulate matter can adhere to the specimen and interfere with staining results.

#### *C. elegans* live imaging and video recording

- **Levamisole solution:** Prepare 1 M levamisole stock in M9 buffer by dissolving 241 mg levamisole hydrochloride into 1 mL M9 buffer. Aliquot and store the 1 M stocks at −20 °C for up to 1 year. Dilute the 1 M stock to 1-5 mM with M9 for imaging. The diluted levamisole can be stored at 4 °C for up to 1 month.
- 1% agarose solution: Boil (microwave) 1% m/v agarose solution in H_2_O. Store at 4 °C for up to 1 year.

**CRITICAL** Also see **Supplementary Movie 1.**

#### Cultured mammalian cells: live imaging and immunofluorescence imaging

- **Cell immunofluorescence (IF) blocking buffer:** Add 300 mg (3% w/v) BSA, 20 µL (0.2% v/v) Triton X-100 in 10 mL PBS. Stir it until fully homogeneous. Freshly prepare before usage.
- **Cell immunofluorescence (IF) incubation buffer:** Add 100 mg (1% w/v) BSA, 2 µL (0.02% v/v) Triton X-100 in 10 mL PBS. Stir it until fully homogeneous. Freshly prepare before usage.
- **DAPI stock and working solution:** Dissolve 10 mg DAPI (0.1% w/v) in 10 mL water to 1 mg/mL, and filter the solution by 0.22-µm filter, aliquot and store in −20 °C up to 3 months. To prepare working solution, dilute stock of DAPI to 1 µg /mL in PBS. Freshly prepare before usage.
- **Hoechst stock and working solution:** Dissolve 100 mg Hoechst (1% w/v) in 10 mL water to 10 mg/mL, and filter the solution by 0.22-µm filter, aliquot and store in −20 °C up to 3 months. To prepare working solution, dilute stock of Hoechst to 10 µg/mL in rinsing medium. Freshly prepare before usage.

#### SILAC experiments

**Light/Medium/Heavy complete media:** Add 10 mL dialyzed FBS (10% v/v), 1 mL 100X sodium pyruvate, 1 mL 100X Pen-Strep, 0.1 mL 200 g/L L-Proline (final concentration 200 mg/L) in 88 mL SILAC drop-off DMEM, then add different isotopic labeled Lysine and Arginine:

- **Light media:** 0.1 mL of 0.8 mol/L L-Lysine and 0.1 mL of 0.5 mol/L L-Arginine
- **Medium media:** 0.1 mL of 0.8 mol/L D4-L-Lysine and 0.1 mL 0.5 mol/L ^13^C_6_ L-Arginine
- **Heavy media:** 0.1 mL of 0.8 mol/L ^13^C_6_, ^15^N_2_-L-Lysine and 0.1 mL of 0.5 mol/L ^13^C_6_, ^15^N_4_ L-Arginine Store the media at 4 °C for up to 1 month.

#### BONCAT experiments

- **BONCAT rinsing medium:** Add 5 mL 100X sodium pyruvate, 5 mL 100X Pen-Strep to 500 mL of DMEM without methionine. Store the medium at 4 °C for up to 1 month.
- **L-Azido-homoalanine (AHA) stock and working solution:** Add 90.3 mg L-Azidohomoalanine hydrochloride in DPBS, to 50 mmol/L concentration, and filter the solution by 0.22-µm filter, aliquot and store in −20°C up to 3 months. To prepare working solution, dilute stock of AHA to 50 µmol/L in BONCAT rinsing medium right before use.

#### DNA electrophoresis, SDS-PAGE and western blot (for both live-cell- and worm-based assays)

- **Tris-acetate-EDTA (TAE) buffer, 50x:** Dissolve 242 g Tris base (2M), 18.61 g Na_2_EDTA·2H_2_O (50 mM), and 57.1 mL glacial acetic acid (1 M) in ddH_2_O to make 1 L buffer. Store at room temperature for up to 6 months.
- **SDS-PAGE running buffer, 10x:** Dissolve 151.4 g Tris base (2.5 M), 720.7 g Glycine (1.92 M), and 50 g SDS (1% w/v) in ddH_2_O to make 5 L buffer. Store at room temperature for up to 6 months.
- **Transfer buffer, 10x:** Dissolve 151.4 g Tris base (2.5 M) and 720.7 g Glycine (1.92 M) in ddH_2_O to make 5 L buffer. Store at room temperature for up to 6 months.
- **Transfer buffer:** Mix 400 mL of 10x transfer buffer with 2.8 L ddH_2_O and 600 mL MeOH. Add ddH_2_O to a final volume to 4 L. Filter the solution by 0.45-μm filter. Store at 4 °C for up to 6 months.

**! CRITICAL**: Do not add MeOH to 10x concentrated buffer to prevent precipitation.

- **TBS buffer, 5x:** Dissolve 303 g Tris base (2.5 M) and 219.2 g NaCl (3.75 M) in ddH_2_O to make 5 L buffer. Adjust the pH to 7.6 using pH meter. Store at room temperature for up to 6 months.
- **TBST buffer:** Add 400 mL 5x TBS buffer and 1 mL Tween-20 (0.1% v/v) in 1600 mL ddH_2_O. Stir it until fully homogeneous. Store at room temperature for up to 1 month.
- **Laemmli buffer (4×):** Dissolve 3 g Tris base (250 mM) and 8 g LDS (8% w/v) in 50 mL ddH_2_O. Adjust pH to 6.8. Then add 40 mL glycerol, 400 mg (0.4% w/v) bromophenol blue, and ddH_2_O until the final volume to 100 mL. Stir it until fully homogeneous. Store at room temperature for up to 6 months, and add 6% (v/v) β-mercaptoethanol (BME) immediately before use.
- **DNA loading dye (5×):** Dissolve 7.5 mL (15% v/v) glycerol and 50 mg Orange G in 50 ml ddH_2_O. Store at room temperature for up to 6 months.

### EQUIPMENT SETUP

#### Imaging plates

Dissolve 1 g agarose in 100 ml H_2_O. Microwave until agarose is fully dissolved and allow to cool until warm, but not solidified (i.e., 50-60 °C). Put 1-3 drops onto glass microscope slides. Cover the agarose drop with another glass slide, held by two other holding glass slides with tape on the surface, to make the agarose drop flat, with ∼1-2 cm^2^ size and ∼0.5 mm thickness. Leave 30 s – 1 min to solidify. The 1% agarose can be reused by microwaving.

> **CRITICAL** Use fresh imaging plates. Long exposure time (> 10 min) in air will cause the agarose pad to dry, leading to an increased agarose concentration and hardened texture. Wrinkles may also form on the surface. These changes interfere with imaging and affect worm morphology.

#### Leica M205 FA setup for worm imaging

Based on the fluorophore used, set the filters to matched channels for GFP, mCherry, or Cy5.

#### Platinum worm-pickers

Cut ∼2 cm length of platinum-iridium wire using scissors. Cut or melt the narrow tip of a glass pipette by heating it over a Bunsen burner flame until the glass begins to soften. While rotating the pipette in the flame, insert one end of the platinum wire into the molten glass tip and let it fuse. Remove it from the flame and cool down the pick at room temperature. Press and flatten the free platinum wire tip using lab spatula.

**! CAUTION**: Use tweezers to handle the platinum wire and glass pipette during the fusing process to prevent accidental burns.

#### Mass spectrometer

Connect the Dionex Ultimate 3000 RSLCnano system online to a Q Exactive HF Orbitrap mass spectrometer. Configure the instrument to acquire data in data-dependent acquisition (DDA) mode. Set full MS scans at a resolution of 120,000 at m/z 200, with a scan range of 350–1,800 m/z and an AGC target of 3 × 10⁶ ion counts. Select the 15 most intense precursor ions per cycle for fragmentation by Higher-energy Collision Dissociation (HCD) at a Normalized Collision Energy (NCE) of 32%, with an isolation window of 0.7 m/z. Acquire MS2 fragment ion spectra at a resolution of 30,000 at m/z 200, with a maximum ion injection time of 120 ms and an AGC target of 1 × 10⁵ ion counts. Enable dynamic exclusion for 40 s. Restrict precursor selection to charge states 2–6 and enable monoisotopic precursor determination. Set the fixed first mass to 100 m/z to ensure coverage of TMT reporter ions. Heat the analytical column to 50 °C using a column heater throughout the run.

> **CRITICAL** The MS2 resolution of 30,000 used here is sufficient for TMT6plex. Users adapting this protocol toward higher-plex TMT reagents (e.g., TMT 11-plex or TMTpro 16/18-plex) should consider increasing MS2 resolution to 45,000 or 60,000 to ensure adequate resolution.

#### Parameters for searching MS data

Process raw files with either MaxQuant or Proteome Discoverer against the UniProt reference proteome of the relevant organism (*C. elegans* or *H. sapiens*), supplemented with the relevant transgene sequence construct used in each experiment. Set trypsin/P as the digestion enzyme with a maximum of 2 missed cleavages. Specify carbamidomethylation of cysteine as a fixed modification, and oxidation of methionine, phosphorylation of serine/threonine/tyrosine, protein *N*-terminal acetylation, and glutamine-to-pyro-glutamate conversion at *N*-terminal glutamine as variable modifications. Enable TMT reporter ion quantification with isotope purity correction applied. Enable cross-run feature alignment (Match between runs in MaxQuant or equivalent retention time alignment in Proteome Discoverer). Apply a 1% FDR threshold at both the peptide-spectrum match and protein levels using the MaxQuant target-decoy approach.

#### Parameters for defining significant Localis-REX hits

The primary hit-defining method in this protocol is setting Gaussian threshold (μ + 2.5σ, Steps 166-167), which does not rely on volcano-plot statistics. For visualization and exploratory analyses, a volcano plot can be generated as a complement. Several alternative approaches are available depending on the analytical framework (see **Box 1**), and the number of replicates (minimally 3 independent biological replicates are necessary).

##### Box 2: Alternative statistical frameworks for hit definition

**Limma-based analysis.** For moderated statistics with low replicate numbers (*n* = 3), differential protein abundance can be assessed using the R/Bioconductor package limma, which applies an empirical Bayes method to stabilize variance estimates across proteins. *p-*values are corrected for multiple testing using the Benjamini–Hochberg procedure, with significant hits defined at FDR < 0.05. This approach is appropriate when comparing multiple experimental conditions simultaneously. (E.g., two control groups in ultraID datasets).

**Perseus-based volcano plot.** Volcano plots can be generated in Perseus using permutation-based FDR correction applied to the *t*-test statistics. A key parameter is s0, which controls the relative weighting of fold change versus *P*-value in the significance score. A starting value of s0 = 0.1 is commonly used for TMT-based enrichment data, though values of 0.5-1.0 may be more appropriate when replicate variance is high. Significance boundaries are drawn at a user-defined FDR threshold (typically 0.01 or 0.05).

**Fixed threshold.** Proteins may be considered significant based on a hard cutoff of log₂FC and *p*value (e.g., |log₂FC| > 1 and *p* < 0.05) without further multiple-testing correction. Note that this approach does not account for multiple testing across the proteome, which is not recommended for definitive hit calling.

## Procedure

### *C. elegans* culture and preparation

#### C. elegans culture media preparation – • TIMING 7 d (hands-on duration: approx. 2 h)

1| Grow OP50 by inoculating glycerol stock into 3 mL LB medium. Incubate overnight at 37 °C with shaking.

2| Transfer 1 mL overnight OP50 culture into a 200 mL flask containing 60 mL LB medium. Incubate again overnight at 37 °C with shaking (typically, the OD_600_ value is 1.5-2.5).

**▴ CRITICAL STEP**

Steps 1 and 2 can be done together with steps 3-5 to shorten the preparation time and keep the OP50 fresh.

3| Prepare media for plates:

• For MYOB plates: Dissolve 0.79 g Tris base (6.5 mM) and 2 g NaCl (34 mM) in 1 L ddH_2_O, adjust pH to 8.0. Then add 3 g tryptone and 17 g agarose. Sterilize by autoclave. Allow it to cool to 50 - 60 °C.

• For NGM plates: Mix 3 g NaCl (51 mM), 17 g agar, and 2.5 g peptone in 975 mL H_2_O. After autoclaving and cooling to 50 - 60 °C, add filter-sterilized 25 mL of 1 M potassium phosphate buffer (108.3 g KH_2_PO_4_, 35.6 g K_2_HPO_4_, H_2_O to 1 L, pH = 6.0), 1 mL of 1 M MgSO_4_, and 1 mL of 1 M CaCl_2_.

**▴ CRITICAL STEP**

Both NGM plates and MYOB plates can be used for general worm maintenance. However, it is essential to maintain consistent media conditions throughout the entire set of experiments. Wild-type worms exhibit normal survival and development on both MYOB and NGM plates, whereas certain mutant strains display distinct phenotypic differences between the two media. Consider the phosphate and pH sensitivities of your strain.

4| Add 1 mL of 5 mg/mL cholesterol (dissolved in EtOH) to 1 L media to achieve a final concentration of 5 µg/mL. Mix.

**▴ CRITICAL STEP**

Do not add cholesterol and extra salts until the autoclaved media is cooled (50 - 60 °C). Do not allow the autoclaved media to over-cool for proper plate pouring. Cholesterol is required for both MYOB and NGM plates.

5| Pour 15 mL of media into each sterile 10 cm dish, or 7.5 mL media into each sterile 6 cm dish. Let media solidify at room temperature.

6| Pipette 1.5 mL of OP50 culture onto the center of each solidified NGM plate (or MYOB) per 10 cm dish in a sterile biosafety cabinet. Pipette 0.5 mL of OP50 culture per 6 cm dish. Let the OP50 grow and dry completely at room temperature for 2-3 days.

**! CAUTION**

Avoid letting the OP50 lawn reach the plate edges to keep worms away from the walls and prevent them from drying out, especially for 6 cm dishes used in phenotypic screening and genetic crosses.

▴ CRITICAL STEP

The normal OP50 lawn should be semi-transparent, pale-yellow film on the surface of the NGM. Abnormal colony growth or dark yellow, orange, as well as unusually thick lawns are regarded as contamination. Any plate showing such abnormalities or growing mold should be discarded immediately to avoid cross-contamination.

**PAUSE POINT**

Store dried OP50-seeded plates in bags at 4 °C for up to 1 month until use.

**? TROUBLESHOOTING**

#### C. elegans stock recovery – • TIMING 2 d (hands-on duration: approx. 20 min)

7| Thaw cryopreserved *C. elegans* vials at room temperature. Gently mix the contents and pour the liquid in each vial onto a 10 cm OP50 plate. Incubate the plate at 20 °C in a temperature-controlled incubator.

▴ **CRITICAL STEP**

Actively moving worms should be observed after 1 hour under the microscope if the stock is of good quality. Clear moving tracks on OP50 lawn show successful worm recovery within the following days.

**? TROUBLESHOOTING**

8| Sterilize the platinum pick by flame and let it cool down for ∼ 5s. Pick 10 – 20 healthy L4-adult worms to other plates using platinum picks for subsequent culture and experiments.

▴ **CRITICAL STEP**

If contamination is observed during recovery, perform worm synchronization and cleaning before continuing (see steps 9-15).

**? TROUBLESHOOTING**

#### C. elegans synchronization and cleaning – • TIMING 2 d (hands-on duration: approx. 1 h)

9| Prepare *C. elegans* plates with a sufficient number (at least hundreds) of gravid adults, which can be obtained ∼4 days after picking 10 - 20 healthy L4-adult worms. Add 5 mL M9 buffer to each 10 cm plate. Use a glass pipette to gently wash worms off the plates and collect the suspension into a 15 mL Falcon tube.

10| Centrifuge the tube at 2000 × g for 1 minute to pellet the worms. Carefully aspirate the supernatant without disturbing the pellet.

11| Add 15 mL of freshly prepared 20% alkaline hypochlorite solution to the tube. Rotate the tube end-over-end gently for up to 5 minutes; check that most worm bodies are dissolved.

**! CAUTION**

Handle the alkaline sodium hypochlorite solution with care as it is corrosive.

▴ **CRITICAL STEP**

The incubation time should not exceed 5 minutes, even if some worm bodies remain undissolved, as prolonged exposure can significantly reduce egg viability.

12| Add M9 buffer to fill the tube to capacity. Centrifuge at 4000 × g for 1 minute to pellet the eggs.

13| Carefully aspirate the hypochlorite-containing supernatant without disturbing the egg pellet. Wash the egg pellet 3 times with 10 mL M9 buffer each time, centrifuging at 4000 × g for 1 minute between washes.

14| Resuspend the washed egg pellet in 7 mL M9 buffer. Incubate the suspension overnight (typically 16-24 hours) at room temperature with gentle end-to-end rotation to allow eggs to hatch into L1 larvae.

15| Distribute synchronized L1 larvae evenly onto 10 cm NGM plates with OP50.

▴ **CRITICAL STEP**

To control the number of L1 larvae per plate, place one drop of the suspension onto an NGM plate and examine under a microscope. A standard 10 cm NGM plate can support approximately 500 worms without reaching starvation before they reach adulthood.

**? TROUBLESHOOTING**

### *C. elegans* male induction, crossing, and selection

#### Male induction – • TIMING 2 w (hands-on duration: approx. 2 h)

**! CRITICAL**

Strains carrying fluorescent markers and showing no obvious abnormal phenotypes are recommended to serve as the male source. (e.g. strains with roller phenotype or reduced mobility have lower mating efficiency.) This will facilitate easier selection of cross progeny and ensure the overall efficiency of mating.

16| Pick 20 healthy L4 hermaphrodites of the desired strain onto 2-3 × 10 cm worm growth plates with OP50 using platinum picks. Incubate the plates at 31 °C for 6 hours to induce male formation.

▴ **CRITICAL STEP**

If the time is restricted, 37 °C incubation for 1 hour can be tried, but it usually causes significantly decreased fertility.

17| Transfer the plates to 20 °C. Culture for 4-5 days to obtain healthy L4 - adult worms.

▴ **CRITICAL STEP**

Besides the differences of tails, male worms are typically thinner and more active than hermaphrodites. These morphology differences become distinguishable from the L4 stage onward. Morphology details are available in WormAtlas.

18| Pick all males from these plates and transfer them to a 6 cm plate containing healthy L4 - young hermaphrodites of the same strain at a 3:1 to 4:1 male-to-hermaphrodite ratio. Repeat this step periodically to maintain the male population.

▴ **CRITICAL STEP**

Use plates where the OP50 lawn does not reach the plate edges to keep worms away from the walls and prevent them from drying out. Males are not self-sustaining in culture. The proportion of males on a mating plate declines with each generation as self-fertilized hermaphrodites only give 0.1-0.2% males. Re-induce males if necessary.

**? TROUBLESHOOTING**

#### C. elegans Crossing – • TIMING 2 w (hands-on duration: approx. 3 h)

19| Incubate the plates at 20 °C for 4-5 days. Pick healthy L4 - adult male worms onto 35 mm plates. Pick hermaphrodites from the desired strain and cross them with these males on 35 mm plates at a 3:1 to 4:1 male-to-hermaphrodite ratio. Culture the worms on crossing plates at 20 °C for 3 days.

▴ **CRITICAL STEP**

To increase the number of the males and enhance mating efficiency, males generated by mating are preferred. Males generated from heat shock are often less fertile.

20| Pick individual heterozygous F1 hermaphrodites at the L4 stage onto fresh 35 mm plates to allow self-fertilization (one animal per plate). Culture F1 worms at 20 °C.

**? TROUBLESHOOTING**

21| Pick 16 - 20 individual hermaphrodite F2 worms onto 35 mm plates (1 worm per plate). Culture F2 worms at 20 °C. Once sufficient F3 progeny are observed, remove F2 mothers and genotype them by single worm PCR (Steps 23-27).

22| Pick 11 individual F3 worms for single worm PCR validation (Steps 23-27) to confirm homozygosity of the target transgene or mutation from the F2 parent. If necessary, perform additional self-crossing steps until homozygosity is achieved.

#### Genotyping by single worm PCR – • TIMING 3 h

23| Pick a single adult worm into a PCR tube containing 10 μL worm digestion buffer (1× HF buffer, 0.5 mg/mL Proteinase K).

▴ **CRITICAL STEP**

Worms can be observed in the liquid under a microscope. Take care when picking to prevent the worm from sticking to the tube walls.

24| Spin the tube at 14,000 × g for 15 seconds to ensure the worm is submerged in the buffer. Flash-freeze the worm in liquid nitrogen for 10 minutes.

25| Incubate at 65 °C for 90 minutes to digest the worm, followed by 95 °C for 15 minutes to inactivate Proteinase K.

26| Use the digested lysate directly as the PCR template. Prepare PCR reactions (50 μL total volume): 2 μL DMSO, 0.2 μM forward and reverse primers (e.g. 295A-seq-s and 295A-seq-a), 0.2 mM dNTP mix, 0.04 U/μL Phusion polymerase, and 1× HF buffer. Run PCR cycles: Initial denaturation at 98 °C, 50 s; 30 cycles of: 98 °C, 10 s; 53 °C, 30 s; 72 °C, 45 s.

▴ **CRITICAL STEP**

Primer sequences should be designed based on the transgene sequences or the genomic regions flanking the mutations.

27| Analyze PCR products on a 1% agarose gel in NEB buffer. Clean PCR products using a PCR clean-up kit and perform Sanger sequencing if needed.

#### C. elegans stock preparation (optional) – • TIMING 3 d (hands-on duration: approx. 30 min)

**! CRITICAL**

Prepare frozen stocks as soon as a strain has been validated as homozygous or, for array-bearing strains, as soon as a well-transmitting line has been established. Restart working cultures from frozen stock rather than propagating indefinitely. Frozen stocks are recommended to be refreshed every half a year.

28| Prepare *C. elegans* plates with sufficient L1 and L2s (one 100 mm plate, or 2-3x 60 mm plates). Add 1.5 mL M9 buffer to each 10 cm plate. Use a glass pipette to gently wash worms off the plates and collect the suspension into a 1.8 mL cryovial. Add an equal volume of stock making buffer (S Buffer + 30% glycerol) to the cryovials. Mix gently by inversion.

▴ **CRITICAL STEP**

The L1 and L2 worms survive freezing the best of all age groups. To make stocks with good quality, freshly starved plates (12-24 hours without food) are ideal. Synchronized L1s are also suitable to make freezing stocks if it is necessary.

29| Place the cryovials into Mr. Frosty freezing container (recommended) or a small Styrofoam container to allow gradual freezing. Transfer the entire container to a −80 °C freezer.

30| The next day, move cryovials to their designated liquid nitrogen or −80 °C box for long-term storage.

**▪ PAUSE POINT**

Worm stocks can be stored at −80 °C or in liquid nitrogen for several years.

31| Thaw one vial and assess viability to ensure successful freezing.

▴ **CRITICAL STEP**

Motile animals should be visible within a few minutes. On the following day, moving traces on OP50 lawn should appear, and recovery from liquid nitrogen is typically 35-45%. Allow them to reproduce for one generation and confirm that the progeny carry the expected marker, especially for extrachromosomal-array strains.

**? TROUBLESHOOTING**

### OS-Localis-REX in *C. elegans*: Optimization of Ht-PreHNE(alkyne) treatment conditions (TMR-blocking) for OS-Localis-REX in *C. elegans*

**CRITICAL** This section should be performed for any newly generated Halo-expressing strain prior to main experiments. Optimal probe concentration may vary across strains due to differences in promoter activity, transgene copy number, mosaic expression of extrachromosomal arrays, and developmental stage-specific variations in promoter strength, protein turnover and synthesis rates, etc. The final Ht-PreHNE(alkyne) concentration should be determined based on the strains and nature of the comparative experiments being planned (typically < 32 μM to prevent off-target labeling).

Also see **Supplementary Movie 2** and **3**.

#### C. elegans culture – • TIMING 4.5 d (hands-on duration – picking per plate: approx. 15 min)

32| Pick 10 - 20 healthy L4-adult worms with desired Halo expression to worm growth plates with OP50 using platinum picks. Culture at 20 °C for 4-5 days to reach a densely populated but non-starved state on 10 cm plates.

**! CAUTION**

Check the plates every day to prevent starvation, which changes the *C. elegans* behavior and gene regulation significantly.

▴ **CRITICAL STEP**

Typically, 1 x 10 cm plate is used per sample for gel-based experiments. (e.g. One plate per specific Ht-PreHNE(alkyne) concentration as one replicate). The titration containing five conditions require 15 plates per strain for three independent biological replicates. The resulting population is mixed-age and mixed-stage as described. If stage-synchronization is required for specific experimental purpose, proceed with bleaching synchronization as described in Steps 9-15.

#### C. elegans collection – • TIMING 1.5 h

33| Add 3 mL M9 buffer to each 10 cm plate. Use a glass pipette to gently wash worms off the plates and collect the suspension into a 15 mL Falcon tube.

▴ **CRITICAL STEP**

On the day before the experiment, inoculate OP50 as described in Step 6 and grow to saturation overnight at 37 °C with shaking.

34| Centrifuge the tube at 3000 × g for 1 minute to pellet the worms. Carefully aspirate the supernatant without disturbing the pellet.

35| Wash the worm pellets with 10 mL S buffer each time, centrifuging at 3000 × g for 1 minute between washes. Repeat until the supernatant appears clear.

36| Resuspend the worm pellets in S medium. Adjust the total volume and evenly split into 1.5 mL Eppendorf tubes, each containing 0.5 mL.

▴ **CRITICAL STEP**

Note that the buffer is changed to S medium for the 6-h incubation step.

#### C. elegans treatment with Ht-PreHNE (alkyne) – • TIMING 9 h

▴ **CRITICAL**

Starting from step 38, all procedures need to be done under dim red light.

37| Pellet 10 mL saturated OP50 culture (Step 33) and resuspend in 2.5 mL S medium to make a 4x food stock. Add 250 μL 4x food stock solution each 1.5 mL Eppendorf tube.

38| Dilute Ht-PreHNE(alkyne) in S medium to make a 400 μM, 200 μM, 100 μM, and 50 μM (4×) stock solution. Vortex and mix the solution well. Add 250 μL 4× Ht-PreHNE(alkyne) stock to each 1.5 mL Eppendorf tube to reach a final concentration of 100 μM, 50 μM, 25 μM, and 12 μM. Add 250 μL of S medium containing the DMSO (vector control) for the control group.

**! CAUTION**

Ensure that the final DMSO concentration is identical across all 4× stock solutions. For the 200 μM, 100 μM, and 50 μM groups, add extra DMSO to match the DMSO content in the 400 μM stock solution.

39| Check and adjust the final volume in each 1.5 mL Eppendorf tube to 1 mL with additional S medium. Incubate the samples for 6 hours at room temperature under dim red light with gentle end-over-end rotation.

▴ **CRITICAL STEP**

A 6-hour incubation is an optimized condition for normal *C. elegans* samples. Incubation times may vary for other probes or strains after optimization. Small-molecule uptake in *C. elegans* occurs mainly through ingestion, with minimal contribution from cuticular diffusion. Special mutant strains, such as *bus-8*, exhibit increased cuticle permeability.

40| After the incubation, wash samples twice with 1 mL S buffer and a third time with 50 mM HEPES (pH 7.6). Each wash should be 30 minutes with gentle rotation. Centrifuge at 3000 × g for 1 minute between each wash. Aspirate the supernatant as much as possible. Flash-freeze in liquid nitrogen and store the samples at -80 °C.

**▪ PAUSE POINT**

Samples can be stored at −80 °C for up to 1 week without significant loss of signal.

#### C. elegans Lysis – • TIMING 3 h

**! CRITICAL**

Worm lysates must be used immediately for subsequent steps. Samples should be kept under red light for the whole process below.

41| Remove worm pellets from −80 °C storage and thaw on ice.

42| Prepare lysis buffer fresh on ice by mixing 4.48 mL basal HEPES buffer containing 1× Roche cOmplete™ mini protease inhibitor (EDTA-free, typically half pellet for 5 mL), 50 μL 100 mM TCEP stock solution, and 0.5 mL 10% Triton X-100 (in 50 mM HEPES, pH 7.6).

43| Add 2 to 8 times the pellet volume of freshly prepared, chilled lysis buffer to each worm pellet (e.g., for typical pellet volume of 10-20 μL, use ∼50-100 μL lysis buffer).

44| Add zirconia beads equal to 1/4 volume of the lysis buffer to each sample tube. Vortex the samples vigorously for 15 seconds, then flash-freeze the samples in liquid nitrogen and thaw them in a water bath at 20 °C. Repeat this vortexing/freezing/thawing cycle three times in total.

45| After three vortexing and freeze-thaw cycles, inspect the lysate under a standard stereomicroscope. If intact or partially disrupted worms remain visible, repeat the freeze-thaw/vortexing cycles and increase vortexing intensity or duration. Keep samples cold throughout to minimize protein degradation.

▴ **CRITICAL STEP**

If the lysate is difficult to inspect directly, briefly spin down the sample to collect worm debris and any intact animals at the bottom of the tube. This brief spin is only for inspection and should not replace the final clarification centrifugation in step 45.

46| Centrifuge lysates at 20,000 × g for 15 minutes at 4 °C. Carefully collect the supernatant containing soluble proteins.

▴ **CRITICAL STEP**

Sometimes flocculent precipitates appear after centrifugation. Prevent carrying them over when collecting the supernatant. These will lead to inaccurate protein quantification.

47| Perform Bradford assay in independent triplicates: Mix 2 μL lysate with 1 mL Bradford reagent. Pipette 200 μL of this mixture into wells of a 96-well plate. Measure absorbance at 595 nm using a plate reader.

**? TROUBLESHOOTING**

48| Calculate protein concentration by comparison with a BSA standard curve. Use the mean value of triplicate measurements.

49| Normalize protein concentration to 1 mg/mL.

#### TMR labeling and analysis – • TIMING 3 h

50| Take 30 μL of each sample and add Halo-TMR to a final concentration of 10 μM. Incubated the tubes at 37 °C for 30 min.

51| Add 10 μL 4× Laemmli dye containing 6% β-mercaptoethanol (BME) to each sample. The samples can be stored at −20 °C overnight or analyzed immediately by SDS-PAGE. For quantification and validation of halo expression, transfer proteins to a PVDF membrane using standard protocols and perform western blot analysis.

▴ **CRITICAL STEP**

The data will show TMR-labeling of Halo protein, the signal of which should be maximum in the set where no photocaged probe [Ht-PreHNE(alkyne)] has been added. Signal should decrease progressively as a function of probe concentration. At the concentration of probe where TMR signal becomes negligible, probe occupancy on Halo in the worms is considered functionally saturated. This is defined as optimal Ht-PreHNE(alkyne) concentration in Localis-REX (**Fig. 6a**). For experiments including Halo expression across strains, it is recommended to perform a parallel western blot using an anti-Halo antibody to directly assess relative Halo expression levels across samples (**Fig. 6b**).

**Figure 6.**
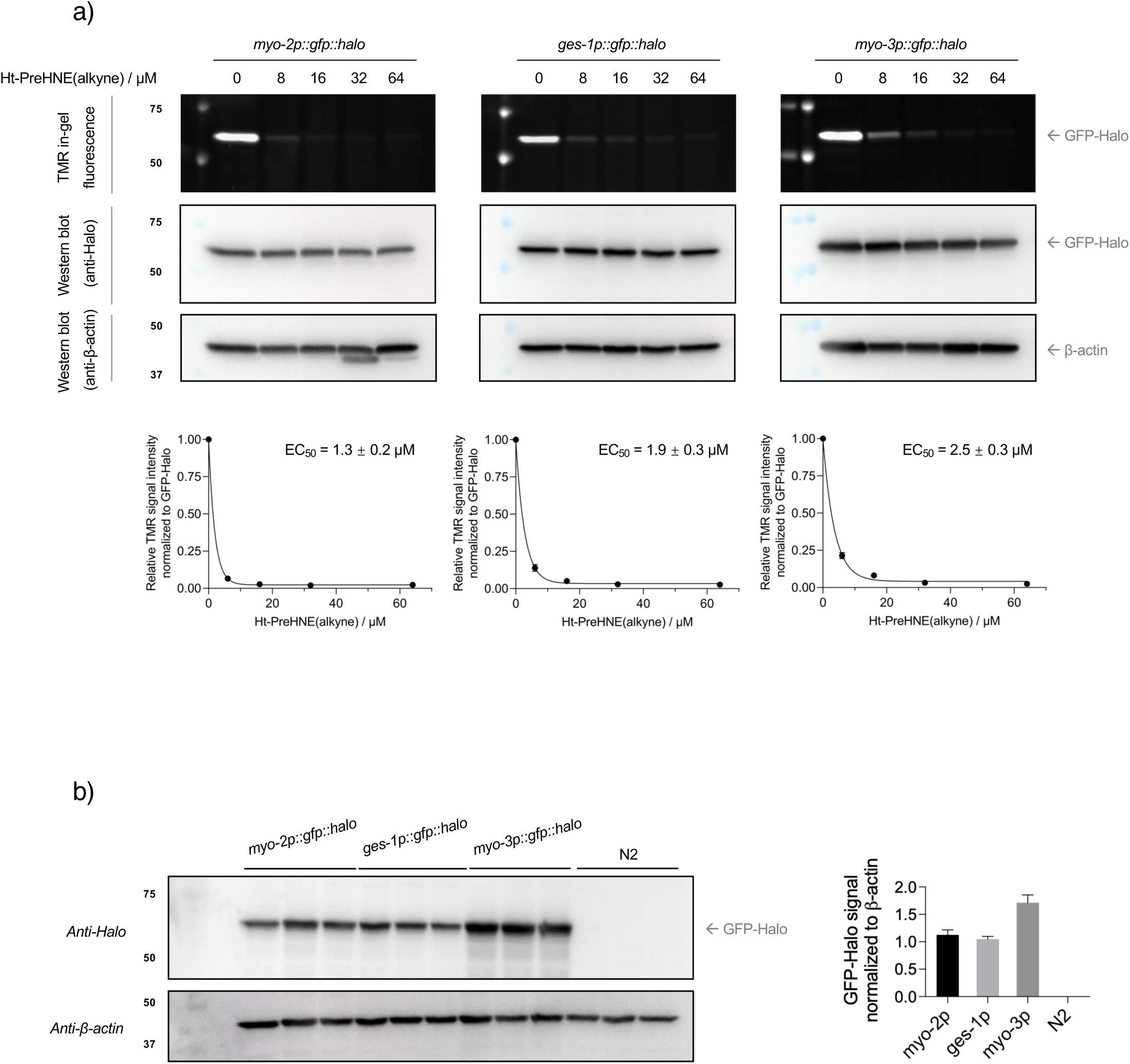
Probe saturation and transgene expression in tissue-specific *C. elegans* strains. **(a)** The functionality of single turn-over enzyme Halo in live transgenic Halo *C. elegans* was assessed by probe saturation of Halo binding sites, leveraging competitive binding assays using TMR-Halo ligand. Live worms were treated with gradient concentration of Ht-PreHNE(alkyne) probe (0 – 64 µM) for 6 hours, followed by worm lysis, and then incubation with 10 µM TMR-Halo ligand for 30 min, at 37 °C. Remaining unoccupied functional Halo sites were visualized by SDS-PAGE and in-gel fluorescence. Representative data are shown from *n* = 3 independent biological replicates. EC_50_^[Ht-PreHNE(alkyne) for saturating GFP-Halo *in vivo*]^ = 1.3 ± 0.2 μM for *myo-2p::gfp::halo* strain (**left**); 1.9 ± 0.3 μM for *ges-1p::gfp::halo* strain (**middle**); 2.5 ± 0.3 μM for *myo-3p::gfp::halo* strain (**right**), with the quantification and normalization based on western blot signals. All data present mean ± SEM. This figure is reproduced from previously published data in reference^11^. **(b)** Transgene expression across tissue-specific strains assessed by western blot analysis. **Left**: Normalized lysates from three *C. elegans* transgenic lines (*myo-2p::gfp::halo*, *ges-1p::gfp::halo*, *myo-3p::gfp::halo*) were blotted. Bristol N2 served as negative control. **Right**: GFP-Halo band intensities normalized to β-actin. *n =* 3 for all conditions. All data present mean ± SEM. This figure is reproduced from previously published data in reference^11^.

#### Localis-REX Precision localized electrophile generation in tissue-specific Halo transgenic *C. elegans*

#### C. elegans culture – • TIMING 4.5 d (hands-on duration – picking per plate: approx. 15 min)

52| The worm preparation is described in TMR-blocking steps. Typically, 1 x 10 cm plate is used per sample for gel-based experiments and imaging, while 6 x 10 cm plates are used per sample for mass spectrometry analysis. For large-scale experiments, use 15 mL Falcon tube during the incubation and biotin pulldown steps if needed. When using Falcon tubes, the reagent volumes should be scaled up to ten times those used in 1.5-mL Eppendorf tubes. Falcon tubes are usually for 3-6x 10 cm plates. The following protocol specifies reagent amounts for 15 mL Falcon tubes.

#### C. elegans collection – • TIMING 1.5 h

53| Perform the worm collection steps described above (Steps 33-35).

54| Resuspend worm pellets in S medium, pool the suspensions from the same experimental setting, and evenly split into new 15 mL Falcon tubes as 1 mL each.

▴ **CRITICAL STEP**

The buffer is changed to S medium for the 6-h incubation step.

#### C. elegans treatment with Ht-PreHNE (alkyne or no alkyne) – • TIMING 9 h

**▴ CRITICAL**

Starting from step 55, all procedures need to be done under dim red light.

55| Pellet 50 mL saturated OP50 culture (Step 33) and resuspend in 12.5 mL S medium to make a 4x food stock. Add 2.5 mL 4x food stock solution to each 15 mL Falcon tube. For 1.5 mL Eppendorf tubes, start with 500 μL worm suspension and add 250 μL 4x food stock.

56| Dilute Ht-PreHNE(alkyne) in S medium to make a 120 μM (10×) stock solution. Vortex and mix the solution well. Add 1 mL of 10× Ht-PreHNE(alkyne) stock to each Falcon tube, or 100 μL to each 1.5 mL Eppendorf tube to reach a final concentration of 12 μM. For the no-alkyne control group, add the same volume of S medium containing 120 μM (10×) Ht-PreHNE(no alkyne): 1 mL per Falcon tube or 100 μL per Eppendorf tube.

▴ **CRITICAL STEP**

The Ht-PreHNE concentration (12 μM) is optimized specifically for the *myo-2p::gfp::halo*, *ges-1p::gfp::halo*, and *myo-3p::gfp::halo* strains. For novel generated strains, different concentrations should be tested, obeying the same procedures and tested by the TMR-blocking assay. (Described in the TMR-blocking section, Steps 32-51)

57| Adjust the final volume in each Falcon tube to 10 mL, or each 1.5 mL Eppendorf tube to 1 mL with additional S medium. Incubate the samples for 6 hours at room temperature under dim red light with gentle end-over-end rotation.

58| After the incubation, wash samples twice with 10 mL S buffer and a third time with 50 mM HEPES (pH 7.6) for samples in Falcon tubes (or 1 mL buffers for samples in Eppendorf tubes). Each wash should be 30 minutes with gentle rotation. Centrifuge at 3000 × g for 1 minute between each wash. Discard most of the supernatant and leave 1 mL 50 mM HEPES with suspended worms.

▴ **CRITICAL STEP**

The final 1 mL volume is for both Eppendorf samples and Falcon tube samples.

▴ **CRITICAL STEP**

Following step 58, Click and worm imaging steps may be performed without freezing the sample. (steps 61-72, **Fig. 2a**)

59| Transfer washed worms into the center of a 6-well plate using a glass pipette for UV photoactivation. Expose washed worms to UV light (365 nm, 5 mW/cm²) for 5 minutes. If a ‘non-light’ group were designed, samples in this group should not be illuminated by UV.

**! CAUTION**

Wear UV-protective goggles if operating the lamp with an unshielded beam path.

▴ **CRITICAL STEP**

5-min illumination is optimized for normal *C. elegans* samples. Adjust and test different illumination durations if needed.

60| Transfer each group of worms into 1.5 mL Eppendorf tubes using a glass pipette. Rinse the remaining well contents with 0.5 mL of 50 mM HEPES (pH 7.6) to recover residual worms and combine with the main sample. Pellet worms by brief centrifugation, typically at 3000 x g for 1 minute. Aspirate as much of the supernatant as possible. Flash-freeze the pellet in liquid nitrogen and store at -80 °C for the subsequent Worm lysis and Click (Steps 73-80).

**▪ PAUSE POINT**

Pellets can be stored at −80 °C for up to 1 week without significant loss of signal.

*Click chemistry and C. elegans imaging –* • TIMING 3 d (hands-on duration: approx. 3 h)

61| (Continue from step 58 with the unfrozen samples intended for imaging) Add 1 mL of 4% paraformaldehyde to fix the worms and incubate overnight at 4 °C with gentle end-to-end rotation.

**! CRITICAL STEP**

This step disrupts the native structure of the pharynx, causing it to twist and shrink. If preservation of the pharynx is critical, particularly for *myo-2p* strains, omit this step and proceed with adding cold methanol (step 62).

62| Remove 4% paraformaldehyde carefully and incubate worms in 1 mL of cold methanol (−20 °C) for 24 hours.

63| Remove the methanol. Wash worms twice with PBS + 0.015% Tween-20 (30 minutes each), followed by two washes with 50 mM HEPES, 150 mM NaCl, pH 7.6.

64| Prepare 1× Click mixture containing final concentrations of 1% SDS, 5% t-BuOH, 1 mM CuSO_4_, 0.1 mM Cu(II)TBTA, 2 mM TCEP, and 10 μM Cy5-azide.

**! CRITICAL STEP**

Add TCEP stock solution last, to initiate reaction just before adding to worms.

65| Incubate worms in the Click mixture at room temperature in a thermomixer at 300 rpm for 1 hour. 66| Wash worms 3 times in PBS + 0.015% Tween-20, then incubate overnight at 4 °C with end-to-end rotation.

67| Wash worms 2 × 30 min with 1 mL PDT buffer (PBS, 0.1% Tween-20, 1% DMSO) at room temperature.

68| Block in 1 mL blocking buffer (PBS, 0.05% Tween-20, 10% FBS, 2% BSA) for 1 hour at room temperature.

69| Incubate worms with goat anti-GFP diluted 1:500 in blocking buffer, overnight at 4 °C.

70| Wash worms 2 × 30 min in PDT buffer, then re-block in fresh blocking buffer for 1 hour at room temperature.

71| Incubate with Alexa Fluor 488-conjugated donkey anti-goat IgG diluted 1:500 in blocking buffer for 1.5 hours at room temperature.

72| Wash 3 times in PDT buffer (1 mL each, 30 minutes per wash). Mount worms on 1% agarose drop and image using Leica M205 FA microscope under stereo and 5× HR objectives.

**? TROUBLESHOOTING**

#### C. elegans Lysis – • TIMING 3 h

**! CRITICAL**

Worm lysates must be used immediately for subsequent steps.

73| Perform the worm lysis steps as described in steps (41-49) and normalize protein concentration to 1-2.5 mg/mL. For samples from 6 x 10 cm plates, the typical pellet volume is 60 - 100 μL, which is lysed using ∼400 μL lysis buffer.

▴ **CRITICAL STEP**

The required protein concentration depends on downstream experiments. For in-gel fluorescence or western blot analysis, a concentration of 1 mg/mL with 40 μL lysate per sample is sufficient. For biotin pulldown or mass spectrometry-based experiments, a concentration of 2.5 mg/mL in a total volume of 1 mL is recommended.

#### Preclearance – • TIMING 2.5 h

**CRITICAL** When preparing lysates for biotin-pulldown, pre-clearance steps (Steps 74-76) are essential for

*C. elegans* lysates prior to the Click reaction, as this organism contains several endogenously biotinylated proteins. Failure to remove these background proteins can significantly interfere with enrichment of HNE(alkyne)-modified proteins, reducing specificity and giving a high background signal.

74| Calculate the total amount of streptavidin beads needed (typically, 50 μL bed volume per 1-2.5 mg of protein in lysate, equivalent to 100 µL of a 50% v/v slurry). Pre-rinse the calculated total volume of beads (e.g., 100 μL) by rotating them for 5 minutes in 1 mL ddH₂O, and then twice in 1 mL of 0.5% LDS in 50 mM HEPES (pH 7.6). Between each rinse, pellet beads by centrifugation at 500 × g for 1 min, then remove the supernatant.

75| Aliquot 50 μL bed volume of the pre-rinsed streptavidin beads into each sample. Incubate for 2 hours at room temperature with end-to-end rotation.

▴ **CRITICAL STEP**

Step 75 can be changed to a two-step sequential pre-clearance. Incubate the lysate with the first batch of 25 μL pre-rinsed beads for 1 hour, centrifuge to collect the supernatant, and transfer it to a second batch of 25 μL pre-rinsed beads for another 1 hour. This strategy maximizes the depletion of endogenous biotinylated proteins.

76| Centrifuge the mixture at 2000 × g for 1 minute and carefully collect the supernatant for the Click reaction.

#### Click chemistry – • TIMING 1 h

77| Prepare 5× Click mixture containing final concentrations of 5% SDS, 25% t-BuOH, 5 mM CuSO_4_, 0.5 mM Cu(II)TBTA, 1 mM biotin azide, and 10 mM TCEP. Add TCEP stock solution last, to initiate reaction just before reaction.

▴ **CRITICAL STEP**

For reference, 1.25mL of 5x Click mixture for 6.25 mL total volume was prepared by mixing 313 μL 20% SDS, 313 μL t-BuOH, 62.5 μL 100 mM CuSO_4_, 62.5 μL 10 mM Cu(II)TBTA in 55% DMSO, 361.5 μL 50 mM HEPES (pH = 7.6), 12.5 μL 100 mM biotin azide, and 125 μL of 100 mM TCEP.

▴ **CRITICAL STEP**

The biotin azide needs to be changed to Cy5-azide for in-gel fluorescence. For Cy5-Click, pre-clearance steps are not necessary.

78| Add 250 μL of 5× Click mixture to 1000 μL precleared lysate (2.5 mg/mL protein) to reach final reaction concentrations of 1% SDS, 5% t-BuOH, 1 mM CuSO_4_, 0.1 mM Cu(II)TBTA, 2 mM TCEP, and 200 μM biotin azide.

79| Vortex briefly (a few seconds) and incubate at 37 °C for 30 minutes.

80| Add 4 volumes of pre-chilled (−20 °C) pure ethanol to each lysate. Mix and store samples at −80 °C for overnight to 1 week to precipitate proteins.

**▪ PAUSE POINT**

Samples can be stored at −80 °C for up to 1 week without significant loss of signal.

#### Biotin pulldown for enrichment of tissue-specific electrophile-responder proteins – • TIMING 8 h

81| Remove the samples from −80 °C freezer. Centrifuge the precipitated samples at 5,000 or 20,000 × g for 30 minutes at 4 °C.

▴ **CRITICAL STEP**

For large-amount samples in Falcon tubes, use 5000 x g. For small-amount samples in Eppendorf tubes, use 20,000 x g.

82| Discard the supernatant and wash pellets twice with 1 mL ice-cold 70% ethanol, and then once with acetone. Vortex the mixture for 30 seconds each time. Centrifuge the resuspended protein pellets at 5,000 x or 20,000 × g for 10 minutes at 4 °C between each wash.

83| Remove acetone and allow pellets to air-dry until residual acetone evaporates.

**! CRITICAL STEP**

Do not allow the pellet to dry completely, as this may make it difficult to fully resolubilize. Depending on the residual volume, evaporation typically takes 1–10 minutes.

84| Add 80 μL resuspension buffer (8% LDS and 1 mM Na_2_EDTA in 50 mM HEPES, pH 7.6) to each pellet. Vortex and sonicate pellets using a Bioruptor® Pico sonication device for 10 minutes.

85| Repeat sonication and vortex cycles until pellets are fully solubilized.

**! CRITICAL STEP**

Avoid excessive heating of the sonication bath during repeated cycles, as elevated temperatures may destroy the samples.

▴ **CRITICAL STEP**

Adding an appropriate volume of 50 mM HEPES buffer (while maintaining LDS concentration > 1%) may facilitate complete resolubilization during repeated cycles.

86| Dilute samples with 50 mM HEPES (pH 7.6) to a final LDS concentration of 0.5%. Centrifuge at 20,000 × g for 10 minutes.

87| Remove 30 μL supernatant as input sample; mix with 10 μL 4× Laemmli buffer. Store the samples at room temperature as they will be analyzed together with the elution samples the same day. Transfer the remaining supernatant to other tubes for the following biotin-streptavidin enrichment.

**! CRITICAL STEP**

Carefully avoid transferring any insoluble material or debris, as contamination with residual streptavidin beads will significantly compromise the accuracy of downstream enrichment results.

88| Calculate the total amount of streptavidin beads needed (typically, 50 μL bed volume per 1-2.5 mg of protein in lysate). Pre-rinse the calculated total volume of beads (e.g., 50 μL) by rotating them for 5 minutes in 0.5 mL ddH₂O, and then twice in 0.5 mL of 0.5% LDS in 50 mM HEPES (pH 7.6). Between each rinse, pellet beads by centrifugation at 500 × g for 1 min, then remove the supernatant.

89| Add the remaining supernatant to the aliquoted pre-rinsed streptavidin beads. Incubate for 2 hours at room temperature with an end-to-end rotation.

90| Centrifuge the mixture at 2000 × g for 1 minute and carefully collect 30 μL of the supernatant, mix with 10 μL 4× Laemmli dye, and store as the flow-through sample.

91| Wash the beads three times with 1 mL of 0.5% LDS in 50 mM HEPES (pH 7.6), each for 30 minutes at room temperature with gentle rotation. Between each wash, centrifuge at 500 × g for 5 minutes and remove the supernatant.

▴ **CRITICAL STEP**

Aspirate the supernatant as thoroughly as possible to maximize the purity of the elution samples. 92| Add 30 μL of 2× Laemmli dye with 3% β-mercaptoethanol (BME) directly to the beads. Gently tap the tube to mix the dye with the beads.

93| Boil the samples at 98 °C for 10 minutes, then centrifuge at 20,000 × g for 5 minutes. Collect the supernatant as the eluted sample for downstream analysis. For analysis by SDS-PAGE and Western blot, proceed to Step 94. Alternatively, to analyse samples by mass spectrometry, proceed to step 153. For mass spectrometry sample preparation, check Mass spectrometry sample preparation in Localis-REX experiments (Steps 153-162). QC (quality control) of mass-spec samples can be performed either by Coomassie staining (Step 94, **Fig. 7a**) or western blot (Step 95, **Fig. 7b**).

**Figure 7.**
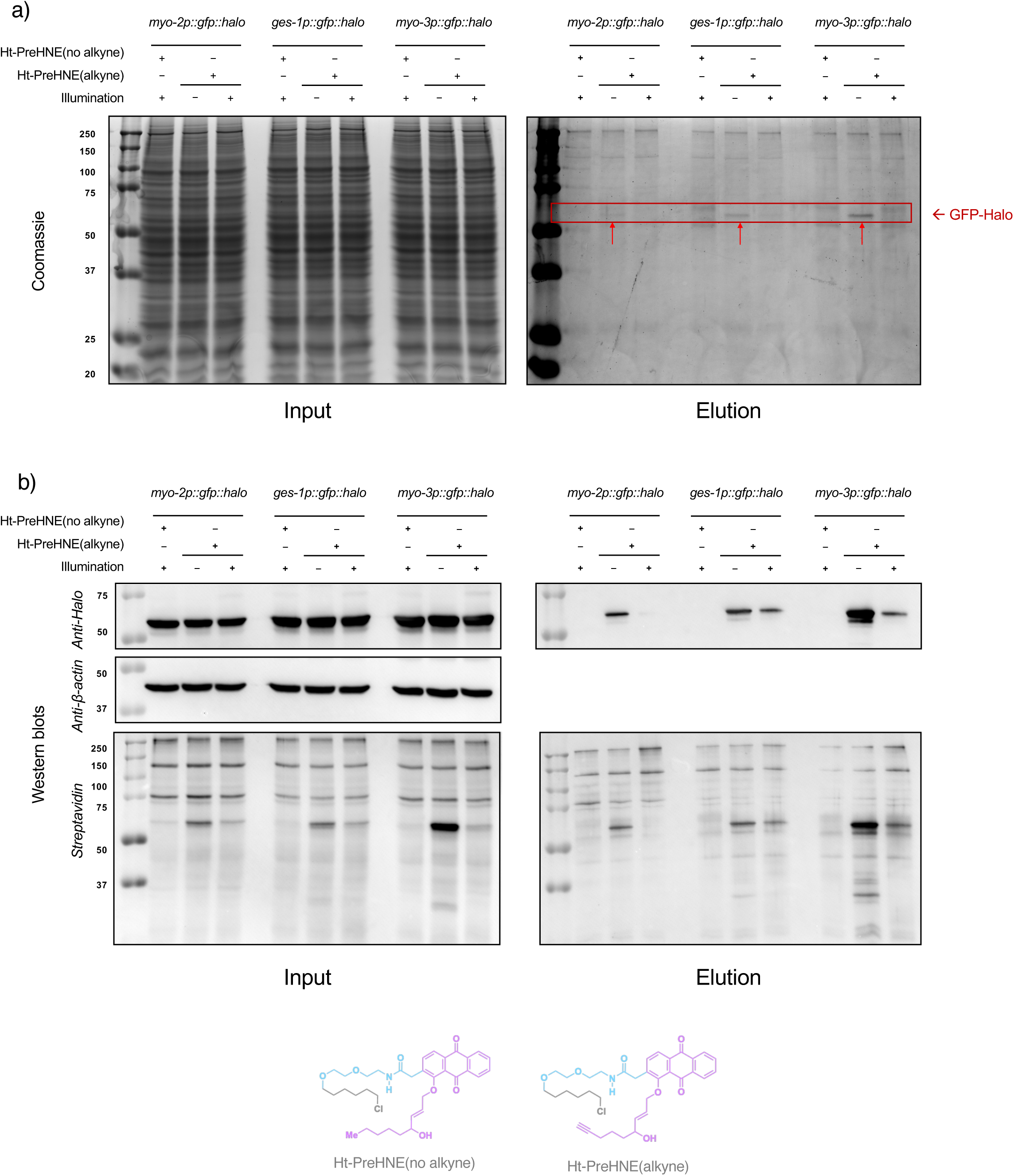
Validation of REM-responder enrichment via Click-biotin pulldown. Representative Coomassie-stained input gels **(a)** and western blots **(b)** show homogeneous loading. Coomassie-stained elution gels show selective enrichment of REM-responders (covalently adducted with alkyne-functionalized REM) in light-exposed, Ht-PreHNE(alkyne) probe [see structure in **(b**), bottom] treated group, with respect to two sets of control groups: (i) Ht-PreHNE(no-alkyne) probe treatment, and (ii) no light exposure. The extent of enriched GFP-Halo (∼60 kDa) decreases as expected in light-exposure group in both western blots and Coomassie-stained gels, whereas endogenously biotinylated proteins (∼130 kDa and ∼80 kDa bands, corresponding to the molecular weight of PYC-1 and PCCA-1) show homogeneous signals across all three conditions in streptavidin blots. **Bottom:** Chemical structures of the no alkyne and alkyne-integrated variants of Localis-REX photocaged-REM probes, wherein REM corresponds to HNE. Also see Fig. 1c.

**▪ PAUSE POINT**

Samples in Laemmli dye can be stored at −20 °C overnight.

**? TROUBLESHOOTING**

#### Enriched protein validation by SDS-PAGE and western blot – • TIMING 2 d (hands-on duration: approx. 3 h)

94| Load equal volumes of each sample (e.g., 10–20 μL) onto a 11% SDS-PAGE gel alongside a pre-stained protein ladder. Run electrophoresis at 110 V for 10 minutes, and then 170 V for 60 - 70 minutes in 1x running buffer until the dye front reaches the bottom.

95| Transfer proteins to a PVDF membrane using standard protocols and perform western blot analysis.

**? TROUBLESHOOTING**

### Localis-REX in cultured mammalian cells

**CRITICAL** Two mammalian cell lines are used, human HEK293T cells, and mouse differentiated N2A cells.

#### Cell culture and preparation – • TIMING 2 d (hands-on duration: approx. 1 h)

**CRITICAL** For SILAC experiments, replace these steps with Steps 99-102.

96| Seed HEK293T or N2A cells in 60 mm dishes in 5 mL of the complete medium (MEM, GlutaMAX for HEK293T, and DMEM for N2A, both are supplemented with 1X non-essential amino acids, 1 mM Sodium Pyruvate, 100 U/mL Penicillin-Streptomycin, and 10% v/v FBS), at 37 °C in a humidified 5% CO₂ incubator, and allow them to reach 70-80% confluence at the point of transfection.

97| Mix 5 µg of pCS2+8 NLS-Halo, pCS2+8 NES-Halo pCS2+8 MOMLS-Halo, or pCS2+8 ERt-Halo plasmid in 500 µL Opti-MEM, then add 20 µL of 1 mg/ml PEI stock solution gently. Incubate the mixture at room temperature for 30 min (HEK293T) or 20 min (N2A), then add it slowly to the cells.

98| 8 hours later, aspirate the old medium and replace it with fresh complete medium for HEK293T cells, or DMEM, supplemented with 1X non-essential amino acids, 1 mM Sodium Pyruvate, 100 U/mL Penicillin-Streptomycin, 2% v/v FBS, and 20 µM retinoic acid to differentiate N2A cells. Incubate the cells for 28 more hours.

**? TROUBLESHOOTING**

#### (Optional) Cell culture and preparation for SILAC experiments– • TIMING 2 d (hands-on duration: approx. 1 h)

**CRITICAL** For SILAC experiments, these steps should replace Steps 96-98.

99| Grow HEK293T Light and Heavy cells in light and heavy complete media, respectively.

100|After four passages, collect a small portion of cells for each following passage, and test the incorporation efficiency of isotopic amino acids by MS proteomics of soluble lysates, following previously published procedures^12, 50^.

101|If the incorporation efficiency of isotopic amino acids reaches 98%^12, 50^, make cell stock and seed cells in a 60 mm dish for Localis-REX, and allow them to reach 70-80% confluence at the point of transfection. Otherwise, culture for more passages.

102|Mix 5 µg of pCS2+8 NLS-Halo plasmid in 500 µL Opti-MEM, then add 20 µL of 1 mg/ml PEI stock solution gently. Incubate the mixture at room temperature for 30 min, then add it slowly to the Light cells. Same procedure for transfecting Heavy cells with pCS2+8 MOMLS-Halo. Incubate the cells for 36 hours.

**? TROUBLESHOOTING**

#### Cell treatment with Ht-PreHNE (alkyne or no alkyne) – • TIMING 5 h

**CRITICAL** For SILAC experiments, replace these steps with Steps 107-109.

▴ **CRITICAL**

Starting from step 103, all steps need to be carried out under dim red light.

103|Aspirate the old medium and incubate cells with 5 mL of rinsing medium (MEM with 1X NEAA, 1X Pyruvate and 1X Pen/Strep) containing 10 µM Ht-PreHNE(alkyne) for 2 hours.

104|Rinse cells three times with 5 mL of fresh rinsing medium, incubate cells for 30 min during each rinse to fully remove unbound probe.

**! CRITICAL STEP**

Be extremely careful and gentle during multiple rinsing to avoid detaching cells.

105|Under Localis-REX condition, expose cells to 365 nm light (5 mW/cm^2^) for 5 min; Under light-alone control, treat cells with DMSO instead of Ht-PreHNE(alkyne) and expose to 365 nm light for 5 min; Under Probe-alone condition, treat cells with 10 µM Ht-PreHNE(alkyne) for 2 h, but no light irradiation; Under DMSO control, treat cells with DMSO instead of Ht-PreHNE(alkyne) and no light irradiation.

**! CAUTION**

Wear UV-protective goggles, should the lamp be in operation with an unshielded beam path.

106|Harvest cells from all conditions and wash cells twice with cold DPBS. Snap-freeze cells in liquid nitrogen for downstream analysis (Step 110).

**! CRITICAL STEP**

Harvest at this step only for responder identification by proteomics. The live-cell readouts of downstream signalling do not continue from the frozen pellet, such as Steps 136-141 for the translation efficiency of a single protein, Steps 142 for proteome-wide ncAA incorporation, and Steps 143-152 for BONCAT.

**▪ PAUSE POINT**

Samples can be stored at −80 °C for up to 1 week without significant loss of signal.

#### (Optional) Cell treatment for SILAC experiments– • TIMING 2 d (hands-on duration: approx. 3 h)

**CRITICAL** For SILAC experiments, these steps should replace Steps 103-106.

107|Aspirate the old medium and incubate cells with rinsing medium (SILAC Light or Heavy media without FBS) containing 10 µM Ht-PreHNE (alkyne) for 2 hours.

108|Expose both Light cells and Heavy cells to 365 nm light (5 mW/cm^2^) for 5 min.

**! CAUTION**

Wear UV-protective goggles, should the lamp be in operation with an unshielded beam path.

109|Harvest both Light cells and Heavy cells from respective conditions. Mix the Light and Heavy cells in a 1:1 ratio based on the cell number. Snap-freeze cells in liquid nitrogen for downstream analysis. For mass spectrometry sample preparation, check Mass spectrometry sample preparation in Localis-REX experiments (Steps 152-168).

**▪ PAUSE POINT**

Samples can be stored at −80 °C for up to 1 week without significant loss of signal.

**? TROUBLESHOOTING**

#### Cell lysis – • TIMING 2 h

▴ **CRITICAL**

All procedures need to be done under dim red light.

110|Remove cell pellets from −80 °C storage and thaw on ice.

111|Prepare lysis buffer fresh on ice by mixing 4.48 mL basal HEPES buffer containing 1× Roche cOmplete™ mini protease inhibitor (EDTA-free, typically half pellet for 5 mL), 50 μL 100 mM TCEP stock solution, and 0.5 mL 10% Triton X-100 (in 50 mM HEPES, pH 7.6).

112|Add an appropriate amount of chilled lysis buffer to each cell pellet (e.g., 300-400 μL lysis buffer for a 60 mm plate).

113|Perform three cycles of freeze-thaw: freeze samples in liquid nitrogen, then thaw in a water bath at 20 °C.

114|Centrifuge lysates at 20,000 × g for 15 minutes at 4 °C. Carefully collect the supernatant containing soluble proteins.

115|Perform Bradford assay in independent triplicates: Mix 2 μL lysate with 1 mL Bradford reagent. Pipette 200 μL of this mixture into wells of a 96-well plate. Measure absorbance at 595 nm using a plate reader.

116|Calculate protein concentration by comparison with a BSA standard. Use the mean value of triplicate measurements.

117|Normalize protein concentration to 1 mg/mL.

#### Click reaction and streptavidin pull-down – • TIMING 5 h

118|The Click reaction and streptavidin pull-down part is described in the OS-localis-REX in *C. elegans* (Steps 77-95). Only omit the pre-clearance steps. Collect the input and elution samples. For mass spectrometry samples preparation, check mass spectrometry sample preparation in Localis-REX experiments (Steps 152-168).

**▪ PAUSE POINT**

Samples in Laemmli dye can be stored at −20 °C overnight.

**? TROUBLESHOOTING**

### Immunofluorescence imaging to validate expected localization of Ht-PreHNE(alkyne) at designated subcellular locales

#### Cell culture and preparation – • TIMING 2 d (hands-on duration: approx. 1 h)

119|Seed N2A or HEK293T cells into 35-mm glass-bottom dishes and allow cells to grow for 24 hours until they reach ∼40% confluence.

120|Transfect cells with 2µg of one of the following plasmids: NLS-Halo, NES-Halo, MOMLS-Halo, or ERt-Halo, using lipofectamine 3000 reagent according to the manufacturer’s instructions.

121|After 5 h, change to differentiation medium (DMEM with 2% FBS and 20 µM retinoic acid) and incubate for 31 hours.

▴ **CRITICAL STEP**

Incubate cells for a minimum of 24 hours to initiate differentiation.

122|Treat cells with 15 µM Ht-PreHNE(alkyne) for 2 hours in rinsing medium (DMEM without FBS). 123|Wash cells three times with rinsing medium to remove unbound probe, 30 min incubation during each time.

124|For the cells ectopically expressing MOMLS-Halo, add 100 nM MitoTracker Red in rinsing medium and incubate for 30 minutes

125|Wash cells twice with DPBS then add 2 mL prechilled methanol to each dish and incubate cells at −20 °C for 20 minutes.

126|Wash cells three times with DPBS.

**▪ PAUSE POINT**

The cell samples can be stored at 4 °C for one week.

127|Block cells using DPBS containing 3% BSA and 0.2% Triton X-100 for 1 hour at room temperature. 128|Prepare Click buffer containing: 1.1 mM CuSO₄, 10 mM sodium ascorbate, 15 µM Cy5-azide in 50 mM HEPES buffer (pH 7.6) and incubate cells in the Click buffer for 30 minutes at room temperature.

129|Wash cells three times with DPBS, 5 minutes incubation for each wash.

130|Incubate cells with anti-Halo rabbit polyclonal antibody (1:1000) in DPBS containing 1% BSA and 0.02% Triton X-100 for 2 hours at room temperature. For cells ectopically expressing ERt-Halo, incubate cells with anti-Halo rabbit polyclonal antibody (1:1000) and anti-Calnexin mouse monoclonal antibody (1:200) in DPBS containing 1% BSA and 0.02% Triton X-100 for 2 hours at room temperature.

▴ **CRITICAL STEP**

Two different primary antibodies (such as anti-Halo rabbit polyclonal antibody and anti-Calnexin mouse monoclonal antibody) can be incubated at the same time with corresponding dilution conditions.

131|Wash cells three times with DPBS, 5 minutes incubation for each wash.

132|Incubate cells with secondary antibody Alexa Fluor 488 anti-rabbit IgG (1:2000) in DPBS containing 1% BSA and 0.02% Triton X-100 for 1 hour at room temperature. For cells ectopically expressing ERt-Halo, incubate cells with secondary antibody Alexa Fluor 488 anti-rabbit IgG (1:2000) and Alexa Fluor 568 anti-mouse IgG (1:2000) in DPBS containing 1% BSA and 0.02% Triton X-100 for 1 hour at room temperature.

**! CRITICAL STEP**

Incubation with secondary antibodies needs to be performed in the dark.

133|Wash cells three times with DPBS, a 5-min incubation for each wash.

134|Stain nuclei with DAPI (2 µg/mL in DPBS) for 5 minutes then rinse cells two more times with DPBS. 135|Capture images using a confocal microscope equipped with a 63X or 100X objective (**Fig. 2b**).

**? TROUBLESHOOTING**

### Measuring changes in specific protein’s translation efficiency in response to controlled, localized REM buildup by Localis-REX**–**GCE

▴ **CRITICAL**

This section and the next measure the consequences of localized REM buildup on protein synthesis. GCE reports on a single nascent reporter protein, and BONCAT reports on the nascent proteome. Both build on the live-cell Localis-REX steps (Steps 96-118).

#### Cell culture, preparation, and treatment – • TIMING 3 d (hands-on duration: approx. 1 h)

136|Perform cell culture, transient transfection and Localis-REX in live cells as described above (Steps 96-106). When aminoacyl-tRNA synthetase MmPyIRS and synthetic actin is co-translated with Halo plasmids: use (pCS2+8 NLS-Halo, pCS2+8 NES-Halo pCS2+8 MOMLS-Halo, or pCS2+8 ERt-Halo); Actin (K118TAG); and MmPylRS aminoacyl-tRNA synthetase (plasmid ratio is 1:1:0.2) with Lipofectamine 3000 reagent per the manufacturer’s recommendation.

**! CRITICAL STEP**

The ratio of plasmid amount of Actin (K118TAG) and MmPylRS aminoacyl-tRNA synthetase is critical. Optimization is required if changed to other aminoacyl-tRNA synthetase/ UAA pairs.

**? TROUBLESHOOTING**

#### GCE based translation report – • TIMING 8 h

137|After the uncaging step is complete (Step 105), incubate cells with rinsing medium with 60 µM BCNK for 4 hours.

138|Wash with DPBS and harvest cells. Wash twice with cold DPBS and freeze the cells in liquid nitrogen.

**! CRITICAL STEP**

To avoid degradation of BCNK, process the cell lysate on the same day or no later than the next day after cell harvest.

139|Perform cell lysis (Steps 110-117) to obtain 1 mg/mL cell lysate.

140|Incubate lysate with Tetrazine-Cy5 (1 µM) for 30 minutes at 37 °C with gentle shaking.

141|Add 4× Laemmli dye containing 6% βME to samples and incubate for an additional 10 min at 37 °C. Samples are now ready for SDS-PAGE and western blot analysis, typically with anti-HA antibody.

### *Measuring nascent proteome synthesis efficiency changes in* response to localized buildup of reactive metabolites by Localis-REX–GCE or Localis-REX–BONCAT

**! CRITICAL**

Two orthogonal labelling chemistries are given for the same question. Either may be used; agreement between them provides a useful cross-check.

#### Proteome-wide ncAA incorporation by Localis-REX–GCE

142|Perform cell culture, Localis-REX and GCE based translation report in live cells as described (Steps 136-141), except the transfection step. Co-transfect the plasmid encoding Val-MmPylRS aminoacyl-tRNA synthetase with a Halo plasmid (pCS2+8 NLS-Halo, pCS2+8 NES-Halo pCS2+8 MOMLS-Halo, or pCS2+8 ERt-Halo) at a 1:1 ratio. Transfect with Lipofectamine 3000 reagent per the manufacturer’s recommendation.

#### BONCAT: Cell culture, preparation, and treatment – • TIMING 2 d (hands-on duration: approx. 6 h)

143|Perform cell culture and transfection as described in *Localis-REX in mammalian cells (Steps 96-98)*.

**? TROUBLESHOOTING**

#### BONCAT: labelling, lysis and Click chemistry – • TIMING 12 h

144|36 h post transfection, aspirate medium and incubate cells with rinsing medium (MEM with 1X NEAA, 1X Pyruvate and 1X Pen/Strep) containing 10 µM Ht-PreHNE(no alkyne) for 2 hours.

145|Rinse cells two times with fresh rinsing medium, incubate cells for 30 min during each rise to fully remove unbound probe.

146|Change medium to BONCAT rinsing medium (DMEM without methionine) and incubate for 30 min. 147|Under Localis-REX condition, expose cells to 365 nm light (5 mW/cm^2^) for 5 min; under light-alone control, treat cells with DMSO instead of Ht-PreHNE(no alkyne) and expose to 365 nm light for 5 min; under no light condition, treat cells with 10 µM Ht-PreHNE(no alkyne) for 2 h, but do not irradiate; under DMSO control, treat cells with DMSO instead of Ht-PreHNE(no alkyne) and no light irradiation.

**! CAUTION**

Wear UV-protective goggles, should the lamp be in operation with an unshielded beam path.

148|Change medium to BONCAT rinsing medium, add L-Azido-homoalanine (final concentration 50 µM), and incubate for 4h.

149|Harvest cells from all conditions and wash cells twice with cold DPBS. Snap-freeze cells in liquid nitrogen for downstream analysis.

**▪ PAUSE POINT**

Samples can be stored at −80 °C for up to 1 week without significant loss of signal.

150|Perform cell lysis (Steps 110-117) described before to obtain 1 mg/mL cell lysate.

151|Perform the Click chemistry steps described before. I. e., incubate the lysate with 1% SDS, 5% t-BuOH, 1 mM CuSO_4_, 0.1 mM Cu(II)TBTA, 2 mM TCEP, and 10 μM Cy5-alkyne for 30 minutes at 37 °C with gentle shaking.

152|Add 4× Laemmli dye containing 6% βME to samples and incubate for an additional 10 min at 37 °C. Samples are now ready for gel loading, in-gel fluorescence after SDS-PAGE and western blot analysis, typically with anti-HA antibody.

**▪ PAUSE POINT**

Samples in Laemmli dye can be stored at −20 °C overnight.

##### Box 3: ultraID mapping: abundance-based organ-specific protein profiling in *C. elegans*

**CRITICAL** For optimal ultraID mapping, we recommend including two control groups in the experimental setup (**Fig. 3a**).

**General culture procedures for OS-ultraID strains –** ▴ **TIMING 5-7 d** (hands-on duration – picking per plate: approx. 15 min; + 2 h)

▴ **CRITICAL**

Steps 1-5 here are essential for obtaining enough worms on MG1655:BioBKan food. Biotin deficiency limits the number of F2 progeny that can be raised from 20 P0 adults on MG1655:BioBKan plates. Maintain all experimental and control groups in the same MG1655:BioBKan feeding background and handle them in parallel.

1| Pick 20 healthy OS-ultraID transgenic gravid adult worms (P0) to growth plates with OP50 using platinum picks (**Fig. 3b**). Culture at 20 °C for 5-7 days to reach a confluent state with F2 progeny (densely populated but not starved).

▴ **CRITICAL STEP**

The F2 progeny are usually at the L1-L2 stage at the collection point. Check the plates every day starting from day 4 to prevent starvation.

2| Add 3 mL S buffer to each 10 cm plate. Use a glass pipette to gently wash worms off the plates and collect the suspension into a 15 mL Falcon tube.
3| Centrifuge the tube at 3000 × g for 1 minute to pellet the worms. Carefully aspirate the supernatant without disturbing the pellet.
4| Wash the worm pellets with 10 mL S buffer each time, centrifuging at 3000 × g for 1 minute between washes. Repeat until the supernatant appears clear.
5| Resuspend the washed worm pellets in S buffer. Adjust the resuspension volume according to the number of destination plates. For example, distribute the worms from one original plate across six freshly prepared 10 cm MG1655:BioBKan plates, adding approximately 0.5 mL of resuspended worms per plate. (i.e., 1 plate to 6 plates).

### Optimization of External Biotin Treatment for OS-ultraID in C. elegans – • TIMING 3 d (hands-on duration: approx. 10 h)

6| Following step 3.5, wait 2 more days until the MG1655:BioBKan plates reach a confluent state. One 10 cm plate is typically used for each condition.
7| Add 3 mL S buffer to each 10 cm plate. Use a glass pipette to gently wash worms off the plates and collect the suspension into a 15 mL Falcon tube.
8| Centrifuge the tube at 3000 × g for 1 minute to pellet the worms. Carefully aspirate the supernatant without disturbing the pellet.
9| Wash the worm pellets with 10 mL S buffer each time, centrifuging at 3000 × g for 1 minute between washes. Repeat until the supernatant appears clear.
10| Resuspend the worm pellets in 1 mL S medium. Pool the suspensions and then split them equally into 1.5 mL Eppendorf tubes.

▴ **CRITICAL STEP**

The buffer is changed to S medium for the incubation step.

11| Prepare a 4× OP50 food source by centrifuging 10 mL of a saturated OP50 culture and resuspending the bacterial pellet in 2.5 mL of S medium. Add 250 μL of this 4× mixture to each tube of pelleted worms.
12| Prepare 4× biotin stock solutions in S medium (0, 0.8, 4, 20 and nominally 100 mM). For high-concentration biotin stocks (20 mM and 100 mM), add 1 M KOH dropwise to aid dissolution while stirring, and adjust the pH to ∼6.0 during preparation. Add the stock solutions to the worm samples to achieve final biotin concentrations of 0, 0.2, 1, 5, and nominally 25 mM. Adjust the final volume in each tube to 1.0 mL with S medium.
13| Incubate the tubes for 0.5, 1, 2, and 4 hours on an end-to-end rotator at room temperature and under dim red light.

▴ **CRITICAL STEP**

Using streptavidin blot analysis (**Fig. 8**), the two parameters are optimised sequentially, not as a factorial matrix: titrate biotin concentration first, at a fixed labelling time of 1 hour, and then using the selected concentration, titrate the labelling time. This requires 4 conditions + 1 control in total rather than 16.

**Figure 8.**
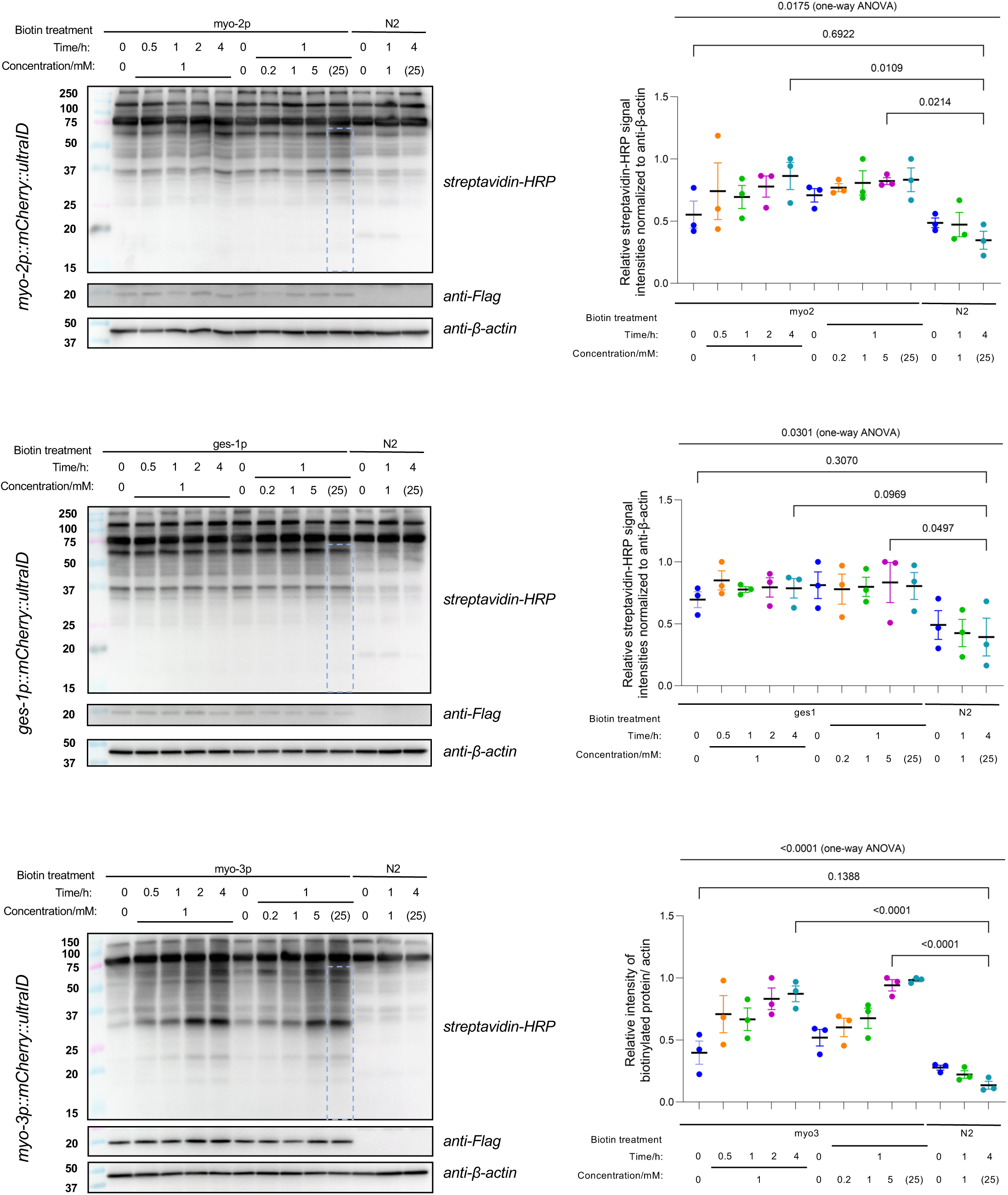
Optimization of external biotin treatment conditions for OS-ultraID. Optimal external biotin treatment conditions were assessed by streptavidin blots. For each OS-ultraID strain, biotin treatments were performed across a time gradient (0 – 4 hours) at 1 mM, or a concentration gradient (0 – 25 mM) for 1 hour. Quantification (**right**) was based on signals below 60 kDa MW marker in corresponding blots (**left**, dotted blue rectangles). Due to the solubility limits of biotin, the 25 mM concentration was not achievable. *P* values were calculated by Dunnett’s multiple comparisons test, following one-way ANOVA. *n =* 3 for all conditions. All data present mean ± SEM. This figure is reproduced from previously published data in reference^11^.

14| After incubation, wash samples twice with 1 mL S buffer and a third time with 50 mM HEPES (pH 7.6). Each wash should be 30 minutes with gentle rotation. Centrifuge at 3000 × g for 1 minute between each wash. Discard as much of the supernatant as possible. Flash-freeze in liquid nitrogen and store the samples at −80 °C. Lyse the worm pellets (see section ‘worm lysis’, Steps 41-49). Adjust lysate concentration to 1.25 mg/mL and add 4× Laemmli buffer with 6% BME. Samples are now ready for gel loading and streptavidin blot analysis (**Fig. 8**).

### Biotinylated protein enrichment for OS-ultraID strains – • TIMING 3 d (hands-on duration: approx. 10 + 8 h)

15| Following step 3.5, wait 2 more days until the MG1655:BioBKan plates reach a confluent state. 6 x 10 cm plate is typically used for each mass-spec sample.
16| Following the harvesting and washing procedure described above (Steps 7-10 in Box 3), treat the worms with 5 mM external biotin (prepared from 4x stock described in step 3.12) for 4 hours. Add buffer instead of biotin stocks for the control group.
17| After worm lysis and protein concentration normalization (steps 41-49), remove a 30 μL aliquot from the lysate. Add 10 μL of 4× Laemmli dye to this aliquot to serve as the input sample for the streptavidin blot analysis.
18| Perform streptavidin-based enrichment by pre-rinsing 50 μL bed volume of streptavidin beads, then proceed with the subsequent enrichment steps as described above until elution samples are collected (Steps 81-91). The pre-clearance steps and the Click chemistry steps from Localis-REX are omitted. For mass-spectrometry analysis of OS-ultraID samples, proceed to Step 153 in the main Procedure.

### Mass spectrometry sample preparation in Localis-REX experiments

**CRITICAL** Mass spectrometry sample preparation and analysis can also be applied to OS-ultraID samples (see Box 3).

#### Trypsin in-gel digestion and desalting for mass-spec samples – • TIMING 1 d (hands-on duration: approx. 4 h)

153|Load 30 μL elution samples or 20 µg of SILAC cell lysate in Laemmli dye into a homemade 11% SDS-PAGE gel or Novex™ Tris-Glycine mini protein gel. Run the gel until the dye-front has penetrated 1-2 cm into the gel to desalt the samples. After running, cut the resolving gel containing protein into pieces (1-2 mm^3^) and collect them into a 1.5 mL Eppendorf tube.

▴ **CRITICAL STEP**

For homemade SDS-PAGE gels, all gel apparatus should be thoroughly cleaned and bleached to minimize potential protein contamination prior to mass spectrometry analysis.

▴ **CRITICAL STEP**

Estimated wet gel volume after cutting will be ∼50-150 μL for 1-2 cm running length.

154|Wash the gel pieces twice with 300 μL of 50% acetonitrile and 50 mM ammonium bicarbonate for 20 minutes each. Repeat until the gel pieces appear nearly colorless and transparent. Dry the pieces by vacuum centrifugation or 100% acetonitrile.

▴ **CRITICAL STEP**

50% ethanol can replace the 50% acetonitrile, but vacuum centrifugation is necessary.

155|Add 100 μL of 10 mM DTT to cover the gel pieces and incubate for 1 hour at 56°C.

156|Wash the gel pieces once with 300 μL 50 mM ammonium bicarbonate for 10 minutes. Then, add 100 μL 55 mM Iodoacetamide and incubate for 45 minutes at 37°C in the dark.

157|Wash the gel pieces once with 300 μL of 50 mM ammonium bicarbonate for 10 minutes. Dry the pieces by vacuum centrifugation or 100% acetonitrile. Add Mass Spectrometry Grade Trypsin (12.5 ng/μL in 50 mM ammonium bicarbonate and 10 mM CaCl₂) to cover the gel pieces (100 μL) and digest overnight (16 h) at 37°C.

158|Extract peptides from the gel pieces twice by incubating with 200 μL of 50% acetonitrile and 5% formic acid for 20 minutes with shaking, collect the supernatant. Repeat the extraction with a further 200 µL and pool the two supernatants. Dry the pooled extract by vacuum centrifugation, and store the resulting pellets at −20 °C.

159|Reconstitute tryptic digests in 0.5% formic acid (FA) at an estimated concentration of 0.1 μg/μL. 160|For Label Free Quantification (LFQ) or SILAC samples: Desalt the peptides using C18 StageTips according to the manufacturer’s instructions. Fractionate the sample using the Pierce High pH Reversed-Phase Peptide Fractionation Kit according to the manufacturer’s instructions. Dry the resulting fractions by vacuum centrifugation and proceed to Step 163. For TMT samples: dry the peptides by vacuum centrifugation and proceed directly to Step 161.

#### TMT labeling (optional) – • TIMING 5 h

161|For TMT labeling, reconstitute the dried peptides in 8 μL of 100 mM HEPES (pH 8.5). Add 3 μL of TMT solution (20 μg/μL in pure acetonitrile) and incubate at room temperature for 1.5 hours. Quench the reaction by adding hydroxylamine to a final concentration of 0.4% (v/v) and incubating for 15 minutes at room temperature.

▴ **CRITICAL STEP**

For affinity-enrichment experiments associated with Localis-REX and ultraID proteomics mapping workflows, total peptide yields are typically very low. The fixed amount of TMT reagent added per channel (60 μg) therefore represents a substantial molar excess relative to the peptide input.

162|Combine all TMT-labeled samples in a 1:1 ratio. Dry the pooled sample by vacuum centrifugation until it is nearly dry. Fractionate the sample using the Pierce High pH Reversed-Phase Peptide Fractionation Kit according to the manufacturer’s instructions to generate 12 fractions. Dry the fractions by vacuum centrifugation and store at −20°C until LC-MS/MS analysis.

### Mass spectrometry data acquisition and analysis

#### LC-MS/MS data acquisition – • TIMING 2 d (hands-on duration: approx. 2 h)

163|Resuspend the 12 fractions (from Step 162) or desalted peptides (from Step 160, label-free/SILAC) in 10 - 15 µL of 2% acetonitrile and 0.1% formic acid, and inject 5 µL. Perform nano-flow separations and data acquisition using the instrument configuration and parameters described in the Equipment Setup section.

#### Database search – • TIMING 1 - 2 d (hands-on duration: approx. 2 h)

164|Process raw files with either MaxQuant or Proteome Discoverer using the search settings specified in the Equipment Setup section. Use the UniProt reference proteome of the relevant organism supplemented with the transgene construct sequence used in the experiment.

165|Filter the output protein tables (proteinGroups.txt for MaxQuant, or Proteins.txt for Proteome Discoverer) as follows: remove entries flagged as reverse decoy hits, potential contaminants, or identified only by site. For Localis-REX experiments, retain proteins identified with ≥ 2 unique peptides; for ultraID experiments, retain proteins identified with ≥ 1 unique peptide. Remove proteins with zero reporter intensity in the channels of interest.

#### Target filtering and analysis – • TIMING 1 - 2 d (hands-on duration: approx. 4 h)

166|For each biological replicate, calculate the per-protein log₂ fold change (log₂FC) isotope-purity-corrected reporter ion intensities in MaxQuant or the Abundance columns in Proteome Discoverer [e.g., log₂(alkyne/no-alkyne) for Localis-REX datasets].

▴ **CRITICAL STEP**

No sample-loading normalization is applied at this stage, as this assumption does not hold for affinity-enrichment experiments.

167|For each Localis-REX replicate, bin the log₂FC values (bin width = 0.1) and fit a Gaussian distribution by nonlinear regression to obtain the replicate-specific mean (μ) and standard deviation (σ). Set the significance threshold at μ + 2.5σ (∼99.4% one-sided confidence). Define a high confidence hit as a protein exceeding this threshold in all biological replicates (*n* = 3).

▴ **CRITICAL STEP**

Use μ + 2.5σ from the fitted Gaussian distribution as an empirical enrichment threshold instead of a universal statistical rule. Users should inspect the fitted distribution before applying this cutoff, because strongly skewed or multimodal distributions may require additional filtering or alternative statistical assessment.

▴ **CRITICAL STEP**

Proteins passing this threshold constitute the locale-specific responder list. Candidate responders may be carried forward to phenotypic validation by RNAi (Steps 170-182), and to cross-species conservation analysis (see Applications).

168|Volcano plot visualization (optional): Use a two-sample Student’s t-test (equal variance, two-tailed, *n* = 3 per group) to compute per-protein *P*-values for volcano plot display.

▴ **CRITICAL STEP**

For alternative statistical frameworks (e.g., limma or Perseus), see Box 1 in Equipment Setup.

169|Whole-proteome abundance analysis (optional): For whole-proteome LFQ analysis, process raw files with MaxQuant with LFQ enabled. In Perseus, filter for proteins with ≥ 2 valid values in at least one group and impute missing values from a normal distribution (width = 0.3, down-shift = 1.8).

▴ **CRITICAL STEP**

These data can be used to verify that transgene expression does not perturb the host proteome and to cross-reference enrichment hits with total protein abundance.

### Phenotypic screening by RNAi in *C. elegans* and imaging of stained animals

**! CRITICAL**

These steps are the functional validations of the protocol and are performed after target identification in *C. elegans*. Localis-REX together with mass-spec (Steps 52-95; Steps 152-168) returns a ranked list of locale-specific electrophile responders. The steps below knock down individual candidates by RNAi feeding so that their contribution to organism-level phenotypes can be scored. They are not required for target identification itself.

#### RNAi plates preparation and RNAi – • TIMING 3 d (hands-on duration: approx. 3 h)

170|When preparing the worm growth plates, beyond adding cholesterol, add IPTG (from a 1 M stock) to a final concentration of 1 mM and ampicillin to a final concentration of 50 μg/mL. Pour 15 mL of media per 10 cm Petri dish under sterile conditions.

171|Transform the relevant clones from the Ahringer RNAi feeding library (or L4440 empty vector as control) into *E. coli* HT115(DE3) competent cells. Grow cultures overnight in LB containing 50 μg/mL ampicillin at 37 °C.

▴ **CRITICAL STEP**

Users need to select the clones corresponding to their own Localis-REX candidates. The clones used in this protocol are listed in the Reagents section.

172|Concentrate the bacterial culture by 5x and seed 1.5 mL onto IPTG-containing NGM plates. Allow the plates to dry and induce dsRNA expression overnight at room temperature.

173|Place synchronized L1 worms onto the RNAi plates. Incubate at 22 °C for 40–48 hours until worms reach late L4 stage.

**? TROUBLESHOOTING**

#### TMR-Halo staining – • TIMING 6 h

174|Collect worms into 1.5 mL Eppendorf tubes, wash twice with M9 buffer.

175|Add TMR-Halo ligand to a final concentration of 5 μM. Incubated the tubes at room temperature (25 °C) for 3 hours with end-to-end rotation, protected from light (under dim red light or dark).

176|Remove supernatant and wash worms three times with 500 μL M9 buffer, 30 min each. Resuspend stained worms in 250 μL M9 buffer.

177|Distribute the stained worms on NGM plates with OP50. Worms can be imaged on plates or on imaging glasses. To align the worms and generate detailed pictures, pick the worms under the mCherry/RFP channel and mount worms on 1% agarose drop with levamisole. Image the worms using Leica M205 FA microscope under stereo and 5× HR objectives.

▴ **CRITICAL STEP**

Do not pick worms showing strong red signals inside the intestinal lumen as they are considered unhealthy and unable to expel unbound TMR-Halo ligand. If strong unspecific binding is observed in the posterior intestine, the stained worms can be imaged the next day.

#### Oil Red O staining and imaging – • TIMING 3 d (hands-on duration: 2-4 h)

178|Collect worms into 1.5 mL Eppendorf tubes, wash 3 times with M9 buffer. Fix worms in 500 μL 60% isopropanol.

**! CRITICAL STEP**

Worms should be synchronized, as different development stages and ages will change the lipid amount in the body. Worms are killed by isopropanol at this step.

179|Add 500 μL freshly 0.22-µm filtered 0.3% Oil Red O in 60% isopropanol. Stain worms for 18 hours at 25 °C with end-to-end rotation. Keep the tubes protected from light (e.g., covered in aluminum foil) and sealed with Parafilm to prevent any solvent evaporation during the long incubation.

**! CRITICAL STEP**

Ensure the working solution is free of precipitates before application (Check the Oil Red O staining solution preparation), as particulates can adhere to the specimen and interfere with staining results.

180|Remove supernatant and wash worms twice with 500 μL 60% isopropanol. Resuspend stained worms in 250 μL 0.01% Triton X-100 in S buffer.

**▪ PAUSE POINT**

Samples in S buffer can be stored at 4 °C for up to 1 month.

181|Distribute the stained worms on worm growth plates without OP50. Pick the worms randomly and mount worms on 1% agarose drop and image using Leica M205 FA microscope under stereo and 5× HR objectives.

182|For Oil Red O quantification, acquire images in RGB mode under 5× HR objective. Analyze blue channel after inverting images, selecting the worm body (excluding pharynx).

#### Phenotypic screening – • TIMING 7 - 25 d

183|Perform phenotypic screening. Follow option A to score the number of fertilized eggs laid; option B to score egg laying following acute HNE(alkyne) challenge; option C to assay viability; option D to assay locomotion by counting body bends; option E to assay locomotion towards food in a food race; and option F to record the defecation motor program. Options A, C and D report the consequences of RNAi knockdown alone; options B and E additionally apply an acute HNE(alkyne) stress, and option F is performed both with and without stress.

**! CRITICAL**

For all phenotypic screening, the OP50 or HT115 *E. coli* density before seeding onto the plates is recommended to be normalized to an OD_600_ of 1-3.

A. Scoring of fertilized eggs laid by *C. elegans **–*** • **TIMING** (hands-on duration: 10-30 min per plate per day)

1. At 40 h post-L1 plating, select 12 L4-young adult hermaphrodites from RNAi-treated 10 cm plates and transfer each worm to an individual 6 cm RNAi plate.
2. Transfer the worms to fresh RNAi plates twice daily (morning and evening) on Days 1 and 2 post-L1 plating. Continue daily evening transfers on subsequent days until no new eggs are laid.
3. Count the number of fertilized eggs at defined intervals: three times per day on Days 1 and 2, twice per day on Days 3 and 4, and once daily thereafter.
4. Confirm that all counted eggs are fertilized by assessing hatching.
B. *C. elegans* egg-laying assay following HNE-alkyne treatment – • **TIMING** *(hands-on duration: 10-30 min per plate per day)*

1. At 40 h post-L1 plating, harvest worms, wash three times with M9 buffer to remove bacteria, and resuspend in 0.5 mL M9 per sample.
2. Add HNE-alkyne solution (final 2 mM in 0.75 mL) or DMSO control. Incubate with rotation for 1 h. Wash worms 3 times with 0.5 mL M9 and resuspend in 0.2 mL M9.
3. Seed worms onto fresh RNAi plates and allow them to recover for 4 h at 20 °C. Select 12 L4-young adult worms and transfer each to a separate 6 cm RNAi plate.
4. Transfer worms and count eggs using the same schedule described in step A.
C. *C. elegans* viability assay – • **TIMING** (hands-on duration: approx. 10 min per plate per day)

1. At 40 h post-L1 plating, pick 10 L4-young adult worms from RNAi-treated 10 cm plates and transfer individually to 6 cm RNAi plates.
2. Monitor worm viability daily. A worm is scored as dead if it fails to respond to gentle head-touching with a platinum wire and shows no pharyngeal pumping.
3. Transfer worms to fresh RNAi plates twice on Days 1–2, then once daily in the evenings thereafter, until egg-laying ceases. Thereafter, score the animals in place, without further transfer, until death.
D. *C. elegans* body bend assay – • **TIMING** (hands-on duration: approx. 20 min per plate per day)

1. At 40 h post-L1 plating, pick 10 L4-young adult worms from RNAi-treated 10 cm plates and transfer individually to 6 cm RNAi plates.
2. Every other day, count the number of body bends for 2 minutes per worm. Record data only when worms are clearly away from the edge of the bacterial lawn.
3. Continue monitoring until each worm dies. Transfer worms to fresh RNAi plates following the same schedule as the viability assay.
E. *C. elegans* food race assay – • **TIMING** (hands-on duration: approx. 10 min per plate per day)

1. Prepare food race plates by spotting 50 μL OP50 1 cm from the plate edge on the equator line. Let dry for at least 2 days.
2. At 40 h post-L1 plating, harvest worms, wash three times with M9 buffer to remove bacteria, and resuspend in 0.5 mL M9 per sample.
3. Add HNE-alkyne solution (final 2 mM in 0.75 mL) or DMSO control. Incubate with rotation for 1 h. Wash worms 3 times with 0.5 mL M9 and resuspend in 0.2 mL M9.
4. Transfer worms to the opposite side of the OP50 dot (∼1 cm from edge) and start timing once M9 evaporates.
5. Count the number of worms that reach the bacterial lawn at 30, 60, 90, and 120 min, and again the next day. Normalize to the total number of viable worms.
F. *C. elegans* defecation assay – • **TIMING** (hands-on duration: approx. 10 min per plate per day)

**! CRITICAL**

The defecation motor program is an rhythm with a period of approximately 50 s in well-fed wild-type animals at 20 °C^89^. Each cycle comprises three stereotyped muscle contractions in a fixed order: a posterior body-wall muscle contraction (pBoc), in which the posterior body wall constricts and intestinal contents are driven forwards; an anterior body-wall muscle contraction (aBoc) approximately 1 s later, which drives the contents back towards the rectum; and an expulsion (Exp), in which the enteric muscles contract and the contents are expelled. The cycle is timed from the onset of one pBoc to the onset of the next, because pBoc is the most sharply defined of the three events and the easiest to score under a stereomicroscope.

1. Prepare RNAi-fed worms on 10 cm plates (15 mL NGM with OP50 normalized to OD600 = 1.2)
2. Raise synchronized L1 worms (Steps 9-15) for 48 h at 22 °C. For defecation dataset of late L4/young adult transition stage worms, record 10 defecation cycles per worm (12 worms per condition) using Leica M205 FA microscope.

**! CRITICAL STEP**

Score only animals that remain on the bacterial lawn and continue feeding throughout the recording. Discard the record from any animal that leaves the lawn and score a replacement.

III. For the HNE-treatment dataset, for each biological replicate and RNAi condition, split worms from the same synchronized population into two matched groups. Keep one group as the untreated control and treat the matched second group with HNE-alkyne (2 mM final in 0.75 mL, 1 h rotation). Wash worms 3 times with 0.5 mL M9 and resuspend in 0.2 mL M9.
IV. Seed worms onto fresh 10 cm RNAi plates and incubate for 16 h at 20 °C.
V. Record 6 defecation cycles per worm (20 worms per condition). Define defecation interval based on pBoc contraction cycles.

##### Box 4: Statistical analysis

**CRITICAL** For *C. elegans* experiments, samples generated from the same pool of animals (e.g., 6 × 10 cm plates, depending on the experiment) are considered one biological replicate. For cultured cell experiments, samples from individual wells or plates are considered biological replicates.

**CRITICAL** *P* values should calculated using t-tests or ANOVA with adjusted comparisons, as explained in the following decision logic for selecting the appropriate test (**Extended Fig. 2**):

1| Organize all collected data points according to experimental design and biological replicates. Group data into experimentally relevant comparisons.
2| Determine data structure. If data points come from a hierarchical or nested design (e.g., multiple measurements taken from the same individual animal), perform a nested t-test to compare two experimental conditions, or a nested ANOVA to compare three or more conditions.

**! CRITICAL STEP**

Nested or hierarchical designs are common in phenotypic recording of individual animals. For example, repeated measurements of defecation intervals nested within individual animals, which are in turn nested within a single experimental condition, should be considered a hierarchical structure and a nested design (**Extended Fig. 3**).

3| If data are collected from different animals or independent samples, perform an unpaired two-tailed Student’s t-test to compare two experimental conditions, or ANOVA to compare three or more conditions. For detailed ANOVA test, refer to the following steps.

▴ **CRITICAL STEP**

Bonferroni correction may also be applied when multiple comparisons increase the risk of Type I errors.

4| For analysis involving one factor, perform one-way ANOVA. Use Tukey’s multiple comparisons test to compare multiple experimental groups to each other or use Dunnett’s multiple comparisons test to compare multiple groups to a single group. This branch is applicable to most single-factor affected phenotypic experiments. E.g., RNAi-affected fertility; the lipid level of multiple small-molecule-treated animals.
5| If analysis involves two factors, perform two-way ANOVA. Use Tukey’s multiple comparisons test to compare multiple experimental groups across factors. When comparing groups restricted to a single factor, use Šídák’s multiple comparisons test for two groups, or refer to (**Box 4**, Step 4) to determine whether to use Tukey’s or Dunnett’s multiple comparisons to compare more than two groups. This branch is applicable to two-factor affected phenotypic experiments. E. g., fertility under gene knockdown (RNAi) and small-molecule treatment; the lipid level of small-molecule-treated animals with different tissue-specific gene knockdown (**Extended Fig. 4**).

▴ **CRITICAL STEP**

Bonferroni correction may also be applied when multiple comparisons increase the risk of Type I errors.

## TROUBLESHOOTING

Troubleshooting advice can be found in Table 1.

**TABLE 1:** Troubleshooting table.

| Step(s) | Problem | Possible reasons | Solution |
| --- | --- | --- | --- |
| 6, 8 | Fungal or bacterial contamination on worm plates | Non-sterile technique during media pouring or worm picking. | Always work in a sterile biosafety cabinet or near the flame. Discard contaminated plates immediately. |
|  | Abnormally thick or dark bacterial lawn | Contamination during OP50 culture or in OP50 stock. | Use autoclaved tips to scratch OP50 stock and change OP50 stocks. Avoid touching the tip with gloved hands. |
| 6 | OP50 bacterial lawn fails to remain centered, flows continuously, and contacts the plate edges | Uneven agar surface during pouring or over-drying of the agar on one side before seeding | Ensure the work surface is level and maintain an appropriate distance from the laminar flow hood air stream. Reduce the volume of OP50 culture. |
| 7, 31 | Worms show low viability from frozen stocks | Poor stock quality, for example, from contamination or improper freezing technique. | Pick the surviving worms onto new plate and clean the progenies by bleaching. Remake stocks. |
| 15 | Low yield of L1 larvae after synchronization | Bleaching duration is too long, killing the eggs | Do not exceed 5 min of incubation, especially when handling multiple samples. Add M9 within 6 min. |
|  |  | Starting worm population is not gravid | Ensure starting plates contain a sufficient number of gravid adults. |
|  | Most of the adult worm bodies are not dissolved | Bleach solution is not fresh and lost activity | Always prepare the alkaline hypochlorite solution immediately before use. |
| 18 | No males | Starting worm population is not L4-young adults or starved | Ensure the worms for heat induction are healthy. Increase the number of worms. |
| 20 | No crossed progeny | Low mating efficiency | Increase the sample sizes. |
|  |  | Males died accidentally or too old | Increase the sample sizes, only pick young and healthy males. |
|  |  | Hermaphrodites are well self-fertilized | Ensure the hermaphrodites for crossing are at L4 stage before self-fertilizing. |
| 47 | Low protein concentration after lysis | Incomplete disruption of the tough worm cuticle | Increase the vortexing time. |
|  |  |  | Increase the lysis buffer volume to at least 4x the worm pellet volume. |
|  |  | Protein degradation during lysis | Add a fresh cOMplete tablet. Do not keep the lysate at room temperature. |
|  | Abnormally high protein concentration in certain samples | Worm debris is collected | Collect the supernatant from top to bottom and slowly. |
| 72,135 | Weak or absent Cy5 signals | The Click reaction was inefficient because TCEP or sodium ascorbate had oxidized | Prepare the TCEP or sodium ascorbate solution immediately before use and add it last. |
|  | High background signals from antibodies | Unspecific antibody binding | Increase the dilution of the antibody. Shorten the incubation time. |
|  |  | Insufficient blocking | Increase blocking time. |
|  | High background signals from Cy5 after Click chemistry reactions | Inadequate washing | Increase the number and duration of wash steps. |
|  |  | Long incubation time | Ensure the incubation time between Click mixture and samples is 30 min. |
| 93,95,118 | Biotin pulldown enriches very little protein, or the input samples have significant protein amount differences | Protein precipitate is not fully resolubilized after ethanol precipitation | Ensure the protein pellet is completely dissolved in the LDS resuspension buffer; sonicate in cycles on ice until no visible pellet remains. |
|  | Significant differences in amount of protein for western blot results in elution samples | Protein precipitate is collected and mixed with streptavidin beads | Ensure the protein pellet is completely dissolved in the LDS resuspension buffer. |
| 98,143 | Low transfection efficiency or high cell death | Poor cell health or suboptimal confluency | Ensure cells are healthy, passaged regularly, and are 70- |
|  |  |  | 80% confluent at the time of transfection. |
| 102 | High detachment after treatment | Washing steps are too harsh | Add and aspirate media gently against the side wall of the dish. |
|  |  | Toxicity from plasmid and PEI | Decrease plasmid concentration or incubation time |
| 136 | High background across the whole lane | Tetrazine-Cy5 reacted non-specifically with the lysate | Run a no-BCNK control in parallel |
| 173 | Positive control RNAi (let-767) fails to produce larval arrest. | RNAi plates were made incorrectly (missing IPTG) | Always add IPTG to RNAi media to induce dsRNA expression. |
|  |  | HT115 bacteria lost the RNAi plasmid | Re-transform the plasmid into a fresh stock of HT115(DE3) cells. Check for ampicillin resistance to ensure the plasmid is present. |

## Timing

Steps 1-15, *C. elegans* culture and preparation: 11 d; hands-on duration: approx. 3-4 h

Steps 16-31, *C. elegans* male induction, crossing, and selection: 30 d; hands-on duration: approx. 8-10 h

Steps 32-51, Optimization of Ht-PreHNE(alkyne) treatment conditions (TMR-blocking) for OS-Localis-REX in *C. elegans*, starting from worm collection: 6 d; hands-on duration: approx. 14 h

Steps 52-95, OS-Localis-REX *in C. elegans*, starting from worm collection: 6 d; hands-on duration: approx. 12-18 h, depending on whether imaging, gel validation, pulldown, or MS preparation is performed

Steps 96-102, mammalian cells culture and preparation: 2 d; hands-on duration: approx. 1-2 h

Steps 103-118, Localis-REX in mammalian cells, starting from cell treatment: 4 d; hands-on duration: approx. 8-12 h

Steps 119-135, Localis-REX validation in mammalian cells (immunofluorescence for probe localization), starting from cell treatment: 2 d; hands-on duration: approx. 8 h, depending on downstream validation workflow

Steps 136-152, Translation-change readouts for Localis-REX validation in mammalian cells (GCE and BONCAT), starting from cell treatment: 4 d; hands-on duration: approx. 12 h

Box 3, Steps 1-19, OS-ultraID mapping in *C. elegans*: 12 d; hands-on duration: approx. 12-18 h Step 153-162, Mass spectrometry sample preparation, 2 d; hands-on duration: approx. 6-8 h

Step 163-169, Mass spectrometry data acquisition and data analysis, 4-6 d; hands-on duration: approx. 6-8 h, excluding instrument run time

Step 170-173, RNAi plates preparation and RNAi treatment, 3 d; hands-on duration: approx. 3 h

Step 174-182, downstream analysis for RNAi worms:

i. TMR-Halo staining and imaging, 6 h; hands-on duration: approx. 2-3 h
ii. Oil Red O staining and imaging, 3 d; hands-on duration: approx. 2-4 h

Step 183, A-F, phenotypic measurements:

A. *C. elegans* egg-laying assay, 5 d; hands-on duration: approx. 10-30 min per plate per day
B. *C. elegans* egg-laying assay following HNE-alkyne treatment, 5 d; hands-on duration: approx. 10-30 min per plate per day
C. *C. elegans* viability assay, 25 d; hands-on duration: approx. 10 min per plate per day
D. *C. elegans* body bend assay, 5 d; hands-on duration: approx. 20 min per plate per day
E. *C. elegans* food race assay, 2 d; hands-on duration: approx. 10 min per plate per day
F. *C. elegans* defecation assay, 2 d; hands-on duration: approx. 10 min per worm per day

## ANTICIPATED RESULTS

REX is a unique tool that enables interrogation of target-specific modification by chemically-defined reactive small molecules, as well as profiling of reactive molecule responsive proteins under near-native signaling conditions. Precisely how much REX mirrors “native signaling conditions” is still under debate. This uncertainty stems from an incomplete understanding of the conditions that give rise to innate reactive metabolite (REM) signaling. Nonetheless, REMs, including reactive lipid derived electrophiles (LDEs) and mitochondria-derived (mt-)REMs, can meaningfully rewire signaling pathways^12, 27, 30^. Given that such facts are quite widely accepted, having a tool to interrogate on-target REM-modification by specific REM-chemotype is very important. Obviously, the relative merits of REX-technologies also have to be weighed against alternative methods. To model acute metabolite stress with defined chemotypes over relatively well-defined durations, the main alternative is bolus dosing with excess reactive small molecules. The bolus approach, regardless of concentration or treatment duration used, does not adequately mimic localized reactive metabolite generation. This is because timing, dosage, chemotype, and locale are not able to be controlled by whole-cell/animal-based bulk administration. REX-technologies control essentially all these parameters.

It is unsurprising that we have been able to ascribe a large number of new signaling roles to endogenous REMs using REX^12, 27, 30^. This includes the first example of neofunctionalization by a reactive small molecule (HNE), for which we give an extended protocol here. Gratifyingly, REX-technologies also work particularly well in model organisms, including zebrafish^26, 27, 30^, and *C. elegans*, the subject of this paper. Mirroring our results in cell culture, REX-technologies in *C. elegans* have provided one of the first and most clear examples of reactive electrophile signaling occurring in a tissue-specific manner. We anticipate that profiling experiments using Localis-REX will reveal new responders and, importantly, provide new mechanistic insights. REX hits link directly to druggable proteins, as reactive metabolite-responsive proteins are inherently chemically malleable.

### Localis-REX-final remarks

REX-technologies have been successful because they are compatible with a large number of different methods. We therefore, hope that other laboratories will try to apply these methods as broadly as possible, combining our approaches with their own custom-designed approaches, using these protocols as a guide. Similarly, we anticipate that Localis-REX will be applicable to use in a wide range of cultured cells and organelles, as well as other nematodes and transparent organisms.

### OUTLOOK

A core premise for Localis-REX’s advanced application potential is that cells, or multicellular organisms, do not uniformly sense endogenous or environmental stress cues (e.g., oxidative stress, heat shock). Especially in *C. elegans*, cell-specific mounting of stress defence has been investigated although generality of the approaches across multiple cell-types largely remains to be demonstrated. For instance in *C. elegans*, thermosensory AFD neurons sense heat shock and enhance serotonin release, activating heat shock transcription factor (HSF1)^90, 91^; age-related decline of ER unfolded protein response (UPR^ER^) is rescued by constitutive expression of neuronal-cell-derived XBP-1 that triggers non-autonomous UPR^ER^ in the intestine^92^; and systemic stress resistance is promoted by GPCR signaling within chemosensory ADL neurons activating intestinal mitochondrial UPR (UPR^mt^)^93^. Conversely, cellular stress response is also highly compartmentalized: fasting induces distinct redox signatures that vary not only between tissues (e.g., neurons and body-wall muscle cells) but also across specific organelles^94^. Similarly, autophagic responses to aging and starvation exhibit strong tissue-specific patterns^95^.

Against this backdrop and growing appreciation surrounding tissue-specific stress regulation, how tissue- or subcellular context-specific reactive metabolite (REM) stress build-up is managed by a battery of locale-specific REM-stress responders, remained difficult to be probed with precision. Our early explorations applying subcellular-targeted Localis-REX have discovered cytoplasm-specific REM-responder targets such as CDK9, despites the nucleus being its primary resident, whereby CDK9-hydroxynonenylation promotes its nuclear translocation downregulating global transcription^12^. Such context-specific nuanced signaling mechanisms cannot be easily mapped using state-of-the-art proximity profiling tools indexing locale-specific protein abundance alone. The growing need of spatiofunctional understanding ideally positions Localis-REX to uncover new localized REM-(and related drugs)-actionable protein networks and how they are rewired in disease.

Aberrant REM-stress signaling is implicated in multiple disease contexts. Lipid-derived REMs, particularly 4-HNE, play critical roles in the pathogenesis of neurological^96, 97^ and cardiovascular^98^ diseases. In each of these contexts, HNE accumulation, HNE-responder protein expression, and detoxification capacity likely converge in specific cell types and compartments, to determine whether the outcome is adaptive signaling or irreversible damage. Traditionally, studies investigating these disease contexts have relied heavily on anti-4-HNE antibodies, or general chemoproteomic platforms that profile HNE-responsive proteins across whole-cell or tissue lysates. These approaches are limited by antibody specificity and inability to achieve real-time tissue-specific perturbation and monitoring *in vivo*. Localis-REX is uniquely positioned to bypass these methodological bottlenecks and precisely map spatial and functional complexity in living cells/animals. The above considerations underscore Localis-REX’s potential to be applied as an *in vivo* locale-specific drug screening, not only in terms of mapping locale-specific drug targets but also to discover or monitor specific phenotypic changes as a result of localised drug buildup, as we demonstrate in this protocol. We have indeed already derived drug-like small molecules from REX-based screens with documented efficacy and reduced off-targets in triple-negative breast cancer xenograft mice models^30, 50^.

## MOVIES

**Supplementary Movie 1.** *C. elegans* imaging

**Supplementary Movie 2.** *C. elegans* Localis-REX treatment

**Supplementary Movie 3.** *C. elegans* Localis-REX sample collection

## DATA AVAILABILITY STATEMENT

*C. elegans* strains generated and reported in this protocol are available at *Caenorhabditis* Genetics Center (CGC). Plasmids are available upon request or via Addgene. Raw and processed data underlying experimental figures and tables that were reproduced, adapted, or replotted from supporting primary research articles—those not included in this protocol—are available through the corresponding primary articles, associated Source Data files and public repositories cited therein. Relevant figure legends indicate where figures and datasets were reproduced, adapted, or replotted from previously published data. Supplementary movies are available through Figshare (https://figshare.com/) with the DOI: 10.6084/m9.figshare.33648007. Proteomics raw datasets are available through the ProteomeXchange PRIDE repository under the accession numbers (PRIDE: PXD027730, PXD052114, PXD052493, PXD051483) reported in the corresponding primary articles, and the corresponding processed data are provided within this protocol manuscript in excel file format. Transcriptomics (RNA-seq) raw datasets associated with cell-based Localis-REX-related applications are available through the Gene Expression Omnibus (GEO) under the accession numbers (GSE265951) and the corresponding processed data are reported in the original research publication.

## ACKNOWLEDGMENTS

Miss Yaren Karakoç (Aye lab) for scientific feedback and proof-reading assistance with the worm-based protocols. Dr. Xin Liang (Aye lab) for assistance with videography. Drs. Shivansh Chawla, Daniel Urul, and Souradyuti Ghosh (Aye lab alumni) for initial establishment of *C. elegans*-based REX-technology workflows. Prof. Kelly Liu (Cornell University) for guidance and mentorship on *C. elegans* strain generation. SUNY Biotech for collaborations with *C. elegans* organ-specific ultraID- and Halo-transgenic strain generation.

## RESEARCH SUPPORT

Localis-REX development and applications across distinct cell-based models and biological contexts are supported by Swiss National Science Foundation (SNSF) Project Funding (grant no. 310030_212211) (2023-2027); Academy of Medical Sciences Professorship (National Academy of Medical Sciences UK) and Department for Science, Innovation & Technology UK (DSIT) (grant no. APR11\1013) (2024-2027); and ERC Advanced Grant 101200375 (2025-2030) (to Y.A.). Localis-REX development and applications across distinct worm-based models and biological contexts were supported by SNSF Project Funding (310030_184729) (2019-2023) (to Y.A.). D.C., a former postdoctoral researcher in Y.A.’s laboratory, was supported by the AMS and SNSF during their postdoctoral tenure. Novartis Foundation for Medical-Biological Research, Switzerland (2023-2024) (to M.J.C.L.).

## AUTHOR CONTRIBUTIONS

Data collection: J.L. (Fig. 2a, 3—4, 6—8); D.L. (Fig. 2b); J.L. and M.J.C.L. (Fig. 5). Detailed experimental descriptions for worm-based protocols: J.L. with proof-reading assistance from M.J.C.L. and Y.A.; for cell-based protocols: M.J.C.L. and Y.A. with initial draft from D.C. Data analysis: J.L., M.J.C.L. and Y.A. (worm-based data); D.C., M.J.C.L. and Y.A. (cell-based data). Project supervision and funding acquisition: Y.A. Development of worm-based protocols: J.L., M.J.C.L. and Y.A.; of cell-based protocols: all authors.

## EXTENDED FIGURE LEGENDS

**Extended Figure 1.**
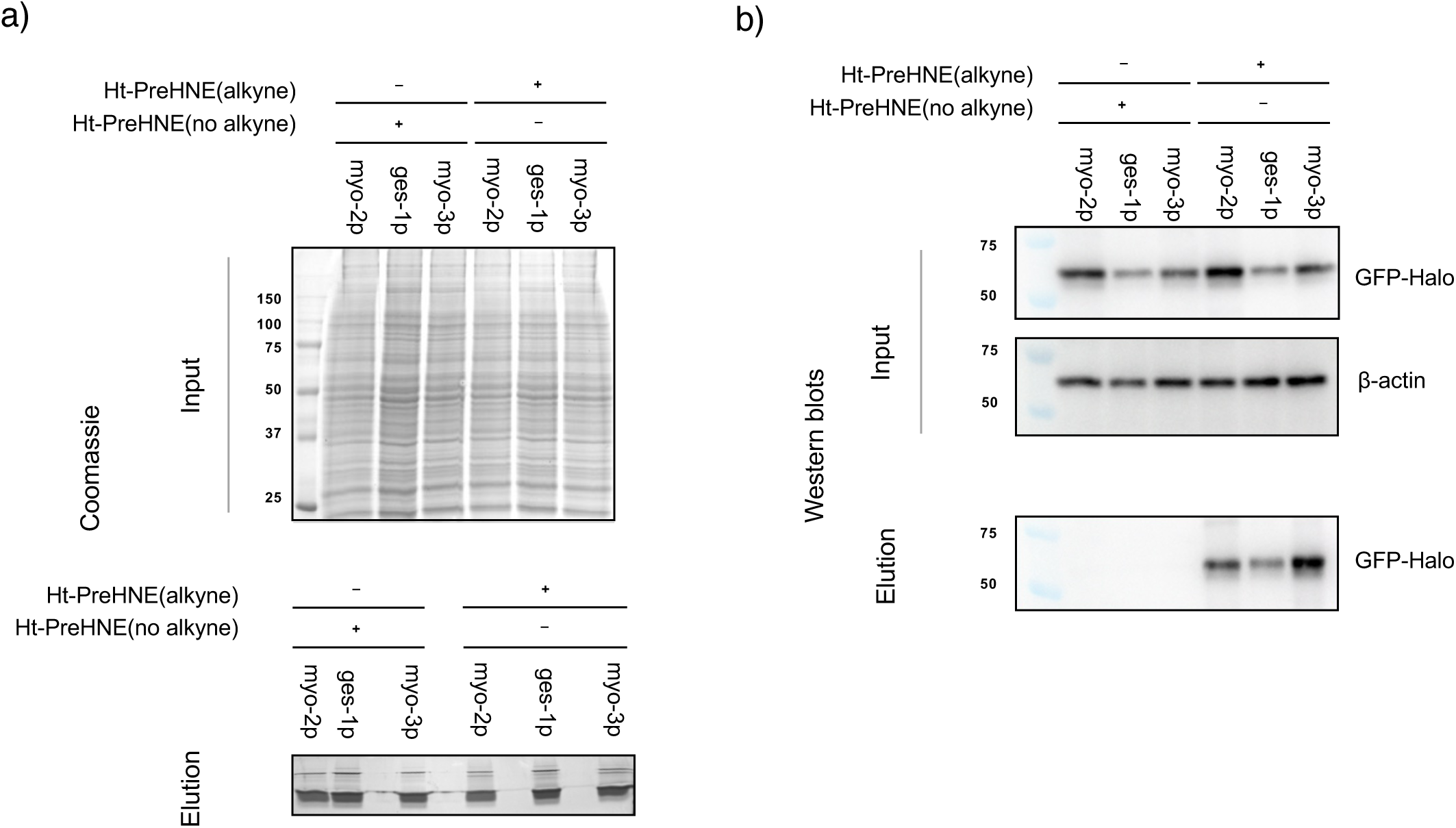
Quality-control gels and blots for Localis-REX mass-spectrometry sample processing. Representative Coomassie blue-stained gels (**a**) and blots (**b**) containing samples for Localis-REX-coupled mass-spectrometry sample processing and data acquisition. Absence of GFP-Halo enrichment in no-alkyne controls confirms specificity in probe-dependent capture. This figure is reproduced from previously published data in reference^11^.

**Extended Figure 2.**
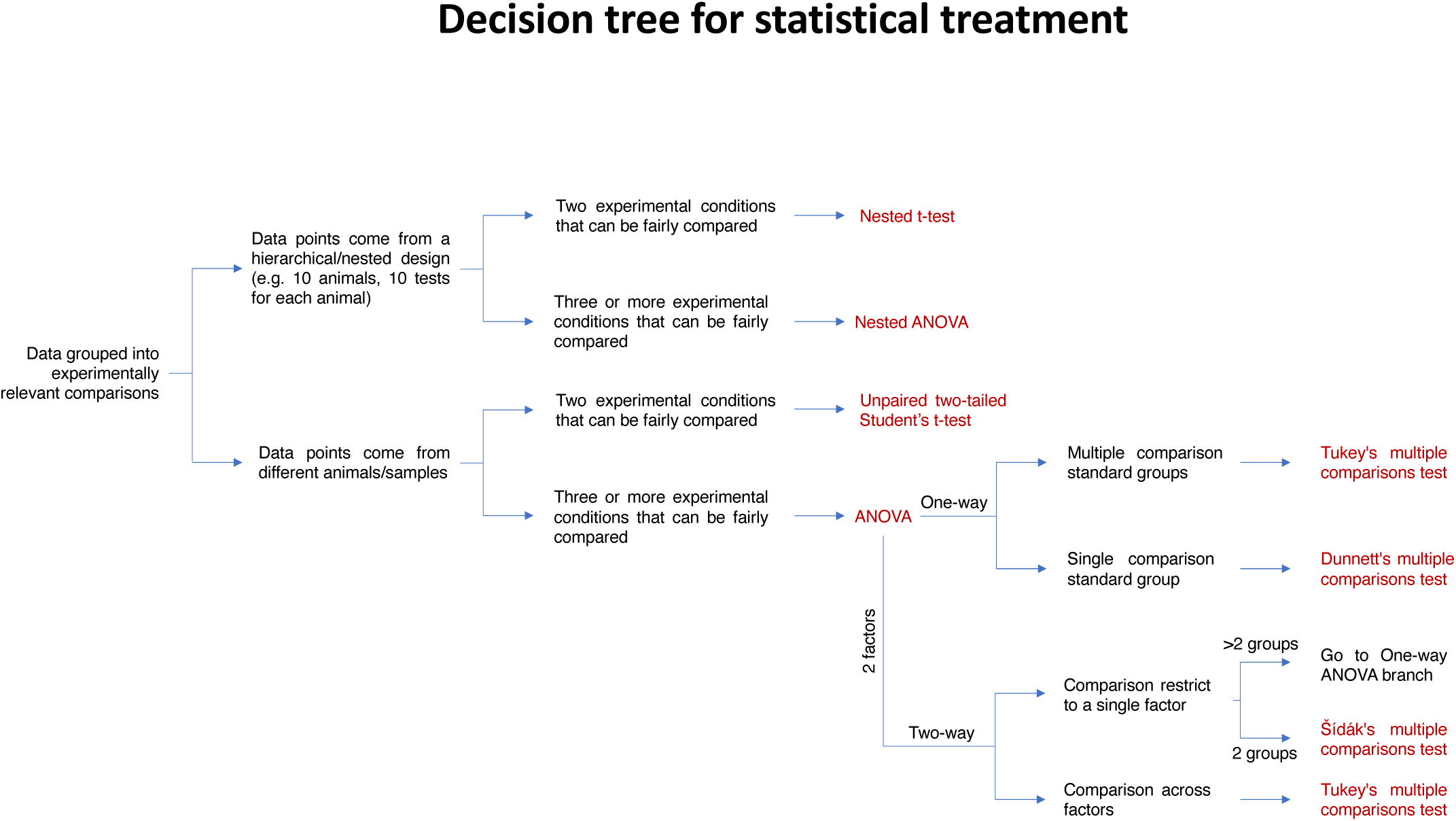
Decision tree for statistical treatment. This figure is reproduced from previously published data in reference^11^.

**Extended Figure 3.**
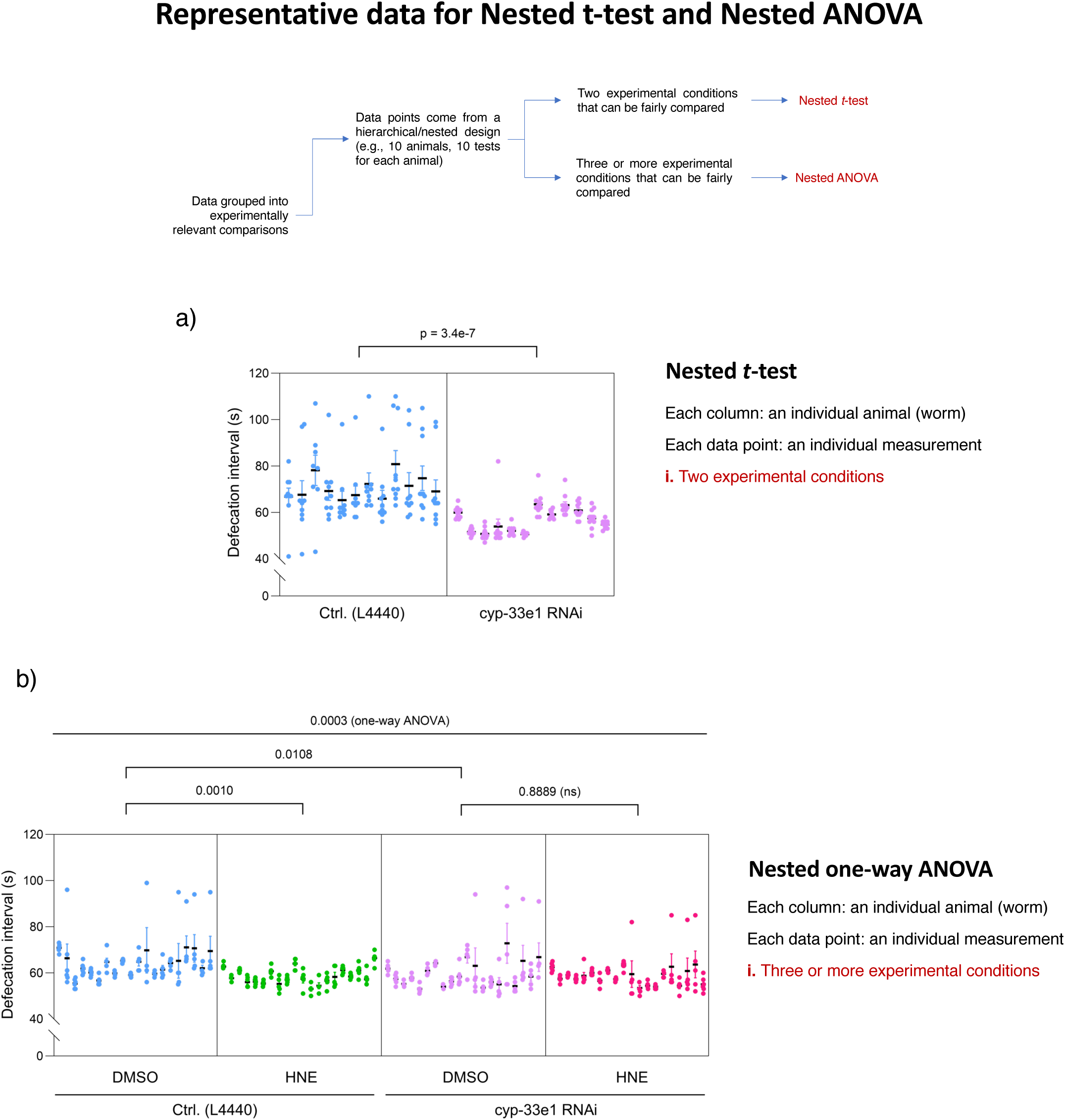
Representative data illustrating nested statistical analyses. Decision tree (top) summarizing criteria for selecting nested *t*-test and nested ANOVA. In both plots, each column represents one animal, and each data point one measurement. Data points come from a nested design: i.e., multiple recording from one signal animal. (a) Representative nested t-test. (b) Representative nested one-way ANOVA with Tukey’s post-hoc test. This figure is adapted from previously published data in reference^11^.

**Extended Figure 4.**
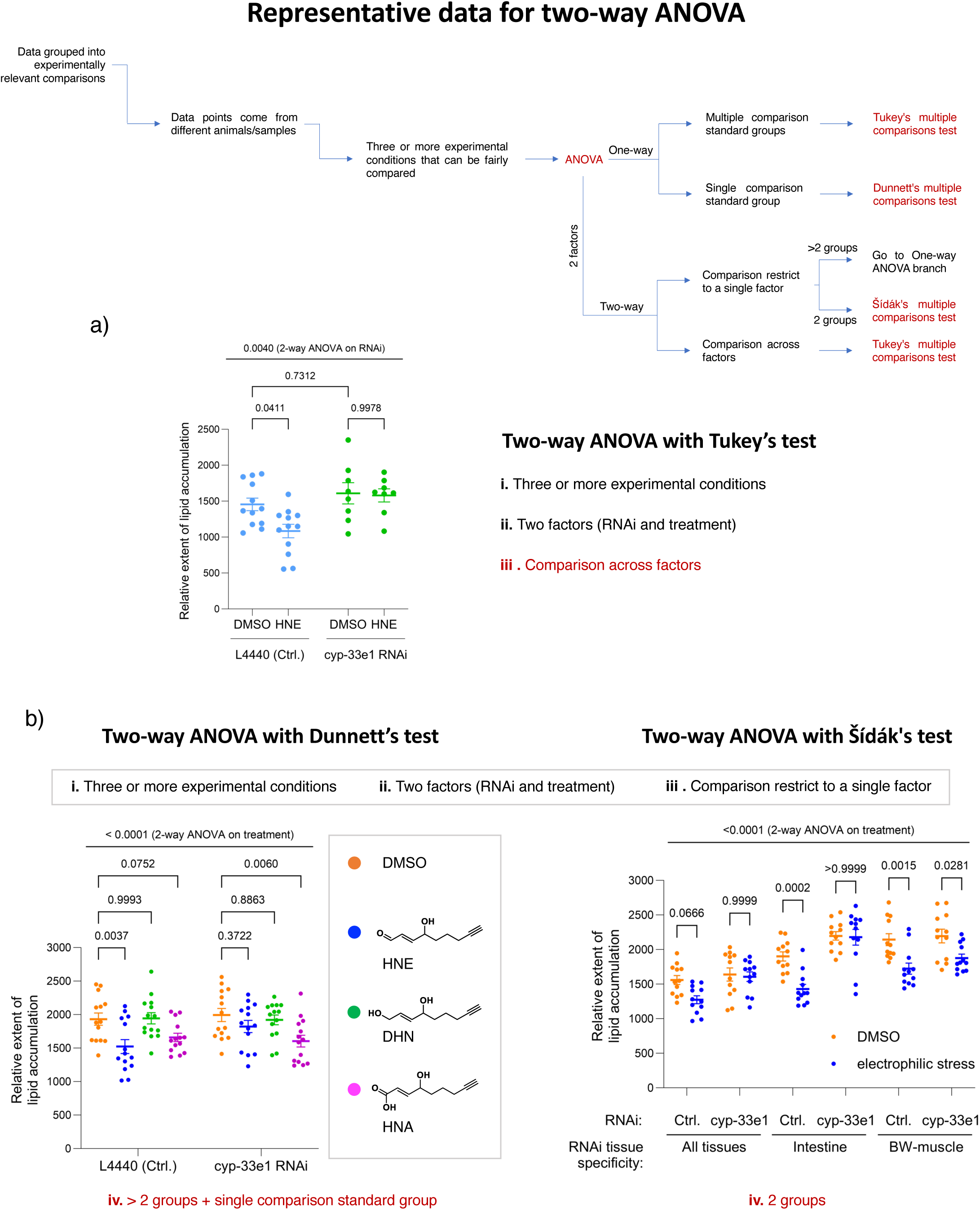
Representative data illustrating two-way ANOVA analyses. Decision tree (top) summarizing criteria for selecting among ANOVA variants. (a) Two-way ANOVA with Tukey’s multiple comparisons test. The multiple comparison test was performed across factors (RNAi and treatment). (b) Left: two-way ANOVA with Dunnett’s multiple comparisons test across three 4 conditions (DMSO, HNE, DHN, HNA). Right: two-way ANOVA with Šídák’s multiple comparisons test comparing DMSO and electrophilic stress across tissue-specific RNAi conditions. This figure is adapted from previously published data in reference^11^.

